# No silver bullet: Patterns of macrosynteny recapitulate systemic conflicts in the higher-level relationships of the arachnids

**DOI:** 10.64898/2026.06.22.733561

**Authors:** Siddharth S. Kulkarni, Benjamin C. Klementz, Jesús A. Ballesteros, Kaitlyn M. Abshire, Tauana J. Cunha, Mohamed K. Hassan, Ethan M. Laumer, Bruno A. S. De Medeiros, Sophie M. Neu, M. Sabrina Pankey, David C. Plachetzki, Carlos E. Santibáñez-López, Emily V.W. Setton, Rebecca M. Varney, Mohamed A. Abdel-Rahman, Gustavo Hormiga, Prashant P. Sharma

**Affiliations:** Department of Integrative Biology, University of Wisconsin-Madison, Madison, WI, USA; CSIR-Center for Cellular and Molecular Biology, Hyderabad, India; Academy of Scientific and Innovative Research (AcSIR), Ghaziabad, India; Department of Biology, Kean University, Union, NJ, USA; Department of Biology, Loyola University Chicago, Chicago, IL, USA; Smithsonian Tropical Research Institute, Panama City, Panama; Zoology Department, Faculty of Science, Port Said University, Port Said, Egypt; Negaunee Integrative Research Center, Field Museum of Natural History, Chicago, IL, USA; Department of Molecular, Cellular and Biomedical Sciences, University of New Hampshire, Durham, NH, USA; Department of Biology, Western Connecticut State University, Danbury, CT, USA; Whitney Laboratory for Marine Bioscience, University of Florida, St. Augustine, FL, USA; School of Biological Sciences, University of Nebraska, Lincoln, NE, USA; Zoology Department, Faculty of Science, Suez Canal University, Ismailia, Egypt; Department of Biological Sciences, The George Washington University, Washington, DC, USA; Zoological Museum, University of Wisconsin-Madison, Madison, WI, USA

**Keywords:** homoplasy, rare genomic changes, polytomy, incongruence, Chelicerata

## Abstract

Rare genomic changes have long been sought by phylogeneticists for their potential to resolve obdurate nodes in the tree of life. Recently, patterns of macrosynteny have been proffered as a breakthrough for challenging relationships within invertebrates. One taxon that stands to benefit from the application of this approach is Chelicerata (the sister group to the rest of Arthropoda), whose radiation has long defied resolution, despite intensive investigations using morphological characters, molecular sequence data, and a combination thereof. Challenges to the resolution of chelicerate phylogeny include an ancient rapid radiation, the incidence of several fast-evolving lineages prone to long-branch attraction artifacts, and extinction of multiple ordinal level lineages that cannot be sampled for breaking long branches. At present, only a subset of nodes has been stably resolved. To break this impasse, we brought to bear multiple classes of phylogenetically informative rare genomic changes, including the sequencing of the first genomes for Ricinulei and Palpigradi. Here, we show that an ancient, shared whole genome duplication event is restricted to Arachnopulmonata (the most recent common ancestor of spiders and scorpions), disfavoring traditional placements of either Ricinulei or Palpigradi as close relatives of tetrapulmonates. Intriguingly, investigation of fusion-with-mixing events identified equal support for mutually exclusive placements for Acariformes, the least stable of the arachnid orders. Our results suggest that fusion-with-mixing, far from being a silver bullet, likely exhibits the same emergent property as all character systems, in that it is prone to homoplasy and conflicting signal stemming from ancient rapid radiations.

## Introduction

Deciphering the ancient radiation of the arachnids is a phylogenetic problem that has long defied resolution, despite intensive investigations using morphological and molecular sequence data (Wheeler and Hayashi 1998; Giribet et al. 2002; Shultz 2007; Sharma, Kaluziak, et al. 2014).

The century-old paradigm of chelicerate evolution held that horseshoe crabs (Xiphosura) comprise the sister group to a monophyletic Arachnida, with groups like the extinct sea scorpions (Eurypterida) and the extant scorpions (Scorpiones) representing progressive stepping-stones between Xiphosura and the remaining arachnids (Scholtz and Kamenz 2006; Shultz 2007; Dunlop 2010). However, phylogenomic studies sampling all extant arachnid orders recover horseshoe crabs as nested within the arachnids, whereas scorpions are recovered as part of a derived group with other book lung-bearing orders (Arachnopulmonata) (Sharma, Kaluziak, et al. 2014; Ballesteros et al. 2019; Ballesteros and Sharma 2019; Ballesteros et al. 2022; Domènech et al. 2026). The latter placement is now well-supported by the discovery of an ancient whole genome duplication shared by the arachnopulmonates, which were later recognized to include pseudoscorpions (Schwager et al. 2017; Ontano et al. 2021) (Fig. 1a).

**Figure 1.**
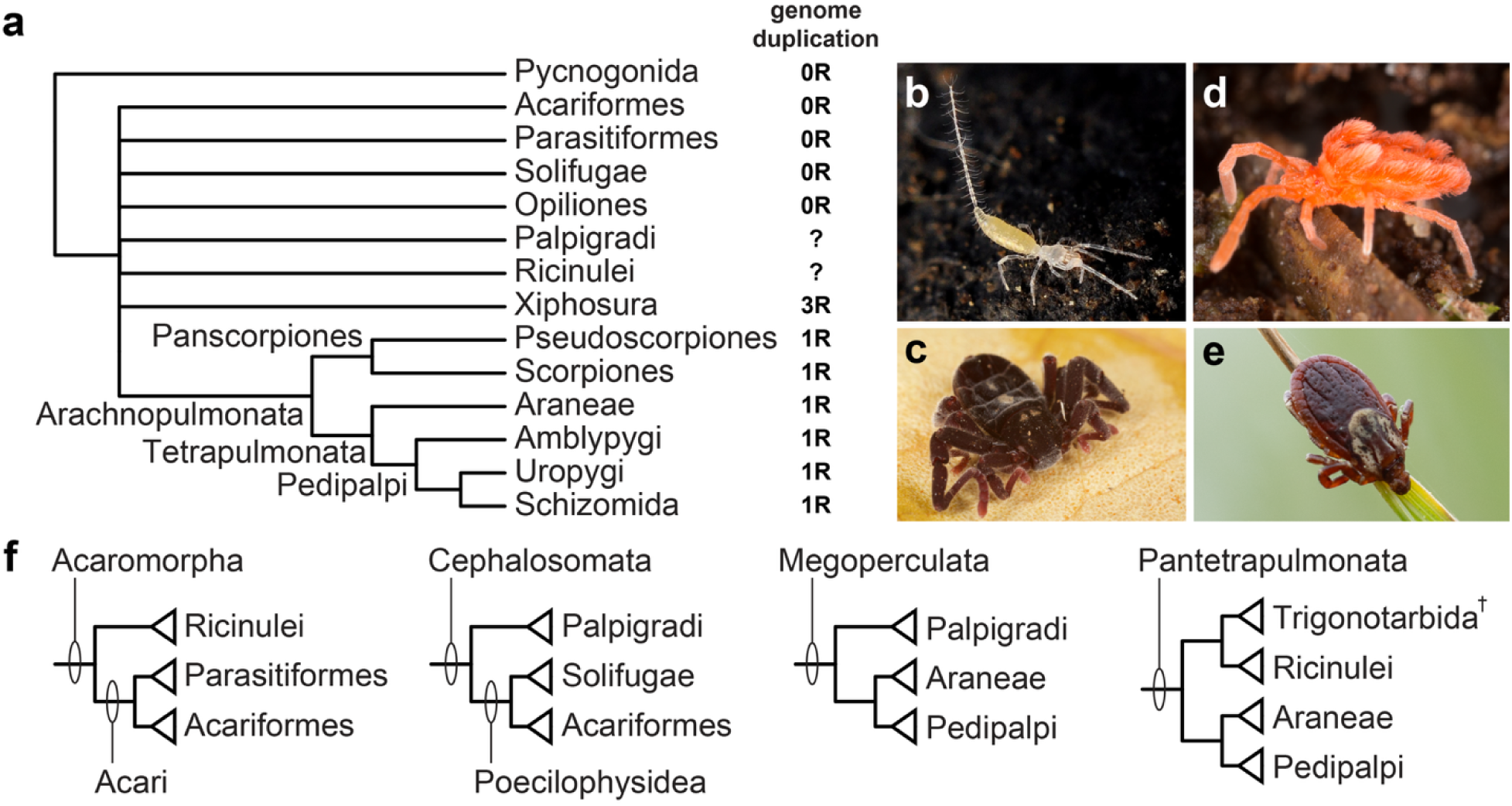
Unresolved and conflicting parts of the chelicerate tree of life. **a**, Simplified phylogeny of Chelicerata showing robustly resolved nodes and known distribution of whole genome duplication events. **b**-**e**, Live habitus of exemplars of unstable orders. **b**, *Eukoenenia spelaea* (Papigradi). **c**, *Cryptocellus* cf. *goodnighti* (Ricinulei). **d**, Chyzeriid mite (Acariformes). **e**, *Ixodes* tick (Parasitiformes). **f**, Competing hypotheses of Acariformes, Palpigradi, and Ricinulei phylogenetic placement from the morphological literature.

Resolving the phylogeny of the remaining chelicerates (the “apulmonate arachnids”; Figure 1a-e) has long been stymied by the combination of rapid radiation in the early Paleozoic and heterogeneous rates of evolution, phenomena which incur deep systematic biases (e.g., long branch attraction; incongruent gene trees) that impact both concatenation- and coalescent-based approaches to tree inference (Regier et al. 2010; Sharma, Kaluziak, et al. 2014; Ballesteros and Sharma 2019). In phylogenomic studies, the most unstable of the arachnid orders is consistently Acariformes, a species-rich taxon comprised of small-bodied mites, whose members include causal agents of human disease (e.g., scabies; tsutsugamushi disease; various allergens) and a variety of agricultural and horticultural pests (Grbić et al. 2011; Fischer and Walton 2014).

Acariformes include some of the oldest unambiguous terrestrial arachnid fossils and presently occupy nearly all ecological niches and habitats, spanning polar regions, soils, bird feathers, and plant tissues (Dunlop 2010; Klimov et al. 2025). At least four groups of mites have secondarily colonized aquatic habitats and one of these may have undergone tertiary recolonization of terrestrial habitats (Pepato et al. 2022). It is estimated that Acariformes may be the most species-rich order of arthropods, a distinction belied by the taxonomic shortfall in documenting their diversity (Klimov et al. 2018).

Morphological and molecular datasets have recovered conflicting placements of Acariformes. Cladistic analyses of anatomical data frequently recover Acariformes and Parasitiformes as sister groups, comprising the group Acari (Wheeler and Hayashi 1998; Shultz 2007). Acari is putatively united by the gnathosoma, a specialized anterior feeding apparatus containing the mouth, chelicerae, and palps, with ventral fusion of the palps to form an integrated unit, though recent work has suggested multiple independent origins of this complex (Bolton 2025). Acari in turn is recovered as the sister group of Ricinulei, forming Acaromorpha (Figure 1f), a taxon united by the presence of a hexapodous larva (Shultz 2007). A minority view instead considers Acariformes as the sister group of Solifugae, an order of large-bodied, predatory arachnids (Pepato et al. 2010; Dunlop et al. 2012; Garwood and Dunlop 2014). This grouping, Poecilophysidea, is united by the presence of a sejugal furrow between the second and third walking leg segments, and is in turn considered to be sister group to the miniaturized order Palpigradi (forming the clade Cephalosomata; Figure 1f). This trio of lineages shares the organization of the anterior segments as three tagmatic units (pro-, meso-, and metapeltidium), rather than the traditional seven-segmented prosoma of most chelicerates (Pepato et al. 2010; Dunlop et al. 2012). Molecular data have supported an array of possible placements for Acariformes, and often as sister group to remaining Euchelicerata (the non-sea spider chelicerates; Figure 1a) (Sharma, Kaluziak, et al. 2014; Ballesteros and Sharma 2019; Ballesteros et al. 2022). Some phylogenies initially recovered the monophyly of Acari (Lozano-Fernandez et al. 2019; Howard et al. 2020), but this result is destabilized by the addition of basally branching groups of Parasitiformes (e.g., Opilioacaridae, the putative sister group to the remaining parasitiforms) or is not supported in sensitivity analyses incorporating taxon permutation, suggesting that Acari monophyly is a long branch attraction artifact (Ballesteros et al. 2019; Ballesteros et al. 2022; Ontano et al. 2022; Domènech et al. 2026). Bayesian inference approaches using site heterogeneous models (e.g., CAT+GTR) have sometimes recovered support for the Poecilophysidea hypothesis (Sharma, Kaluziak, et al. 2014; Ballesteros et al. 2022), as well as Cephalosomata in one study sampling Palpigradi (Ballesteros et al. 2022). These dissonant resolutions of acariform placement in the arachnid tree of life are difficult to adjudicate in the absence of strong external evidence for phylogenetic accuracy.

A potential solution to breaking soft polytomies is to survey datasets for rare genomic changes, a class of homoplasy-limited characters that account for genome architecture (Rokas and Holland 2000). Such characters include whole genome duplication (WGD) events, shared intergenic insertions of retrotransposons, and shared gene family expansions (Hazkani-Covo 2009; Salichos and Rokas 2014; Fröbius and Funch 2017; Schwager et al. 2017). Recently, chromosomal fusion-with-mixing events have also shown remarkable utility as a complex phylogenetic character at the base of the animal tree of life, due to the improbability of reverting to the ancestral unmixed condition after extensive mixing of genomic regions (Schultz et al. 2023; Lewin, Sakagami, et al. 2025). The use of such complex characters that exhibit deeply asymmetrical probability of gain versus loss provides a considerable advantage in distinguishing between competing hypotheses, which serves the dual goals of phylogenetic resolution as well as arbitrating the accuracy of substitution models. To date, the use of these data classes has been limited to a subset of chelicerate taxa, due to the unavailability of genomes representing the major arachnid groups that constitute the polytomy. To leverage the phylogenetic potential of rare genomic changes, we established a series of genomic resources for key groups of chelicerates; together with available datasets, these new resources enable the sampling of all eight lineages comprising the polytomy (Figure 1a).

## Results

### Genome sequencing and assembly of key arachnid taxa

We generated genome assemblies for a Ricinulei (*Cryptocellus* cf. *goodnighti*), a palpigrade (*Prokoenenia wheeleri*), a scorpion (*Centruroides sculpturatus*), a mesothele spider (*Liphistius suwat*) and a mygalomorph spider (*Aphonopelma hentzi*) using a combination of field collection, PacBio HiFi sequencing, and (excepting the palpigrade) Dovetail HiC scaffolding of the genome assembly (Supplementary Figure S1; Supplementary Tables S1, S2). Although several arachnopulmonate genomes are available, the generation of new genomes for the scorpion and tarantula species was predicated upon their ongoing relevance to developmental genomics (Sharma, Schwager, et al. 2014; Leite et al. 2016; Schwager et al. 2017; Sharma 2017; Leite et al. 2018; Setton et al. 2024). In brief, the haploid assemblies for these taxa varied in size from 797 Mb (*P. wheeleri*) to 5.75 Gb (*A. hentzi*). BUSCO completeness of these assemblies spanned 93.8-98.5%. Arachnopulmonates (spiders and scorpion) exhibited 4.8-7.3% duplication of BUSCO genes, consistent with previously reported assemblies, whereas *C.* cf. *goodnighti* exhibited a 3.75% BUSCO duplication rate. Where available, karyotype data for the genus or the species accorded closely with the number of pseudochromosomes recovered by HiC scaffolding for the arachnopulmonates and the Ricinulei.

Due to miniaturization in Palpigradi (with most species not exceeding 1.5-2 mm in length), the palpigrade genome required pooling of several field-collected individuals (n=21) and a hybrid dataset to generate a highly complete assembly (i.e., with and without genome amplification). As a result, 52.1% of BUSCOs were duplicated in this assembly, which is consistent with artifactually high heterozygosity resulting from pooling and additional artifactual errors introduced by genome amplification. Due to the scaffold-level quality of this genome, it was excluded from comparisons of genome architecture and macrosynteny.

### The arachnopulmonate whole genome duplication excludes Xiphosura, Ricinulei, and Palpigradi

Shared whole genome duplication events constitute a powerful character system for phylogenetic inference due to their systemic nature and the persistence of their genomic signature over long temporal intervals. The discovery of a shared WGD uniting the most recent common ancestor of spiders and scorpions strongly corroborated the Arachnopulmonata hypothesis, which had been recurrently recovered by phylogenomic analyses (Sharma, Kaluziak, et al. 2014; Schwager et al. 2017). The WGD event also proved essential to the placement of Pseudoscorpiones, a long-branch order that alternatively grouped with Scorpiones or Acariformes across phylogenomic studies. Pseudoscorpions were also recently shown to share the arachnopulmonate WGD event, whereas Acariformes, Parasitiformes, Solifugae, Opiliones, and Pycnogonida all exhibit the plesiomorphic unduplicated genome condition (Ontano et al. 2021; Gainett, Klementz, Setton, et al. 2024; Klementz et al. 2025; Papadopoulos et al. 2025).

An untested group with respect to the arachnopulmonate duplication is Ricinulei (hooded tick spiders; Figure 1c), which externally resemble the book lung-bearing fossil order Trigonotarbida (Dunlop 2010). A clade formed by this pair of taxa is occasionally recovered as closely related to tetrapulmonates in cladistic analyses of morphological matrices (Dunlop 1998), whereas phylogenomic analyses have recovered Ricinulei in various unstable placements, including as the sister group of Xiphosura, of Solifugae, and in a clade with Solifugae and Opiliones (Sharma, Kaluziak, et al. 2014; Ballesteros et al. 2019; Lozano-Fernandez et al. 2019; Ballesteros et al. 2022) (Figure 1f). Similarly, the miniaturized order Palpigradi (microwhip scorpions) is recovered by morphological data as the sister group of the tetrapulmonates (Shultz 1990; Wolfe 2017) or closely related with apulmonate groups such as mites and solifuges (Garwood and Dunlop 2014) (Figure 1f). We reasoned that if either Ricinulei or Palpigradi were nested within Arachnopulmonata or closely related to Xiphosura, either or both could exhibit the signature of a shared WGD, as previously shown for pseudoscorpions. We surveyed 24 genomes (19 arachnids, three horseshoe crabs, one sea spider, one myriapod outgroup; Supplementary Table S2) for five gene clusters that have been shown to retain duplicated paralogs with high fidelity in animal genomes: the Hox, Nk, Iroquois, HRO, and SINE clusters (Laga-Trillo and Meyer 2001; Larroux et al. 2007; Nong et al. 2021; Aase-Remedios et al. 2023).

In the Ricinulei genome (Figure 2a) we recovered nine out of ten Hox genes (all but *proboscipedia*) in a single cluster spanning 3.84 Mb. The ricinuleid Nk genes were distributed as a single cluster comprising 4.89 Mb that contained single copies of all Nk genes except *Emx* (two copies), *Msx* (two copies), and *Nk6* (not found). A copy of *Msx* and *Noto* were separately found on a 9.57 Mb fragment on a separate pseudochromosome. Singletons of *Nk2.1* and *Nk2.2* were found on separate pseudochromosomes. Single copies of all three Iroquois genes were organized as a single cluster spanning 0.80 Mb. The HRO genes were distributed as one 1.71 Mb cluster with single copy homologs of three genes and a singleton of *Islet* on a separate chromosome. We found two of the three members of the SINE family of genes (*Six1/2* and *Six3/6*) on a single cluster comprising 0.156 Mb. In comparison to these patterns, genomes of arachnopulmonates consistently exhibited evidence for two clusters for all five gene families surveyed, whereas non-arachnopulmonate arachnids exhibited an unduplicated condition (Supplementary Figures S2, S3; Supplementary Tables S3-25). Despite the lack of a chromosome-level assembly, surveys of these genes in the palpigrade *P. wheeleri* revealed only single-copy homologs or minimally divergent paralogs that are attributable to heterozygosity in the hybrid assembly of this species (Figure 2b; Supplementary Figures S4-S8).

**Figure 2.**
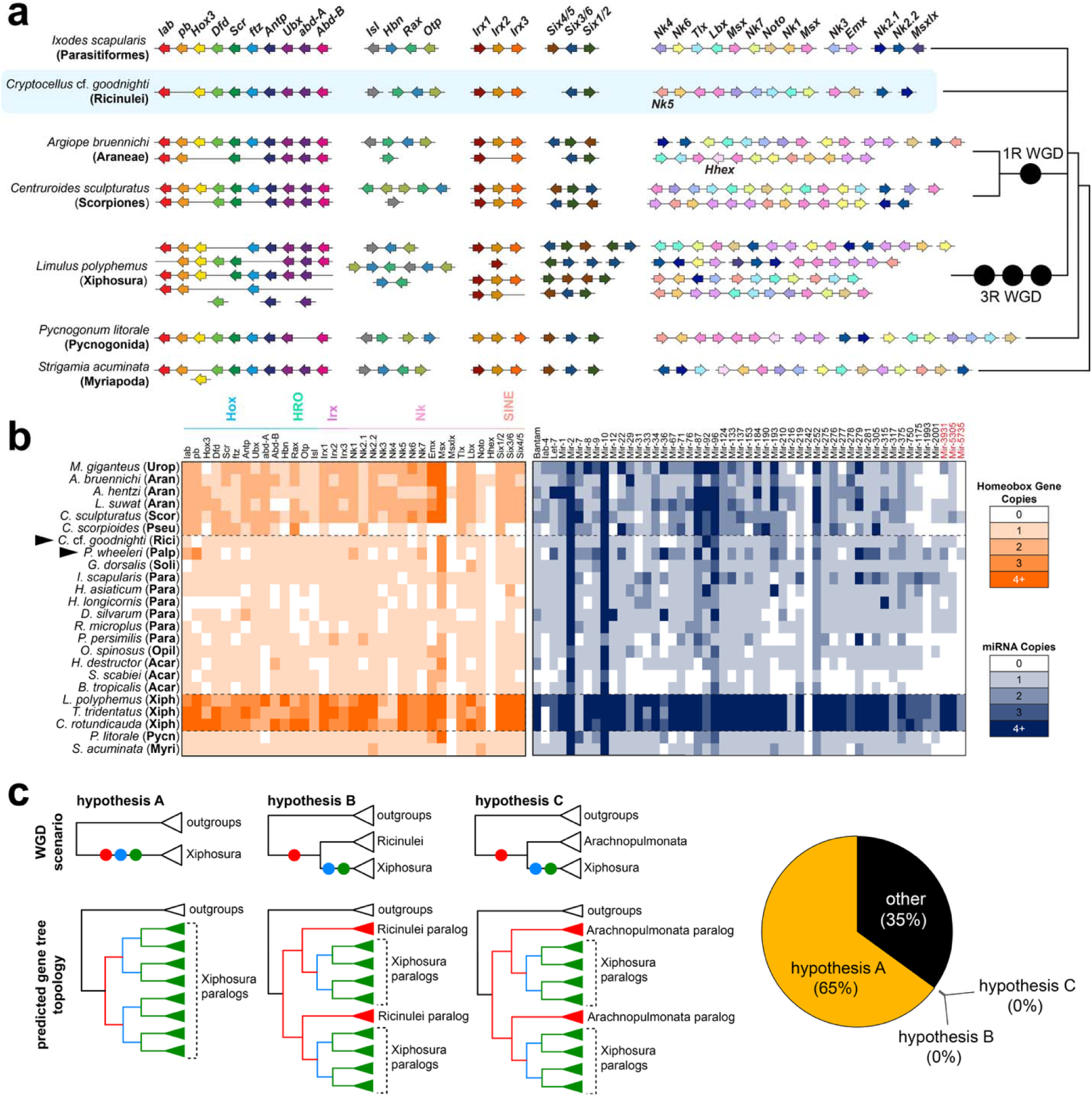
Signatures of whole genome duplications in chelicerate genomes. **a**, Architecture of Hox, HRO, Irx, SINE, and Nk clusters across chelicerate genomes reveals no signature of whole genome duplication in Ricinulei. **b**, Incidence of paralogy in homeobox genes (orange) and microRNA families (blue). **c**, Competing hypotheses for incidence of whole genome duplications on the branch subtending Xiphosura, with predicting gene tree topologies. Hypothesis A (three genome duplications restricted to Xiphosura) was supported by the majority of gene tree topologies (65%).

To validate these inferences, we examined the microRNA complements in these genomes, as microRNA copies can reliably capture the signature of ancient genome duplication (Leite et al. 2016; Umu et al. 2023). We compared the distributions of 48 conserved microRNA families and three chelicerate-specific families across the 24 arthropod genomes (Figure 2b; Supplementary Table S26). In the Ricinulei *C.* cf. *goodnighti*, we did not find any cases of shared microRNA duplications restricted to arachnopulmonates and Ricinulei. Patterns of microRNA duplication were ambiguous in the palpigrade, due to heterozygosity and the quality of the scaffold-level assembly.

The presence of single-copy genes alone does not rule out the possibility of an ancient, shared duplication, followed by lineage-specific gene losses. It has been postulated that one of the horseshoe crab WGD events may have been ancestral to a larger subset of chelicerate orders, with extensive losses of genes in the remaining arachnid orders (Munegowda et al. 2025). To test for this, we examined gene tree topologies of homologs from the four clusters that exhibit reliable signatures of duplication and lack tandem duplicates in apulmonate arachnid exemplars (Hox, HRO, Irx, and SINE). The Nk cluster was not included, as it had already undergone tandem duplications in several members prior to the diversification of Chelicerata (e.g., *Msx*; *Emx*) (Aase-Remedios et al. 2023). We interrogated the topological relationships of horseshoe crab paralogs specifically, to test the possibility that any arachnid order consistently grouped within the Xiphosura cluster (implying a signal of shared genome duplication). The majority of gene trees recovered the Xiphosura paralogs as single clusters (13/20) (Supplementary Figures S4-S8), consistent with previous analyses of Hox gene trees of Xiphosura that used a subset of arachnid orders as outgroups (Nong et al. 2021; Castellano et al. 2025). In cases where horseshoe crab copies did not form a single cluster, no single arachnid group was consistently associated with the horseshoe crabs. We also did not recover Palpigradi or Ricinulei sequences as consistently nested within the arachnopulmonate clusters. Taken together with the architecture of clusters in arachnid genomes, these results support the present definition of Arachnopulmonata and the separate instances of WGD events subtending the arachnopulmonates (1R) and the horseshoe crabs (3R), to the exclusion of all other chelicerate orders. As a corollary, our results conclusively rule out the Megoperculata and Pantetrapulmonata hypotheses (Figure 1f), further underscoring the limitations of anatomical datasets in predicting higher-level chelicerate relationships (Sharma et al. 2021; Gainett, Klementz, Setton, et al. 2024).

### Patterns of macrosynteny recapitulate conflicting signal within the apulmonate arachnid orders

The relationships of the apulmonate arachnid orders have historically been among the most adamantine to resolution, due to conflicting phylogenetic signal in molecular datasets. Acariformes is the least stable of these groups, owing to regressive evolution of several anatomical traits and accelerated rates of evolution that are related to miniaturization (Grbić et al. 2011). We explored macrosynteny of ancient linkage groups, a promising source of rare genomic changes in the form of chromosome fusion-with-mixing events, which are thought to be irreversible once gained (Schultz et al. 2023; Lewin, Sakagami, et al. 2025). To maximize informativeness for the focal ordinal relationships, we inferred a euchelicerate-specific ancestral linkage group using three apulmonate orders: *Sarcoptes scabiei* (Acariformes), *Ixodes scapularis* (Parasitiformes), and *Odiellus spinosus* (Opiliones).

Oxford plots supported the cohesion of ancestral linkage groups in representatives of all major lineages except Palpigradi (chromosome-level assembly not available) and Pseudoscorpiones (a member of Arachnopulmonata) (Supplementary Figs. S9-S34). We identified three informative patterns of chromosome fusion-with-mixing informing the euchelicerate polytomy. One mixing event (J ⊗ 2) was shared by Acariformes and Ricinulei; one event (A ⊗ C) was shared by Acariformes and Opiliones; and one other event (G ⊗ 23) was shared by Acariformes and Parasitiformes (Figure 3). Some of these mixing events proved challenging to interpret in the genome of the solifuge *Gluvia dorsalis*, which has a reduced karyotype (2n = 10) and as a result, reduced cohesion of the ancestral linkage groups. We therefore examined a highly contiguous genome of a second solifuge, *Paragaleodes pallidus* (Garcia et al. 2026); this species showed no evidence of any of these three identified fusion-with-mixing events, suggesting that putative shared states of fusion and mixing in the *G. dorsalis* genome are the result of homoplasy (Supplementary Figure S35). We additionally found two mixing events that superficially seemed to unite Acariformes and Ricinulei (I ⊗ 8 and 14 ⊗ 38), but closer examination of these events in *C.* cf. *goodnighti* showed that these events did not meet our minimum criteria of (1) at least 40 total genes involved in the mixing and (2) five swapping events between gene blocks required to explain the physical distribution of the genes in all taxa (Supplementary Figure S36).

**Figure 3.**
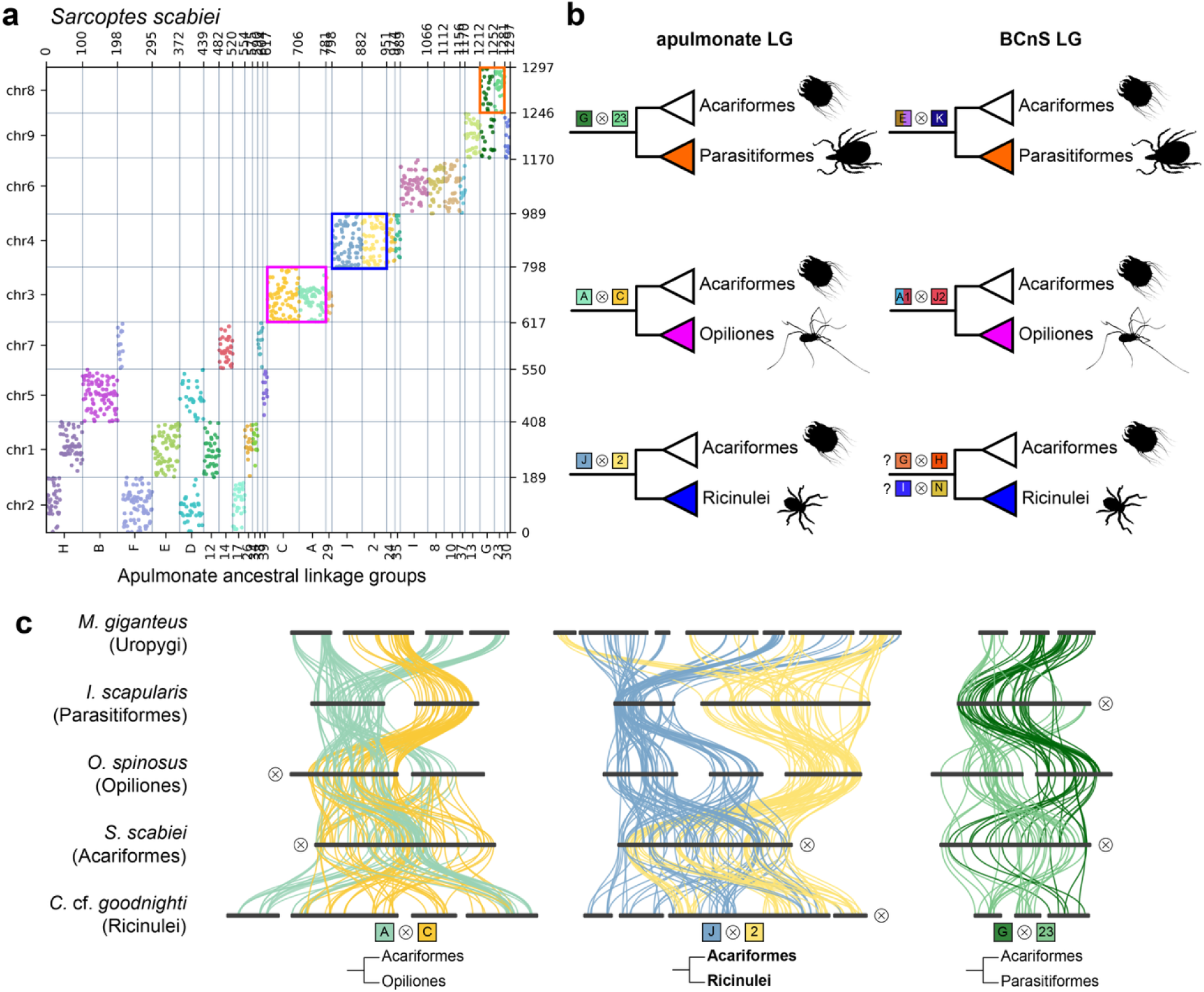
Chromosomal fusion-with-mixing supports mutually exclusive hypotheses. **a**, Oxford dot plot of an acariform mite *Sarcoptes scabiei* showing five fusion-with-mixing events that are shared by Acariformes and one other order. Colored outlines in the dot plot correspond to identity of the putative sister group shown to the right. **b**, Summary figure of the implied relationships resulting from informative mixing events under the apulmonate linkage group (left) and the BCnS linkage group (right) **c**, Ribbon plots showing phylogenetically informative mixing events inferred using the apulmonate linkage group. Black bars, chromosomes; colored lines, orthologous genes from ancestral linkage groups (ALGs) as shown in **a**. Crossed icons (⊗) next to chromosomes with interspersed colors indicate fusion-with-mixing events.

To rule out the possibility that these results were an artifact of linkage group construction, we performed the same analysis using a previously established Bilateria-Cnidaria-and-Sponge (BCnS) linkage group (Simakov et al. 2022). Our previous exploration of this linkage group for a subset of chelicerate genomes had found two fusion-with-mixing events with mutually exclusive outcomes, one supporting Acariformes + Opiliones (A1a ⊗ A1b ⊗ J2) and one supporting Acariformes + Parasitiformes (Ea ⊗ Eb ⊗ K) (Klementz et al. 2025). In the present analysis, we found that the inclusion of additional exemplars of Acariformes, Opiliones, and Parasitiformes did not reduce support for either competing hypothesis (Figure 3b; Supplementary Figures S37-S62). In addition to finding the same two fusion-with-mixing patterns across the surveyed taxa, we found two events partially shared by Acariformes + Ricinulei (G ⊗ H and I ⊗ N, albeit with partial mixing in *C.* cf. *goodnighti*, with a subset of genes in the G and H linkage groups and a subset of the N linkage group included in the mixing; Supplementary Figure S43), paralleling analytical outcomes using the euchelicerate linkage group.

### Patterns of macrosynteny are uninformative with respect to arachnid monophyly and arachnopulmonate relationships

The proposition that Arachnida is not monophyletic has stirred some controversy, not least among proponents of site heterogeneous model-based phylogenetics (Lozano-Fernandez et al. 2019; Howard et al. 2020), though this position has recently softened with the gradual recognition that even site heterogeneous models such as CAT+GTR struggle to recover Arachnida outside of a small and inconsistent minority of cases (Domènech et al. 2026), recapitulating the results of early explorations of arachnid datasets using PhyloBayes twelve years ago (Sharma, Kaluziak, et al. 2014). The visibility of this controversy led us to examine whether fusion-with-mixing events potentially support or refute arachnid monophyly as well. We mapped every fusion-with-mixing event in the euchelicerate linkage group on a tree topology of Chelicerata to infer the distribution of events potentially identifying the placement of Xiphosura. We found no shared events uniting the arachnid orders to the exclusion of Xiphosura, nor any events linking Xiphosura to any subset of arachnid orders (Figure 4), recapitulating a similar analysis with the BCnS linkage group with a subset of chelicerate orders (Klementz et al. 2025).

**Figure 4.**
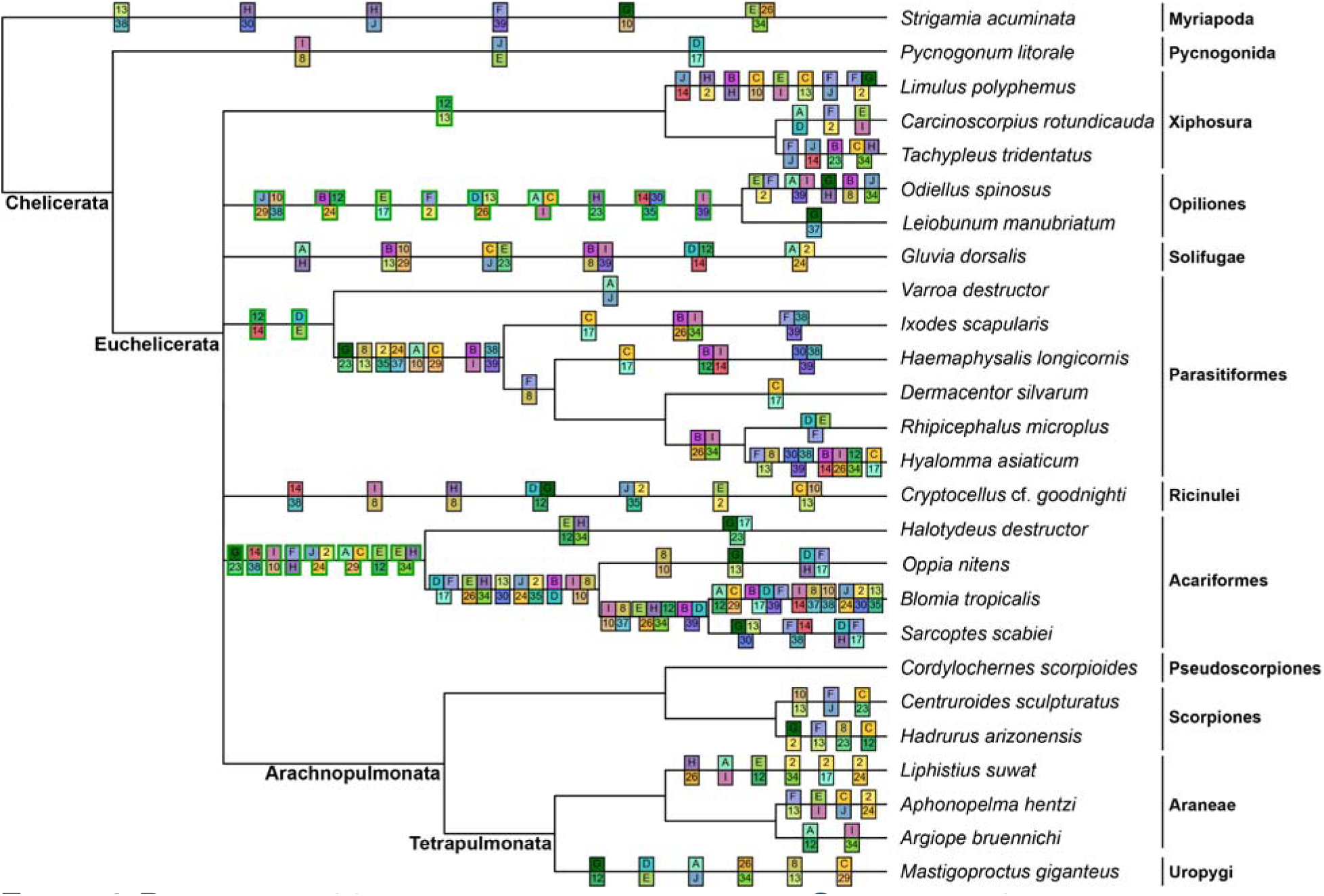
Distribution of fusion-with-mixing events across Chelicerata inferred using the apulmonate linkage group. Green outlines indicate events congruent with ordinal monophyly for orders sampled with multiple genomes. Note the absence of any mixing events supporting Arachnida, Euchelicerata, Arachnopulmonata, and Tetrapulmonata.

Numerous events were found in support of the monophyly of a subset of arachnid orders, with the exception of Pseudoscorpiones, which lacks strong signal of cohesion of ancestral linkage groups in the sole available chromosome-level genome. In addition, we found no events uniting Arachnopulmonata, Tetrapulmonata, or Euchelicerata, the three most strongly supported interordinal relationships within the chelicerates. All other fusion-with-mixing events observed were autapomorphic to individual species, or united well-established groupings within arachnid orders, such as Ixodida (nine events), Metastriata (the non-*Ixodes* Ixodidae; one event), and sarcoptiform mites within Acariformes (six events). These results recapitulate recent explorations of macrosyntenic patterns within Acariformes and Parasitiformes with an “Acari” linkage group, which revealed that diagnostic changes in fusion-with-mixing events occur within these arachnid lineages (Bhoi et al. 2026).

Taken together, the distribution of these events suggests that fusion-with-mixing events by themselves are imperfect readouts of deep phylogenetic relationships, as their occurrence does not reliably capture well-established splits in metazoan phylogeny (e.g., Euchelicerata; Tetrapulmonata).

### Evolutionary dynamics of lineage-restricted orthogroups reinforce conflicts among apulmonate arachnid orders

The three competing placements of Acariformes inferred by fusion-with-mixing differ markedly from the most common placement of this order in molecular phylogenetic analyses, which frequently place Acariformes as the sister group to the remaining Euchelicerata. We sought to distinguish these competing hypotheses using phylogenomics and orthogroup evolutionary dynamics. To this end, we generated a 94-taxon phylogenomic matrix using a set of 400-slowest evolving arthropod-specific homologs (BUSCOs). Maximum likelihood inference with a per mean site frequency (PMSF) model recapitulated relationships supported by previous studies, with Xiphosura nested within the arachnids, fast-evolving taxa (e.g., Acariformes; Parasitiformes) forming a grade near the base of the euchelicerates, and Acariformes as the sister group of the remaining Euchelicerata (Figure 5). We compared this benchmark tree topology to eight other additionally constrained trees reflecting competing hypotheses of chelicerate relationships from the literature, as well as from our analysis above: (1) Acaromorpha (=Acariformes + Parasitiformes + Ricinulei), (2) Acariformes + Ricinulei, (3) Acariformes + Opiliones, (4) Acari (=Acariformes + Parasitiformes), (5) Poecilophysidea (=Acariformes + Solifugae), (6) Cephalosomata (=Acariformes + Solifugae + Palpigradi), (7) Solifugae + Ricinulei + Opiliones, (8) Arachnida, (9) Xiphosura + Arachnopulmonata, and (10) Xiphosura + Ricinulei + Arachnopulmonata. We additionally compared the PMSF tree topology to the best unconstrained maximum likelihood tree with standard partitioning model-fitting. Compared to the unconstrained PMSF tree topology (with Acariformes sister group to the remaining Euchelicerata and Xiphosura nested inside the arachnids), Approximately Unbiased and Shimodaira-Hasegawa tests of monophyly rejected all competing hypotheses except Xiphosura + Arachnopulmonata (*p*_AU_ = 0.189; *p*_SH_ = 0.766) and the unconstrained tree with partitioned models (*p*_AU_ = 0.340; *p*_SH_ = 0.734; Supplementary Table S27). With respect to Acariformes, these results could alternatively reflect that Acariformes occupy a basally branching position with Euchelicerata, or that the results of these tests of monophyly are biased by model choice.

**Figure 5.**
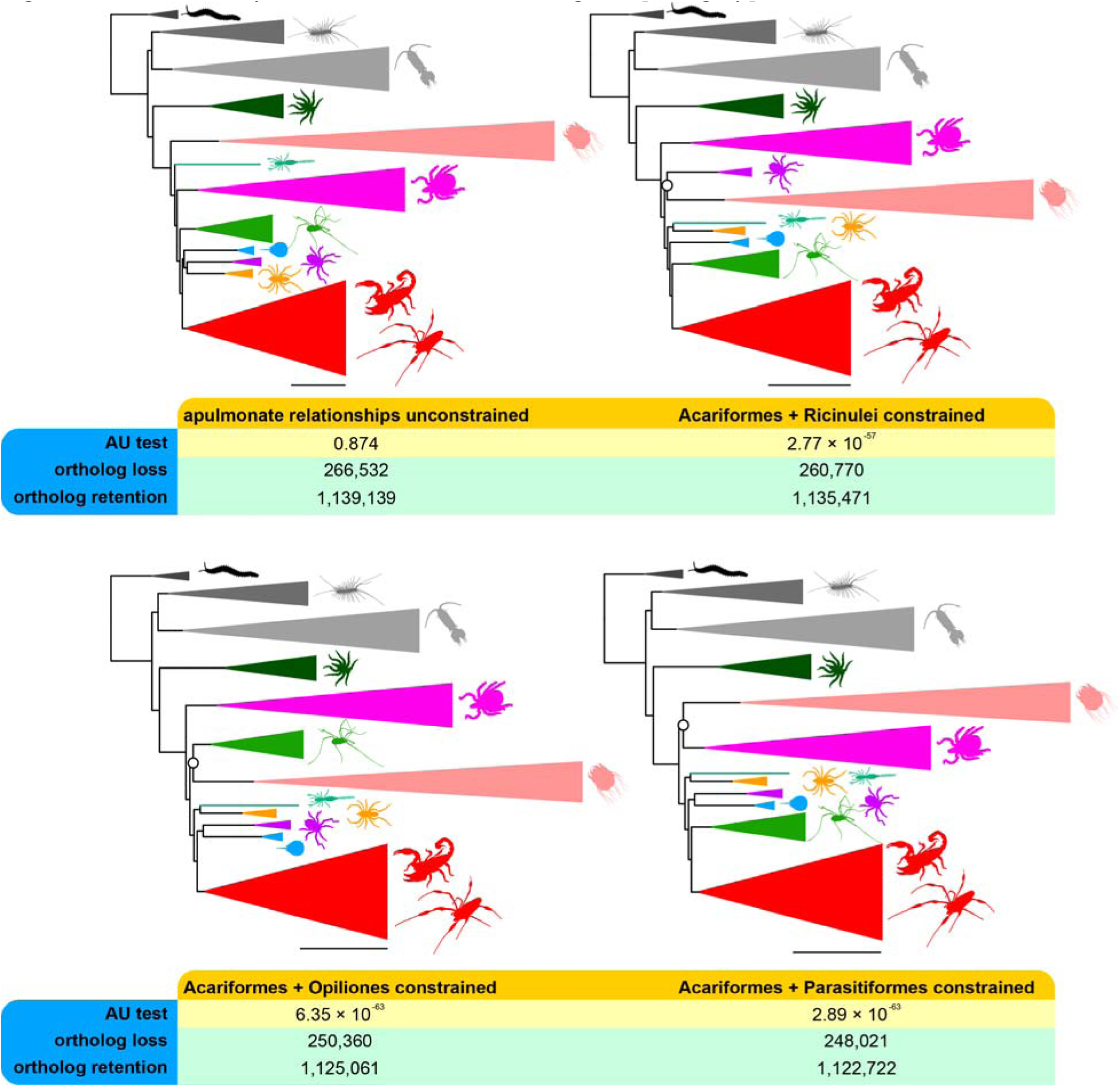
Phylogenomic trees with a nested placement of Acariformes within Euchelicerata recover a more parsimonious scenario of ortholog losses and retentions. Top row: Unconstrained 94-taxon maximum likelihood tree from 400 slowest-evolving BUSCO genes with PMSF model (left) *versus* maximum likelihood tree from the same matrix and constrained to recover Ricinulei + Acariformes (right). Bottom row: Maximum likelihood trees from the same matrix, constrained to recover Acariformes + Opiliones (left) and Acariformes + Parasitiformes (right). Silhouettes from PhyloPic.

We therefore examined the evolutionary dynamics of orthogroups across the tree to identify whether competing topologies exhibited measurably different signal in the distribution of orthogroup gains and losses. This approach assumes that losses of orthogroups are irreversible and applies a weighted parsimony framework to optimize gain and loss events of orthogroups on a phylogeny. We inferred orthogroups using OrthoFinder v.2.3.7 and compared their mapped distribution of losses on competing topologies. The tree topology constrained to recover Acariformes + Ricinulei was significantly more parsimonious than the unconstrained topology recovering Acariformes at the base of the euchelicerates (loss-to-retention ratio of 0.2297 *vs.* 0.2340, Fisher’s exact test, *p*_adj_ = 1.4 ×10^-8^; Fig. 4a, 4b; Supplementary Figure S63, Supplementary Table S28). The Acariformes + Ricinulei topology in turn was outperformed by the Acariformes + Parasitiformes topology (loss-to-retention ratio of 0.2209, Fisher’s exact test, *p*_adj_ = 2.9 ×10^-34^) and by the Acariformes + Opiliones topology (loss-to-retention ratio of 0.2225, Fisher’s exact test, *p*_adj_ = 1.0 ×10^-22^; Supplementary Figure S63). The latter two hypotheses were not statistically distinguishable (*p*_adj_ = 0.217).

As an ancillary analysis, we compared topologies constrained to recover either arachnid monophyly or the clade Xiphosura + Arachnopulmonata. The orthogroup loss-to-retention ratio was statistically indistinguishable in the maximum likelihood tree constrained to recover arachnid monophyly (0.2236) *versus* constrained to recover Xiphosura + Arachnopulmonata (0.2239; Fisher’s exact test, adjusted *p* = 0.641; Supplementary Figure S64, Supplementary Table S28).

Taken together, these results are congruent with inferences based on analyses of fusion-with-mixing events as well as molecular phylogenetic analyses, recapitulating the same conflicts in phylogenetic signal.

## Discussion

### Analyses of macrosynteny are unable to break the apulmonate polytomy

The approach of using chromosome fusion-with-mixing events as readouts of phylogenetic relationship has shown remarkable promise for resolving deep relationships, as exemplified by the placement of Ctenophora as the sister group to the remaining Metazoa, and of lophophorate monophyly, relationships that are supported by multiple independent fusion-with-mixing events (Schultz et al. 2023; Lewin, Sakagami, et al. 2025; but see Copley 2025). Chelicerata is a similar case, in that a large number of ingroup lineages comprises the focal polytomy and the ancient radiation of these lineages appears to have been rapid, as inferred from short internode distances in phylogenomic studies. As shown in this study, what makes the chelicerates different from such success stories is that exploration of macrosynteny reveals conflicting signal for mutually exclusive topologies, a result that is not attributable to the use of a specific linkage group. This is not the same as a lack of signal resulting from high cohesion of ancestral chromosomes, as previously shown for various invertebrate taxa (Schultz et al. 2023; Lewin, Sakagami, et al. 2025), because an uninformative trait is not conceptually problematic for any given phylogenetic character system. Rather, the incidence of fusion-with-mixing events supporting mutually exclusive topologies suggests that this character system can suffer the same shortcomings as every other character system, in that it is prone to homoplasy resulting from convergent evolution.

Why are chelicerates so prone to homoplasy in their genome architecture, when other phyla of invertebrates reflect picturesque cohesion of ancestral chromosomes conserved across Metazoa? One possibility is that the informativeness of fusion-with-mixing events varies as a function of karyotype, with lineages harboring a small number of chromosomes exhibiting extensive loss of signal. The intuitive effect of drastically reducing the number of haploid chromosomes is that numerous linkage groups are juxtaposed on contiguous genomic regions—the *sine qua non* of fusion-with-mixing events. Reduction in chromosome counts appears to be associated with loss of syntenic signal in some taxa, such as *Drosophila melanogaster* (2n=8) (Zimmermann et al. 2023). Indeed, some rapidly evolving groups of Acariformes and Parasitiformes exhibit small chromosome complements, as well near-total loss of signal resulting from chromosome rearrangements (e.g., *Tetranychus* urticae, 2n=6; *Phytoseuilus persimilis*, 2n=4; Eriophyoidea gen. sp., 2n=2) (Supplementary Figure S65). The rearrangement of gene order in mite genomes has previously been reported at a smaller scale in the mesostigmatan mite *Metaseiulus occidentalis*, which exhibits a totally atomized Hox gene cluster (Hoy et al. 2016). But the genomes of other mite taxa examined here appear to capture transitional states of chromosome fusion, with linkage groups exhibiting end-to-end joining and limited mixing, which are presumed to correspond to relatively recent fusion events (e.g., compare the sarcoptiform mites *Blomia tropicalis* and *Sarcoptes scabiei*; Supplementary Figures S11, S65). The pervasiveness of this phenomenon is difficult to assess across Chelicerata because the availability of chromosome-level genome assemblies remains highly asymmetrical. But karyotype counts are known to vary dramatically within arachnid orders (i.e., by an order of magnitude), even within genus-level taxa, which may portend lineage-specific patterns of signal loss in macrosynteny (Šťáhlavský et al. 2021; Král et al. 2025; Schöneberg et al. 2025).

Karyotype variation alone does not explain retention of macrosyntenic signal, as there does not appear to be a direct relationship between karyotype number and the degree of chromosome rearrangements (Lewin et al. 2024). Extensive chromosome rearrangements that disrupt ancestral linkage groups and Hox clusters have been reported in other invertebrate taxa, such as hexactinellid sponges, clitellate annelids, tunicates, and some groups of lepidopterans and Bryozoa, but these do not consistently exhibit low chromosome counts (Chen et al. 2023; Schultz et al. 2023; Lewin et al. 2024; Wright et al. 2024; Lewin, Liao, et al. 2025; Vargas-Chavez et al. 2025). Similarly, we observed derived lineages within arachnopulmonates exhibiting loss of signal despite a high karyotype count, such as in araneomorph spiders (e.g., *Argiope bruennichi*, 2n = 26 in females) and pseudoscorpions (e.g., *Cordylochernes scorpioides*; 2n = 48; Supplementary Figures S10, S14, S38, S42). Explorations of extensive chromosome rearrangements and loss of synteny have previously been reported across Araneomorphae, regardless of chromosome complement (Aase-Remedios et al. 2023). The precise mechanisms underlying the rearrangements of invertebrate ancestral linkage groups in the absence of dramatic karyotype reduction are not fully understood, but knowledge of their incidence suggests that broad sampling of focal exemplar taxa may be essential to identifying groups that retain the signal of ancestral patterns (Vargas-Chávez et al. 2025).

The outcome of our analyses is that macrosynteny by itself is unable to resolve competing hypotheses of apulmonate arachnid relationships. It is possible that sampling additional genomes may break this impasse by falsifying one or more competing hypotheses, as additional basally branching lineages are sampled. In addition, macrosynteny has shown some promise in the form of clear signal in the resolution of challenging relationships within species-rich arachnid orders and using custom, lineage-specific linkage groups (e.g., Acariformes and Parasitiformes; (Bhoi et al. 2026). Macrosynteny analyses often reveal numerous hits that cannot be assigned to specific linkage groups, as the majority of genes in metazoan genomes are not included in linkage group designs (Simakov et al. 2020; Schultz et al. 2023). Consequently, the extent of conserved macrosynteny might be underestimated, potentially obscuring patterns of chromosome evolution. The occurrence of unclassified homologs may also reduce the phylogenetic signal derived from genome architecture. Future improvements in inference of linkage group design, together with inclusion of genome assemblies for early-diverging lineages within arachnid orders, may help resolve unclassified hits and bolster analytical power.

### Relationships of the apulmonate arachnids

Of the three competing hypotheses of the acariform sister group, a relationship between Ricinulei and Acariformes to the exclusion of other taxa is seldom recovered in phylogenomic analyses, in contrast to Acari or Poecilophysidea, and no anatomical characters outright support this grouping (Figure 6). Acariformes and Ricinulei share a hexapodous larva, some degree of integration of mouthparts, and some degree of fusion of opisthosomal segments (e.g., Ricinulei have some elements of the gnathosoma, as well as diplotergites, the fusion of adjacent dorsal segments into plates), but these traits also occur in Parasitiformes and are likely evolutionarily labile (Dunlop and Lamsdell 2017). These three orders also share a subdivided femur with Solifugae, which has been argued to reflect an ancestral condition in Chelicerata (but is secondarily gained in some pseudoscorpions) (Shultz 1989; Klementz et al. 2024).

**Figure 6.**
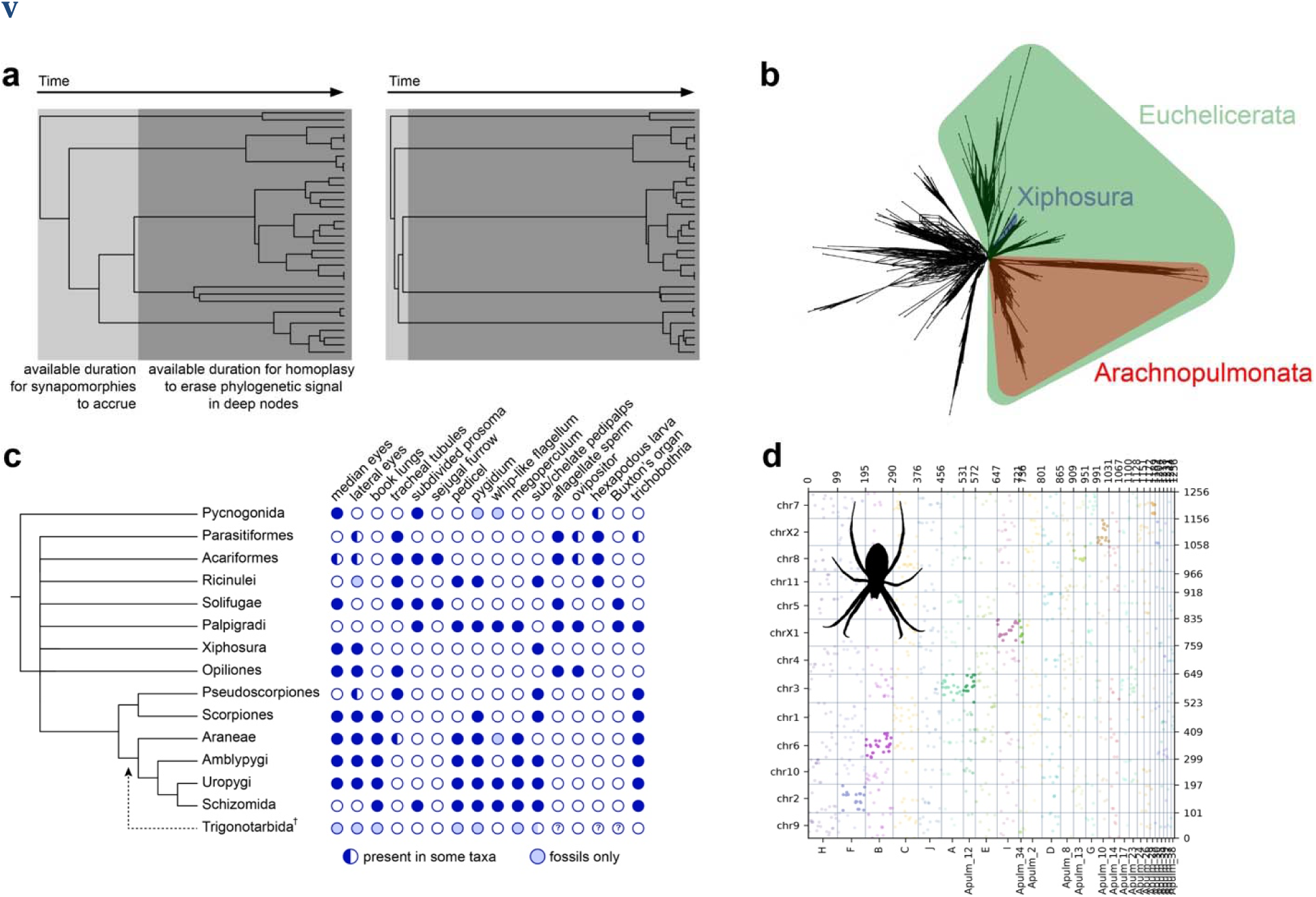
Challenging parts of the tree of life (“bushes”) are replete with systemic phylogenetic conflict. **a**, Recapitulation of the conceptual model of Rokas and Carroll (2006). The same tree topology is more difficult to resolve when a radiation is rapid and ancient (i.e., permitting little time for synapomorphies to accrue; light gray region), and followed by an extended period of time over which homoplasy can erase phylogenetic signal (dark gray region). **b**, Supernetwork of 200 BUSCO genes from the phylogenomic matrix, with reticulations corresponding to conflicting signal. **c**, Morphological trait evolution exhibits homoplasy and extensive character conflict across chelicerate phylogeny, regardless of topological resolution of the polytomy. Selected traits and character codings are drawn from the literature (Shultz 1990; Shultz 2007; Ballesteros et al. 2022). **d**, Oxford dot plot of the wasp spider *Argiope bruennichi* using the apulmonate linkage group, showing substantial loss in the cohesion of ancestral chromosomes. Silhouette of *A. bruennichi* from PhyloPic.

Similarly, a sister group relationship between Opiliones and Acariformes is seldom recovered in the literature. Across phylogenomic analyses, Opiliones are typically recovered in a clade with Ricinulei and Solifugae (Sharma, Kaluziak, et al. 2014; Howard et al. 2020) or as the sister group to Arachnopulmonata (Sharma, Kaluziak, et al. 2014; Ballesteros and Sharma 2019; Domènech et al. 2026), with the result highly contingent on matrix composition and model choice. Opiliones and Acariformes do not share any anatomical characters to the outright exclusion of other orders. The third topology supported by analyses of macrosynteny is consistent with the traditional grouping Acari, which has faced withering support with respect to both morphological and phylogenomic study (Ballesteros et al. 2019; Ballesteros et al. 2022; Bolton 2025; Domènech et al. 2026).

The only direction provided by surveys of rare genomic changes come from the unambiguous lack of a whole genome duplication in Ricinulei, which refutes morphological hypotheses uniting this group with the arachnopulmonates and places it within the apulmonate polytomy. This outcome implies that various anatomical traits of Ricinulei are the consequence of homoplasy, such as a characteristic spermatozoan with a coiled axoneme (shared with many arachnopulmonates), a chelate pedipalp (shared with most arachnopulmonates and Xiphosura), and a pygidium *sensu* Shultz (shared with various arachnopulmonates, the extinct order Trigonotarbida, Xiphosura, and Palpigradi). These results reinforce the conclusion that morphological data may be of limited value in disentangling higher-level arachnid relationships, with painfully uncomfortable implications for the placement of fossil taxa within Chelicerata (Sharma et al. 2021).

### Twenty years of bushes in the tree of life

Near the dawn of phylogenomics, a highly influential conceptual model was advanced to explain why molecular sequence data struggle with ancient rapid radiations, or “bush”-shaped tree topologies (Rokas and Carroll 2006). It was contended that the spacing of internode distances played a key role in the diagnosability of evolutionary relationships. Short and deep internodes at the base of the bush suffered from two challenges to resolution: a brief period for the accrual of synapomorphies and a long subsequent period for homoplasy to erase the phylogenetic signal (Figure 6a). The proposed solution to the ensuing polytomy at the base of the bush was to search for rare genomic changes that could resolve incongruence in matrices of sequence data (Rokas and Holland 2000; Rokas and Carroll 2006). Two decades later, understanding of the causes of phylogenetic incongruence, both biological and analytical, has advanced dramatically (Steenwyk et al. 2023). Various potential sources of incongruence have been interrogated in arachnid phylogeny, including the influence of data quantity, missing data, taxonomic sampling, outgroup selection, removal of fast-evolving taxa, incomplete lineage sorting, model misspecification, character recoding, alignment trimming, accidental in concatenation- versus coalescence-based tree inference, and alternative strategies of ortholog identification (Borner et al. 2014; Sharma, Kaluziak, et al. 2014; Ballesteros et al. 2019; Ballesteros and Sharma 2019; Lozano-Fernandez et al. 2019; Howard et al. 2020; Ontano et al. 2021; Ballesteros et al. 2022; Ban et al. 2022; Domènech et al. 2026). The emerging consensus is that the polytomy is not attributable to systematic or algorithmic error; it likely represents a true ancient and rapid radiation, with little time for the establishment of congruent phylogenetic signal across loci (Ballesteros et al. 2022).

This scenario contrasts with the textbook depiction of chelicerate evolution, which proposes a gradual and stepwise transition from Xiphosurida to Eurypterida to scorpions to the remaining arachnids (i.e., the strict interpretation that the chronological sequence of appearance of chelicerate fossils reflects a phylogenetic branching order). The inertia that has long maintained the textbook scenario in place has begun to erode as new fossils, developmental genetic data, comparative genomics, and recurring outcomes of phylogenomic studies have begun to rearrange relationships and the ensuing understanding of morphological trait evolution in Chelicerata (Schwager et al. 2017; Sharma 2017; Ballesteros and Sharma 2019; Noah et al. 2020; Ontano et al. 2021; Ban et al. 2022; Gainett, Klementz, Blaszczyk, et al. 2024; Gainett, Klementz, Setton, et al. 2024; Bolton 2025; Domènech et al. 2026; Lerosey-Aubril and Ortega-Hernández 2026).

This shift has included widespread acceptance of a derived placement of scorpions, as well as increasing receptivity to the possibility that arachnid evolution may have encompassed multiple terrestrialization events (Garwood and Dunlop 2023; Boyce and Nelsen 2025; Benítez Álvarez et al. 2026; Bolton 2026; Lamsdell 2026; Palacios et al. 2026).

An intriguing parallel therefore exists between all phylogenetic data classes at the base of Euchelicerata, in that conflicting signal affects not just phylogenomic matrices, but also morphology, patterns of macrosynteny, and the distribution of orthogroup gains and losses (Figure 6b-d). We postulate that chromosome fusion-with-mixing may follow the same principles as all other phylogenetic data classes in being subject to homoplasy and the vagaries of time between cladogenetic events. Like nucleotide or peptide sequences, bushes in the tree of life render chromosomal rearrangements little time to accrue in short internodes, but ample spans of time to endure homoplasy that can erase or mask those synapomorphies with conflicting signal (Figure 6d). The absence of fusion-with-mixing events uniting uncontroversial clades (e.g., Tetrapulmonata; Euchelicerata) in the chelicerate tree also suggests that phylogenetic signal in this character system is adventitious and may be highly contingent upon lineage-specific heterogeneity in the rate of chromosomal fusion events.

## Conclusion

Historic bursts of enthusiasm for new classes of rare genomic changes and putative homoplasy-limited character systems are predictably tempered by the gradual recognition of their limitations. This is exemplified by the waxing and waning of mitochondrial gene order, retrotransposon insertions, and microRNAs as putative silver bullets in phylogenetics (Hillis 1999; Masta et al. 2009; Thomson et al. 2014; Figueroa and Baco 2015; Kern et al. 2020). Rare genomic changes continue to offer promise in resolving bushes across the tree of life, as exemplified by nuclear mitochondrial introgression events in vertebrate phylogeny, as well as the whole genome duplication uniting Arachnopulmonata (Hazkani-Covo 2009; Schwager et al. 2017; Liang et al. 2018; Ontano et al. 2021). Nevertheless, the case of Chelicerata suggests that homoplasy in macrosyntenic patterns can obscure phylogenetic signal via incongruence in the case of challenging phylogenetic problems—just as with any other character system.

## Materials and Methods

### Field collection

*Aphonopelma hentzi* were collected in the Commanche National Grasslands near La Junta, Colorado (37.789478, −103.534146) in May 2021 by E.V.W.S. and P.P.S.; females were brought to the surface of burrows by pouring water into the entrances. Specimens of the palpigrade *Prokoenenia wheeleri* were collected in Cypress Creek Park, Travis County, Austin, Texas, United States (30.438459, −97.874670) in January 2023 by S.S.K., P.P.S., and Joseph Sardina, Jr., and in January 2024 by S.S.K. and Joseph Sardina, Jr.; individuals were collected from the soil layer under leaf litter. *Centruroides sculpturatus* were collected in Oracle, Arizona (32.609778, −97.874670) in January 2023 by S.S.K., P.P.S., and Joseph Sardina, Jr., and in 110.766528) at nighttime under UV light by Melody Albright. *Cryptocellus* cf. *goodnighti* were collected at the Smithsonian Tropical Research Institute field station, Bocas del Toro, Panama (9.354322, −97.874670) in January 2023 by S.S.K., P.P.S., and Joseph Sardina, Jr., and in 82.257926) by T.J.C. and B.A.S.D.M.; individuals were collected by sifting leaf litter and exported internationally under export permit number PA-01ARBG-039-2023.

### Sequencing and genome assembly

Accession and metadata for sequencing projects are provided in Supplementary Table S1. Genomic DNA was extracted using the Qiagen HMW DNA kit (Qiagen, Hilden, Germany) following manufacturer’s protocols. For the *A. hentzi* and the *L. suwat* genomes, a single adult female individual was used for the genome assembly. For the *C. sculpturatus*, an adult female was used for the genome assembly and a second female from the same population was extracted for HiC scaffolding. For *C.* cf. *goodnighti*, an adult male (FMNH 4600586) was extracted for genome assembly and three individuals from the same populations (two adult females, one juvenile; FMNH4600597, FMNH4600608, FMNH4600587) were extracted for HiC scaffolding. For the small-bodied palpigrade *P. wheeleri*, DNA was amplified using the REPLI-g Mini Kit (Qiagen, Hilden, Germany) with the manufacturer’s protocol. Separately, standard high molecular weight DNA extraction was performed for a pooled sample of 20 individuals.

Extractions for HiC scaffolding were trialed for *P. wheeleri* but could not be processed due to negligible capture efficiency.

Sequencing was performed on PacBio Sequel II, using standard manufacturer’s protocols for the Sequel II Sequencing Kit 2.0. Each library was sequenced on 2 SMRT Cells (8 M) in CCS mode for 30 h, except for the *A. hentzi* library, which was sequenced on 4 SMRT Cells due to the genome size. Analysis was performed with SMRT Link v10.1 software, requiring a minimum of three passes for CCS generation. Genome size and heterozygosity were estimated with a k-mer approach using GenomeScope (Vurture et al. 2017).

PacBio HiFi CCS reads were assembled independently using hifiasm v.0.15.1-r329 (Cheng et al. 2021) assemblers to generate contigs. The assembly graph generated by hifiasm was converted to a set of primary contigs in multi-fasta format using “awk ’/^S/{print “>”$2;print $3}’”.

Additional haplotigs and contig overlaps were removed with purge_dups v1.2.5 (Guan et al. 2020). Genome length and N50 were calculated using assemblyStatistics v.1.1.3 (Lin 2023). The assembly with longer N50 contiguity was used for further analyses.

### HiC sequencing and genome scaffolding

Omni-C library preparation and sequencing was performed using the Dovetail Omni-C Kit (Cantata Bio, Scotts Valley, California) for animal tissues using the manufacturer’s protocol. Briefly, frozen tissue was thoroughly ground in a mortar with pestle in liquid nitrogen followed by fixation of chromatin with formaldehyde. The chromatin was digested with DNase I, extracted, end-repaired and ligated to a biotinylated bridge adapter followed by proximity ligation of adapter ends. Crosslinks were reversed and the DNA was purified from proteins.

Biotin containing fragments were separated using streptavidin beads. The sequencing libraries tagged using unique dual indices and generated using Illumina-compatible adapters. The libraries were sequenced on an Illumina NovaSeq X+ platform at the Biotechnology Center (University of Wisconsin, Madison) to generate approximately 450 million 2×150 bp read pairs per sample.

DNA and library samples were quantified using Qubit 2.0 Fluorometer (Life Technologies, Carlsbad, CA, USA) and the fragment size distribution was assessed with a TapeStation Genomic DNA ScreenTape (Agilent Technologies).

The paired Omni-C sequenced data was mapped against the corresponding assembly with BWA-MEM (Li 2013). Ligation junctions among Omni-C pairs were detected using pairtools (Open2C et al. 2024) to sort the high-quality valid pairs. The resulting bam file was used to scaffold the genome using YaHS v.1.2a.1 (Zhou et al. 2023). Contact maps were generated using juicer_tools v.1.22.01 (Durand et al. 2016). Snail plots for the final assembly statistics were plotted with assembly-stats v17.02 (Challis et al. 2020) (Supplementary Fig. S35). Completeness of the assemblies was evaluated with BUSCO v5.4.5 by comparison with the arthropoda_odb10 database (Manni et al. 2021). Quality statistics for the genomes are provided in Supplementary Table S28.

### RNA sequencing

RNA sequencing was performed using sections of muscle, brain, and gonad tissue from the same individuals used for HMW DNA extractions. Tissue was homogenized in TRIzol reagent (Thermofisher, Waltham, US) and total RNA extracted following manufacturer’s protocols.

Stranded mRNA libraries were generated using the Illumina TruSeq library preparation kit, following manufacturer’s protocols. Accession data are provided in Supplementary Table S28.

### Genome annotation

A library of repetitive elements was compiled using RepeatModeler2 v.2.0.5 (Flynn et al. 2020) for each genome assembly. The identified repeats were soft masked using RepeatMasker v.4.1.5 (Smit et al. 2015). For annotation, de novo generated RNASeq data or from our previous works available on NCBI Sequence Read Archive were mapped to the assemblies using HISAT2 v.2.2.1 (Kim et al. 2019). Full gene structure annotations were predicted using BRAKER3 v.3.0.3 (Gabriel et al. 2024). For genomes without available RNASeq data, annotations were performed by mapping the masked genome to Arthropod protein database available at https://bioinf.uni-greifswald.de/bioinf/partitioned_odb11/Arthropoda.fa.gz.

### Gene orthology and microRNAs

To identify genes that capture the signature of whole genome duplication in chelicerate genomes, BLASTp searches were performed using queries of Hox, HRO, Irx, SINE, and Nk gene family homologs in the tick *I. scapularis* and the spider *Parasteatoda tepidariorum*, following recent approaches (Aase-Remedios et al. 2023; Klementz et al. 2025; Kulkarni et al. 2025). Sequences were extracted from all chelicerate genomes, as well as the centipede *S. acuminata* as an outgroup. Gene tree analyses were performed using IQ-Tree v.2(Nguyen et al. 2014) with automated model fitting and the tree topology was used to identify orthologs. Searches of microRNA families were implemented using MirMachine (Umu et al. 2023), with Chelicerata as the query node and using a combined model (protostome and deuterostome) for identifying conserved microRNAs.

### Macrosynteny and linkage group identification

To test for the incidence of phylogenetically informative chromosome fusion-with-mixing events, a customized linkage group set was developed using a three-way orthology search by finding reciprocal-best protein matches using DIAMOND v.2.1.24 (Buchfink et al. 2015) and OrthoFinder v.2.3.7 (Emms and Kelly 2019), as implemented in odp v.0.3.0 (Schultz et al. 2023). To resolve relationships of apulmonate arachnids, one representative of Acariformes (*S. scabiei*), Parasitiformes (*I. scapularis*), and Opiliones (*Odiellus spinosus*) were selected for inference of ancestral linkage groups. Genomes of taxa are high-quality, chromosome-level, and known to lack evidence of whole genome duplication. All linkage groups were generated using the “odp_nway_rbh” followed by “odp_rbh_to_alignments”. Briefly, odp_nway_rbh identifies orthologs using a reciprocal-best DIAMOND hits between proteins in the genome, guided by an annotation file for all chelicerate genomes. The false-discovery rate was set to α > 0.05 for finding the same sets of orthologs on the same chromosome and 1M iterations were performed. The chrom file was generated using a custom a script (gff_to_chrom.py) from the gff annotation file. Identical protein duplicates were removed using a custom script (removeDuplicateProt.py). The resulting 39 linkage groups were mapped against various chelicerate genomes and the centipede *Strigamia acuminata* (Myriapoda) as an outgroup to identify shared macrosyntenic patterns across Chelicerata. Fusion and mixing events that were uninformative (i.e., shared across all chelicerates) were collapsed as follows: 5 ⊗ 31 (=A); 1 ⊗ 22 (=B); 6 ⊗ 27 ⊗ 32 (=C); 16 ⊗ 20 (=D); 11 ⊗ 19 (=E); 3 ⊗ 36 ⊗ 39 (=F); 25 ⊗ 28 (=G); 4 ⊗ 15 (=H); 9 ⊗ 18 (=I); 7 ⊗ 21 (=J). The final ‘ApulmLG’ set consisted of of 27 linkage groups. Oxford dot plots and ribbon plots for all taxa were generated using odp v.0.3.0 to visualize syntenic patterns.

### Phylogenomics

Phylogenomic analysis was performed using genomes or transcriptomes of 94 panarthropod species (2 Onychophora; 7 Myriapoda; 12 Pancrustacea; 7 Pycnogonida; 3 Xiphosura; 11 Acariformes; 12 Parasitiformes; 1 Palpigradi; 3 Solifugae; 3 Ricinulei; 8 Opiliones; 4 Pseudoscorpiones; 5 Scorpiones; 2 Amblypygi; 2 Uropygi; 1 Schizomida; 11 Araneae). These taxa included all 27 genomes analyzed in this study; the remaining terminals were drawn from a recent phylogenomic analysis of Chelicerata (Ballesteros et al. 2022), prioritizing representation of major basal splits in each lineage and the inclusion of genomic resources with a BUSCO completeness score >85%. All 94 datasets were filtered using AlienIndex v.2.1 (https://github.com/josephryan/alien_index) to remove potential contamination and CD-HIT (Fu et al. 2012) with a similarity threshold of 0.98 to remove splice variants. BUSCO v.5 (Manni et al. 2021) was used to extract putative single-copy homologs from filtered datasets, using the arthropoda_odb10 dataset. For mapping results, a phylogenomic tree was inferred using the single-copy BUSCO genes, which were further filtered to retain the 400 slowest-evolving genes, following previous approaches (Ontano et al. 2022). Tree topologies were inferred with the per mean site frequency (PMSF) model in IQ-Tree v.2. The resulting tree was taken as the benchmark tree topology. For Approximately Unbiased and Shimodaira-Hasegawa tests with IQ-Tree v.2., the following relationships were tested as constraints: (1) Acaromorpha (=Acariformes + Parasitiformes + Ricinulei), (2) Acariformes + Ricinulei, (3) Acariformes + Opiliones, (4) Acari (=Acariformes + Parasitiformes), (5) Poecilophysidea (=Acariformes + Solifugae), (6) Cephalosomata (=Acariformes + Solifugae + Palpigradi), (7) Arachnida, (8) Xiphosura + Arachnopulmonata, and (9) Xiphosura + Ricinulei + Arachnopulmonata.

### Ancestral state reconstruction of orthogroups using Dollo parsimony

Orthogroup (OG) presence/absence was reconstructed across the phylogeny using a custom Dollo parsimony implementation in R (https://github.com/scriptomika/Arachnids), applied to a binary species × OG character matrix (OrthoFinder table Orthogroups.GeneCount.tsv) and a rooted phylogeny. Under Dollo parsimony, each character (OG) can be gained only once but lost independently multiple times. For each OG, the algorithm first identified tips scored as present, then traversed the tree upward to infer ancestral states. Direct parent nodes of present-state tips were marked present, followed by all internal nodes whose descendant sets included at least one present tip. The MRCA of all present-state nodes was then computed and added to the present list if applicable, along with any intermediate nodes connecting it to already-identified present nodes. Finally, the root was assigned the present state only if both of its immediate descendant subtrees contained at least one present tip, enforcing the single-gain assumption. The procedure was applied across all OGs, producing a node × OG matrix of inferred ancestral states. Each node of the tree was then classified by comparing inferred states to its immediate ancestral nodes: labeled as *gain* (absent → present), *loss* (present → absent), *inherited*, or *nonexistent*. To evaluate which topology required significantly fewer OG losses, pairwise Fisher’s exact tests were applied to a 2×2 contingency table of total retained versus lost OGs under each tree hypothesis, with p-values adjusted for multiple comparisons. Effect sizes were quantified as the difference in loss-to-retention ratios between each pair of topologies. The topology requiring significantly fewer losses was interpreted as the better-supported hypothesis under Dollo parsimony.

## Acknowledgements

Joseph Sardina, Jr. and Melody Albright assisted with field collection of specimens. Discussions with Georg Brenneis, Gonzalo Giribet, Nikolaos Papadopoulos, and Ward C. Wheeler refined the ideas presented. Sequencing was performed at the University of California-Irvine Genomics Research and Technology Hub and the BioTechnology Center (UW-Madison). High-throughput computational analyses were performed through the Bioinformatics Resource Center (BRC) of UW-Madison and through the Pegasus High Performance Computing Cluster of the George Washington University. This material is based on work supported by the National Science Foundation grant no. IOS-2016141 to P.P.S and a U.S.-Egypt Science and Technology Joint Fund award to P.P.S. and M.A.R. S.S.K was supported by the ANRF Ramanujan Fellowship (RJF/2023/000045) and an Early Career Research Grant (ANRF/ECRG/2024/000947/LS). Additional support was provided by National Science Foundation grant no. DEB-2154246 to G.H.

## Author contributions

S.S.K, B.C.K., and P.P.S. designed the project. P.P.S. supervised the work. B.A.S.D.B. and

T.J.C. performed field collection of the Ricinulei. S.S.K. performed the genome sequencing, assemblies, and macrosynteny analyses. R.M.V. performed the genome sequencing of the palpigrade. S.S.K., B.C.K., K.M.A., E.M.L., S.M.N., and P.P.S. analyzed macrosyntenic patterns. B.C.K., E.M.L., S.M.N., and P.P.S. analyzed gene duplications. J.A.B. and C.E.S.L. performed the phylogenomic analyses. M.S.P. and D.C.P. performed the orthogroup gain/loss analyses. B.C.K., G.G., and E.V.W.S. contributed RNAseq data. G.H. provided computational resources. M.K.H., M.A.R.-H., and P.P.S. obtained funding. P.P.S wrote the manuscript with input from all co-authors. All co-authors edited the manuscript.

---

**Supplementary Figure S1.** Snail plots and HiC contact maps for newly sequenced arachnid genomes. Numbers below contact maps indicate number of large pseudomolecules and percentage of genome sequence mapped to large pseudomolecules.

**Supplementary Figure S2.** Architecture of evolutionarily conserved gene clusters in genomes of Arachnopulmonata.

**Supplementary Figure S3.** Architecture of evolutionarily conserved gene clusters in genomes of apulmonate arachnids. Note the atomization of the Hox cluster in the fast-evolving Mesostigmata genome.

**Supplementary Figure S4.** Maximum likelihood tree topology of Hox cluster genes. **Supplementary Figure S5.** Maximum likelihood tree topology of HRO cluster genes. **Supplementary Figure S6.** Maximum likelihood tree topology of Irx cluster genes.

**Supplementary Figure S7.** Maximum likelihood tree topology of SINE cluster genes.

**Supplementary Figure S8.** Maximum likelihood tree topology of Nk cluster genes.

**Supplementary Figure S9.** Oxford dot plot showing the location of apulmonate ancestral linkage group (ALG) homologs in the genome of the mygalomorph spider *Aphonopelma hentzi*. Colors correspond to individual ALGs, with solid color indicating significant hits (*p <* 0.05) and transparent color indicating insignificant hits.

**Supplementary Figure S10.** Oxford dot plot showing the location of apulmonate ancestral linkage group (ALG) homologs in the genome of the araneomorph spider *Argiope bruennichi*. Colors correspond to individual ALGs, with solid color indicating significant hits (*p <* 0.05) and transparent color indicating insignificant hits.

**Supplementary Figure S11.** Oxford dot plot showing the location of apulmonate ancestral linkage group (ALG) homologs in the genome of the acariform mite *Blomia tropicalis*. Colors correspond to individual ALGs, with solid color indicating significant hits (*p <* 0.05) and transparent color indicating insignificant hits.

**Supplementary Figure S12.** Oxford dot plot showing the location of apulmonate ancestral linkage group (ALG) homologs in the genome of the horseshoe crab *Carcinoscorpius rotundicauda*. Colors correspond to individual ALGs, with solid color indicating significant hits (*p <* 0.05) and transparent color indicating insignificant hits.

**Supplementary Figure S13.** Oxford dot plot showing the location of apulmonate ancestral linkage group (ALG) homologs in the genome of the scorpion *Centruroides sculpturatus*. Colors correspond to individual ALGs, with solid color indicating significant hits (*p <* 0.05) and transparent color indicating insignificant hits.

**Supplementary Figure S14.** Oxford dot plot showing the location of apulmonate ancestral linkage group (ALG) homologs in the genome of the pseudoscorpion *Cordylochernes scorpioides*. Colors correspond to individual ALGs, with solid color indicating significant hits (*p <* 0.05) and transparent color indicating insignificant hits.

**Supplementary Figure S15.** Oxford dot plot showing the location of apulmonate ancestral linkage group (ALG) homologs in the genome of the ricinuleid *Cryptocellus* cf. *goodnighti*. Colors correspond to individual ALGs, with solid color indicating significant hits (*p <* 0.05) and transparent color indicating insignificant hits.

**Supplementary Figure S16.** Oxford dot plot showing the location of apulmonate ancestral linkage group (ALG) homologs in the genome of the parasitiform tick *Dermacentor silvarum*. Colors correspond to individual ALGs, with solid color indicating significant hits (*p <* 0.05) and transparent color indicating insignificant hits.

**Supplementary Figure S17.** Oxford dot plot showing the location of apulmonate ancestral linkage group (ALG) homologs in the genome of the solifuge *Gluvia dorsalis*. Colors correspond to individual ALGs, with solid color indicating significant hits (*p <* 0.05) and transparent color indicating insignificant hits.

**Supplementary Figure S18.** Oxford dot plot showing the location of apulmonate ancestral linkage group (ALG) homologs in the genome of the scorpion *Hadrurus arizonensis*. Colors correspond to individual ALGs, with solid color indicating significant hits (*p <* 0.05) and transparent color indicating insignificant hits.

**Supplementary Figure S19.** Oxford dot plot showing the location of apulmonate ancestral linkage group (ALG) homologs in the genome of the parasitiform tick *Haemaphysalis longicornis*. Colors correspond to individual ALGs, with solid color indicating significant hits (*p <* 0.05) and transparent color indicating insignificant hits.

**Supplementary Figure S20.** Oxford dot plot showing the location of apulmonate ancestral linkage group (ALG) homologs in the genome of the acariform mite *Halotydeus destructor*. Colors correspond to individual ALGs, with solid color indicating significant hits (*p <* 0.05) and transparent color indicating insignificant hits.

**Supplementary Figure S21.** Oxford dot plot showing the location of apulmonate ancestral linkage group (ALG) homologs in the genome of the parasitiform tick *Hyalomma asiaticum*. Colors correspond to individual ALGs, with solid color indicating significant hits (*p <* 0.05) and transparent color indicating insignificant hits.

**Supplementary Figure S22.** Oxford dot plot showing the location of apulmonate ancestral linkage group (ALG) homologs in the genome of the parasitiform tick *Ixodes scapularis*. Colors correspond to individual ALGs, with solid color indicating significant hits (*p <* 0.05) and transparent color indicating insignificant hits.

**Supplementary Figure S23.** Oxford dot plot showing the location of apulmonate ancestral linkage group (ALG) homologs in the genome of the harvestman *Leiobunum manubriatum*. Colors correspond to individual ALGs, with solid color indicating significant hits (*p <* 0.05) and transparent color indicating insignificant hits.

**Supplementary Figure S24.** Oxford dot plot showing the location of apulmonate ancestral linkage group (ALG) homologs in the genome of the horseshoe crab *Limulus polyphemus*. Colors correspond to individual ALGs, with solid color indicating significant hits (*p <* 0.05) and transparent color indicating insignificant hits.

**Supplementary Figure S25.** Oxford dot plot showing the location of apulmonate ancestral linkage group (ALG) homologs in the genome of the Mesothelae spider *Liphistius suwat*. Colors correspond to individual ALGs, with solid color indicating significant hits (*p <* 0.05) and transparent color indicating insignificant hits.

**Supplementary Figure S26.** Oxford dot plot showing the location of apulmonate ancestral linkage group (ALG) homologs in the genome of the vinegaroon *Mastigoproctus giganteus*. Colors correspond to individual ALGs, with solid color indicating significant hits (*p <* 0.05) and transparent color indicating insignificant hits.

**Supplementary Figure S27.** Oxford dot plot showing the location of apulmonate ancestral linkage group (ALG) homologs in the genome of the harvestman *Odiellus spinosus*. Colors correspond to individual ALGs, with solid color indicating significant hits (*p <* 0.05) and transparent color indicating insignificant hits.

**Supplementary Figure S28.** Oxford dot plot showing the location of apulmonate ancestral linkage group (ALG) homologs in the genome of the acariform mite *Oppia nitens*. Colors correspond to individual ALGs, with solid color indicating significant hits (*p <* 0.05) and transparent color indicating insignificant hits.

**Supplementary Figure S29.** Oxford dot plot showing the location of apulmonate ancestral linkage group (ALG) homologs in the genome of the sea spider *Pycnogonum litorale*. Colors correspond to individual ALGs, with solid color indicating significant hits (*p <* 0.05) and transparent color indicating insignificant hits.

**Supplementary Figure S30.** Oxford dot plot showing the location of apulmonate ancestral linkage group (ALG) homologs in the genome of the parasitiform tick *Rhipicephalus microplus*. Colors correspond to individual ALGs, with solid color indicating significant hits (*p <* 0.05) and transparent color indicating insignificant hits.

**Supplementary Figure S31.** Oxford dot plot showing the location of apulmonate ancestral linkage group (ALG) homologs in the genome of the acariform mite *Sarcoptes scabiei*. Colors correspond to individual ALGs, with solid color indicating significant hits (*p <* 0.05) and transparent color indicating insignificant hits.

**Supplementary Figure S32.** Oxford dot plot showing the location of apulmonate ancestral linkage group (ALG) homologs in the genome of the centipede *Strigamia acuminata*. Colors correspond to individual ALGs, with solid color indicating significant hits (*p <* 0.05) and transparent color indicating insignificant hits.

**Supplementary Figure S33.** Oxford dot plot showing the location of apulmonate ancestral linkage group (ALG) homologs in the genome of the horseshoe crab *Tachypleus tridentatus*. Colors correspond to individual ALGs, with solid color indicating significant hits (*p <* 0.05) and transparent color indicating insignificant hits.

**Supplementary Figure S34.** Oxford dot plot showing the location of apulmonate ancestral linkage group (ALG) homologs in the genome of the parasitiform mite *Varroa destructor*. Colors correspond to individual ALGs, with solid color indicating significant hits (*p <* 0.05) and transparent color indicating insignificant hits.

**Supplementary Figure S35.** Oxford dot plots showing the location of apulmonate ancestral linkage group (ALG) homologs in the genomes of two solifuges for the apulmonate (top row) and the BCnS (bottom row) linkage groups. Left column: *Gluvia dorsalis.* Right column: *Paragaleodes pallidus*. Colors correspond to individual ALGs, with solid color indicating significant hits (*p <* 0.05) and transparent color indicating insignificant hits.

**Supplementary Figure S36.** Insignificant cases of fusion-with-mixing in Acariformes + Ricinulei. **a**, Oxford dot plot of the hooded tick spider *Cryptocellus* cf. *goodnighti* showing fusion-with-mixing events that are putatively shared with Acariformes (blue outlines). Dotted lines indicate mixing events with fewer than 40 genes included and fewer than five minimum swap events. **b**, Oxford dot plot of the acariform mite *Sarcoptes scabiei* showing the same three events as in **a**. **c**, Summary of patterns shown in panels **a** and **b**. Question marks correspond to insignificant mixing. **d**, Ribbon plots showing insignificant mixing for two cases (I ⊗ 8 and 14 ⊗ 38).

**Supplementary Figure S37.** Oxford dot plot showing the location of BCnS ancestral linkage group (ALG) homologs in the genome of the mygalomorph spider *Aphonopelma hentzi*. Colors correspond to individual ALGs, with solid color indicating significant hits (p < 0.05) and transparent color indicating insignificant hits.

**Supplementary Figure S38.** Oxford dot plot showing the location of BCnS ancestral linkage group (ALG) homologs in the genome of the araneomorph spider *Argiope bruennichi*. Colors correspond to individual ALGs, with solid color indicating significant hits (p < 0.05) and transparent color indicating insignificant hits.

**Supplementary Figure S39.** Oxford dot plot showing the location of BCnS ancestral linkage group (ALG) homologs in the genome of the acariform mite *Blomia tropicalis*. Colors correspond to individual ALGs, with solid color indicating significant hits (p < 0.05) and transparent color indicating insignificant hits.

**Supplementary Figure S40.** Oxford dot plot showing the location of BCnS ancestral linkage group (ALG) homologs in the genome of the horseshoe crab *Carcinoscorpius rotundicauda*. Colors correspond to individual ALGs, with solid color indicating significant hits (p < 0.05) and transparent color indicating insignificant hits.

**Supplementary Figure S41.** Oxford dot plot showing the location of BCnS ancestral linkage group (ALG) homologs in the genome of the scorpion *Centruroides sculpturatus*. Colors correspond to individual ALGs, with solid color indicating significant hits (p < 0.05) and transparent color indicating insignificant hits.

**Supplementary Figure S42.** Oxford dot plot showing the location of BCnS ancestral linkage group (ALG) homologs in the genome of the pseudoscorpion *Cordylochernes scorpioides*. Colors correspond to individual ALGs, with solid color indicating significant hits (p < 0.05) and transparent color indicating insignificant hits.

**Supplementary Figure S43.** Oxford dot plot showing the location of BCnS ancestral linkage group (ALG) homologs in the genome of the ricinuleid *Cryptocellus* cf. *goodnighti*. Colors correspond to individual ALGs, with solid color indicating significant hits (p < 0.05) and transparent color indicating insignificant hits.

**Supplementary Figure S44.** Oxford dot plot showing the location of BCnS ancestral linkage group (ALG) homologs in the genome of the parasitiform tick *Dermacentor silvarum*. Colors correspond to individual ALGs, with solid color indicating significant hits (p < 0.05) and transparent color indicating insignificant hits.

**Supplementary Figure S45.** Oxford dot plot showing the location of BCnS ancestral linkage group (ALG) homologs in the genome of the solifuge *Gluvia dorsalis*. Colors correspond to individual ALGs, with solid color indicating significant hits (p < 0.05) and transparent color indicating insignificant hits.

**Supplementary Figure S46.** Oxford dot plot showing the location of BCnS ancestral linkage group (ALG) homologs in the genome of the scorpion *Hadrurus arizonensis*. Colors correspond to individual ALGs, with solid color indicating significant hits (p < 0.05) and transparent color indicating insignificant hits.

**Supplementary Figure S47.** Oxford dot plot showing the location of BCnS ancestral linkage group (ALG) homologs in the genome of the parasitiform tick *Haemaphysalis longicornis*. Colors correspond to individual ALGs, with solid color indicating significant hits (p < 0.05) and transparent color indicating insignificant hits.

**Supplementary Figure S48.** Oxford dot plot showing the location of BCnS ancestral linkage group (ALG) homologs in the genome of the acariform mite *Halotydeus destructor*. Colors correspond to individual ALGs, with solid color indicating significant hits (p < 0.05) and transparent color indicating insignificant hits.

**Supplementary Figure S49.** Oxford dot plot showing the location of BCnS ancestral linkage group (ALG) homologs in the genome of the parasitiform tick *Hyalomma asiaticum*. Colors correspond to individual ALGs, with solid color indicating significant hits (p < 0.05) and transparent color indicating insignificant hits.

**Supplementary Figure S50.** Oxford dot plot showing the location of BCnS ancestral linkage group (ALG) homologs in the genome of the parasitiform tick *Ixodes scapularis*. Colors correspond to individual ALGs, with solid color indicating significant hits (p < 0.05) and transparent color indicating insignificant hits.

**Supplementary Figure S51.** Oxford dot plot showing the location of BCnS ancestral linkage group (ALG) homologs in the genome of the harvestman *Leiobunum manubriatum*. Colors correspond to individual ALGs, with solid color indicating significant hits (p < 0.05) and transparent color indicating insignificant hits.

**Supplementary Figure S52.** Oxford dot plot showing the location of BCnS ancestral linkage group (ALG) homologs in the genome of the horseshoe crab *Limulus polyphemus*. Colors correspond to individual ALGs, with solid color indicating significant hits (p < 0.05) and transparent color indicating insignificant hits.

**Supplementary Figure S53.** Oxford dot plot showing the location of BCnS ancestral linkage group (ALG) homologs in the genome of the mesothele spider *Liphistius suwat*. Colors correspond to individual ALGs, with solid color indicating significant hits (p < 0.05) and transparent color indicating insignificant hits.

**Supplementary Figure S54.** Oxford dot plot showing the location of BCnS ancestral linkage group (ALG) homologs in the genome of the vinegaroon *Mastigoproctus giganteus*. Colors correspond to individual ALGs, with solid color indicating significant hits (p < 0.05) and transparent color indicating insignificant hits.

**Supplementary Figure S55.** Oxford dot plot showing the location of BCnS ancestral linkage group (ALG) homologs in the genome of the harvestman *Odiellus spinosus*. Colors correspond to individual ALGs, with solid color indicating significant hits (p < 0.05) and transparent color indicating insignificant hits.

**Supplementary Figure S56.** Oxford dot plot showing the location of BCnS ancestral linkage group (ALG) homologs in the genome of the acariform mite *Oppia nitens*. Colors correspond to individual ALGs, with solid color indicating significant hits (p < 0.05) and transparent color indicating insignificant hits.

**Supplementary Figure S57.** Oxford dot plot showing the location of BCnS ancestral linkage group (ALG) homologs in the genome of the sea spider *Pycnogonum litorale*. Colors correspond to individual ALGs, with solid color indicating significant hits (p < 0.05) and transparent color indicating insignificant hits.

**Supplementary Figure S58.** Oxford dot plot showing the location of BCnS ancestral linkage group (ALG) homologs in the genome of the parasitiform tick *Rhipicephalus microplus*. Colors correspond to individual ALGs, with solid color indicating significant hits (p < 0.05) and transparent color indicating insignificant hits.

**Supplementary Figure S59.** Oxford dot plot showing the location of BCnS ancestral linkage group (ALG) homologs in the genome of the acariform mite *Sarcoptes scabiei*. Colors correspond to individual ALGs, with solid color indicating significant hits (p < 0.05) and transparent color indicating insignificant hits.

**Supplementary Figure S60.** Oxford dot plot showing the location of BCnS ancestral linkage group (ALG) homologs in the genome of the centipede *Strigamia acuminata*. Colors correspond to individual ALGs, with solid color indicating significant hits (p < 0.05) and transparent color indicating insignificant hits.

**Supplementary Figure S61.** Oxford dot plot showing the location of BCnS ancestral linkage group (ALG) homologs in the genome of the horseshoe crab *Tachypleus tridentatus*. Colors correspond to individual ALGs, with solid color indicating significant hits (p < 0.05) and transparent color indicating insignificant hits.

**Supplementary Figure S62.** Oxford dot plot showing the location of apulmonate ancestral linkage group (ALG) homologs in the genome of the parasitiform mite *Varroa destructor*. Colors correspond to individual ALGs, with solid color indicating significant hits (*p <* 0.05) and transparent color indicating insignificant hits.

**Supplementary Figure S63.** Mapping of ortholog gains and losses on unconstrained tree topology *versus* topology constrained to recover alternative placements for Acariformes.

**Supplementary Figure S64.** Mapping of ortholog gains and losses on topology constrained to recover Arachnida *versus* topology constrained to recover Xiphosura + Arachnopulmonata.

**Supplementary Figure S65.** Oxford dot plot showing the location of ancestral linkage group (ALG) homologs in the genomes of three mites for the apulmonate (top row) and the BCnS (bottom row) linkage groups. Left column: *Phytoseiulus persimilis* (Parasitiformes). Middle: *Tetranychus urticae* (Acariformes). Right: Eriophyidae gen. sp. (Acariformes). Colors correspond to individual ALGs, with solid color indicating significant hits (*p <* 0.05) and transparent color indicating insignificant hits.

**Supplementary Table S1.** Summary statistics for newly generated genomic resources.

**Supplementary Table S2.** Accession data for new and published genome assemblies included in analyses.

**Supplementary Table S3.** Annotations and genomic locations of Hox, HRO, Irx, SINE, and Nk gene cluster homologs in the genome of the mygalomorph spider *Aphonopelma hentzi*.

**Supplementary Table S4.** Annotations and genomic locations of Hox, HRO, Irx, SINE, and Nk gene cluster homologs in the genome of the araneomorph spider *Argiope bruennichi*.

**Supplementary Table S5.** Annotations and genomic locations of Hox, HRO, Irx, SINE, and Nk gene cluster homologs in the genome of the acariform mite *Blomia tropicalis*.

**Supplementary Table S6.** Annotations and genomic locations of Hox, HRO, Irx, SINE, and Nk gene cluster homologs in the genome of the horseshoe crab *Carcinoscorpius rotundicauda*.

**Supplementary Table S7.** Annotations and genomic locations of Hox, HRO, Irx, SINE, and Nk gene cluster homologs in the genome of the scorpion *Centruroides sculpturatus*.

**Supplementary Table S8.** Annotations and genomic locations of Hox, HRO, Irx, SINE, and Nk gene cluster homologs in the genome of the pseudoscorpion *Cordylochernes scorpioides*.

**Supplementary Table S9.** Annotations and genomic locations of Hox, HRO, Irx, SINE, and Nk gene cluster homologs in the genome of the hooded tick spider *Cryptocellus* cf. *goodnighti*.

**Supplementary Table S10.** Annotations and genomic locations of Hox, HRO, Irx, SINE, and Nk gene cluster homologs in the genome of the parasitiform tick *Dermacentor silvarum*.

**Supplementary Table S11.** Annotations and genomic locations of Hox, HRO, Irx, SINE, and Nk gene cluster homologs in the genome of the solifuge *Gluvia dorsalis*.

**Supplementary Table S12.** Annotations and genomic locations of Hox, HRO, Irx, SINE, and Nk gene cluster homologs in the genome of the parasitiform tick *Haemaphysalis longicornis*.

**Supplementary Table S13.** Annotations and genomic locations of Hox, HRO, Irx, SINE, and Nk gene cluster homologs in the genome of the acariform mite *Halotydeus destructor*.

**Supplementary Table S14.** Annotations and genomic locations of Hox, HRO, Irx, SINE, and Nk gene cluster homologs in the genome of the parasitiform tick *Hyalomma asiaticum*.

**Supplementary Table S15.** Annotations and genomic locations of Hox, HRO, Irx, SINE, and Nk gene cluster homologs in the genome of the horseshoe crab *Limulus polyphemus*.

**Supplementary Table S16.** Annotations and genomic locations of Hox, HRO, Irx, SINE, and Nk gene cluster homologs in the genome of the Mesothelae spider *Liphistius suwat*.

**Supplementary Table S17.** Annotations and genomic locations of Hox, HRO, Irx, SINE, and Nk gene cluster homologs in the genome of the vinegaroon *Mastigoproctus giganteus*.

**Supplementary Table S18.** Annotations and genomic locations of Hox, HRO, Irx, SINE, and Nk gene cluster homologs in the genome of the harvestman *Odiellus spinosus*.

**Supplementary Table S19.** Annotations and genomic locations of Hox, HRO, Irx, SINE, and Nk gene cluster homologs in the genome of the parasitiform mite *Phytoseiulus persimilis*.

**Supplementary Table S20.** Annotations and genomic locations of Hox, HRO, Irx, SINE, and Nk gene cluster homologs in the genome of the palpigrade *Prokoenenia wheeleri*.

**Supplementary Table S21.** Annotations and genomic locations of Hox, HRO, Irx, SINE, and Nk gene cluster homologs in the genome of the sea spider *Pycnogonum litorale*.

**Supplementary Table S22.** Annotations and genomic locations of Hox, HRO, Irx, SINE, and Nk gene cluster homologs in the genome of the parasitiform tick *Rhipicephalus microplus*.

**Supplementary Table S23.** Annotations and genomic locations of Hox, HRO, Irx, SINE, and Nk gene cluster homologs in the genome of the acariform mite *Sarcoptes scabiei*.

**Supplementary Table S24.** Annotations and genomic locations of Hox, HRO, Irx, SINE, and Nk gene cluster homologs in the genome of the centipede *Strigamia acuminata*.

**Supplementary Table S25.** Annotations and genomic locations of Hox, HRO, Irx, SINE, and Nk gene cluster homologs in the genome of the horseshoe crab *Tachypleus tridentatus*.

**Supplementary Table S26.** Counts of microRNAs recovered in arthropod genomes using the MirMachine pipeline.

**Supplementary Table S27.** Results of Approximately Unbiased and Shimodaira-Hasegawa tests for unconstrained PMSF tree topology *versus* trees constrained to recover conflicting hypotheses of arachnid relationships.

**Supplementary Table S28.** Counts of ortholog gains and losses for competing tree topologies.

