## Supplementary Figures S1-S65 for "No silver bullet: Patterns of macrosynteny recapitulate systemic conflicts in the higher-level relationships of the arachnids"

**Supplementary Figure S1.** Snail plots and HiC contact maps for newly sequenced arachnid genomes. Numbers below contact maps indicate number of large pseudomolecules and percentage of genome sequence mapped to large pseudomolecules.

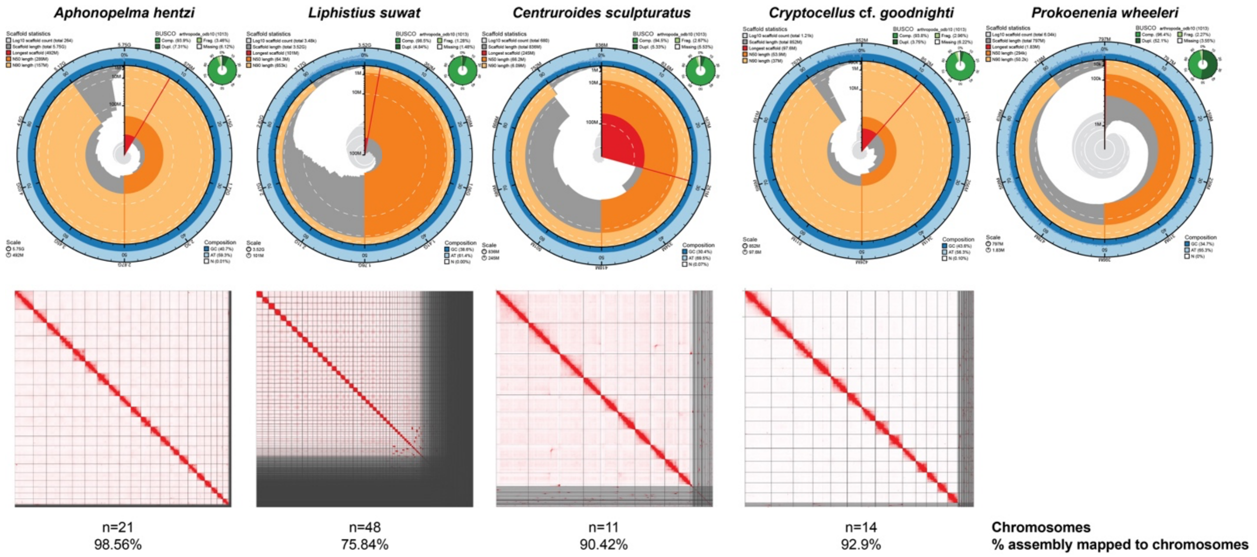

**Supplementary Figure S2.** Architecture of evolutionarily conserved gene clusters in genomes of Arachnopulmonata.

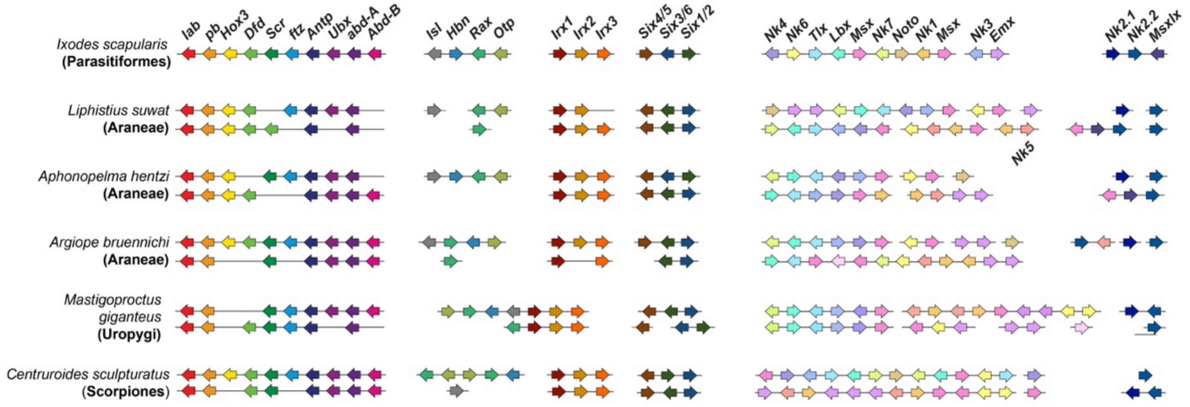

**Supplementary Figure S3.** Architecture of evolutionarily conserved gene clusters in genomes of apulmonate arachnids. Note the atomization of the Hox cluster in the fast-evolving Mesostigmata genome.

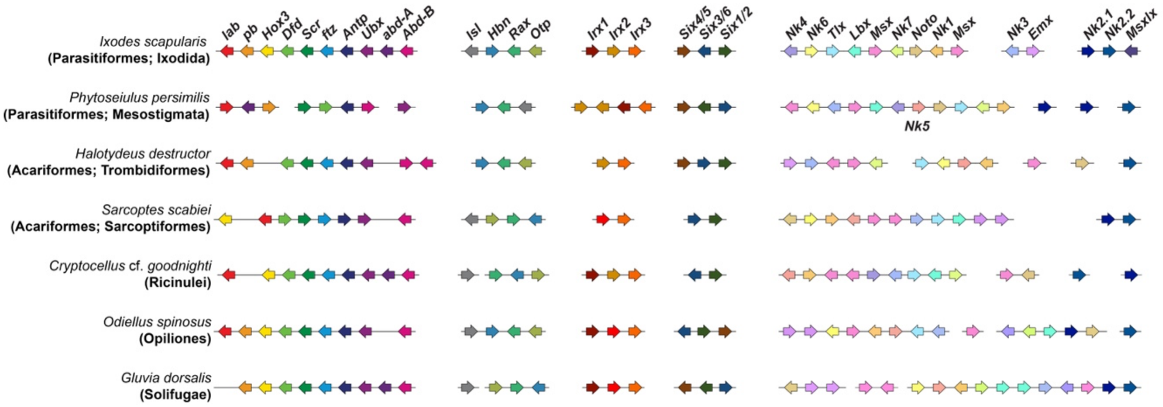

**Supplementary Figure S4.** Maximum likelihood tree topology of Hox cluster genes.

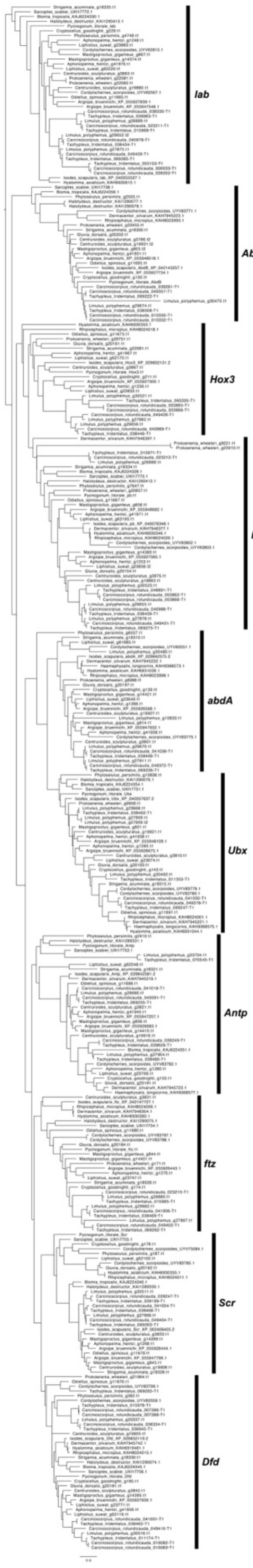

**Supplementary Figure S5.** Maximum likelihood tree topology of HRO cluster genes.

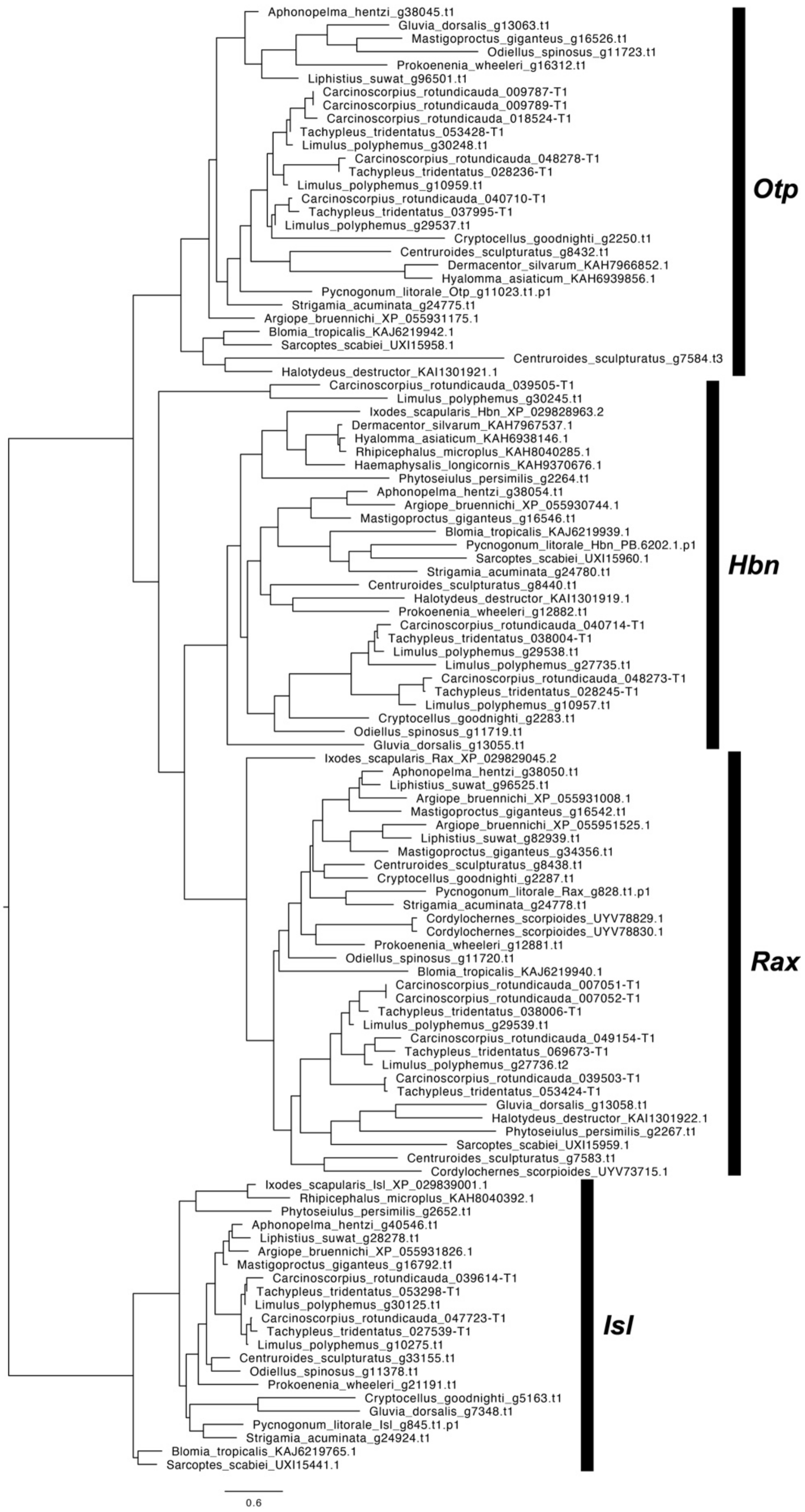

**Supplementary Figure S6.** Maximum likelihood tree topology of Irx cluster genes.

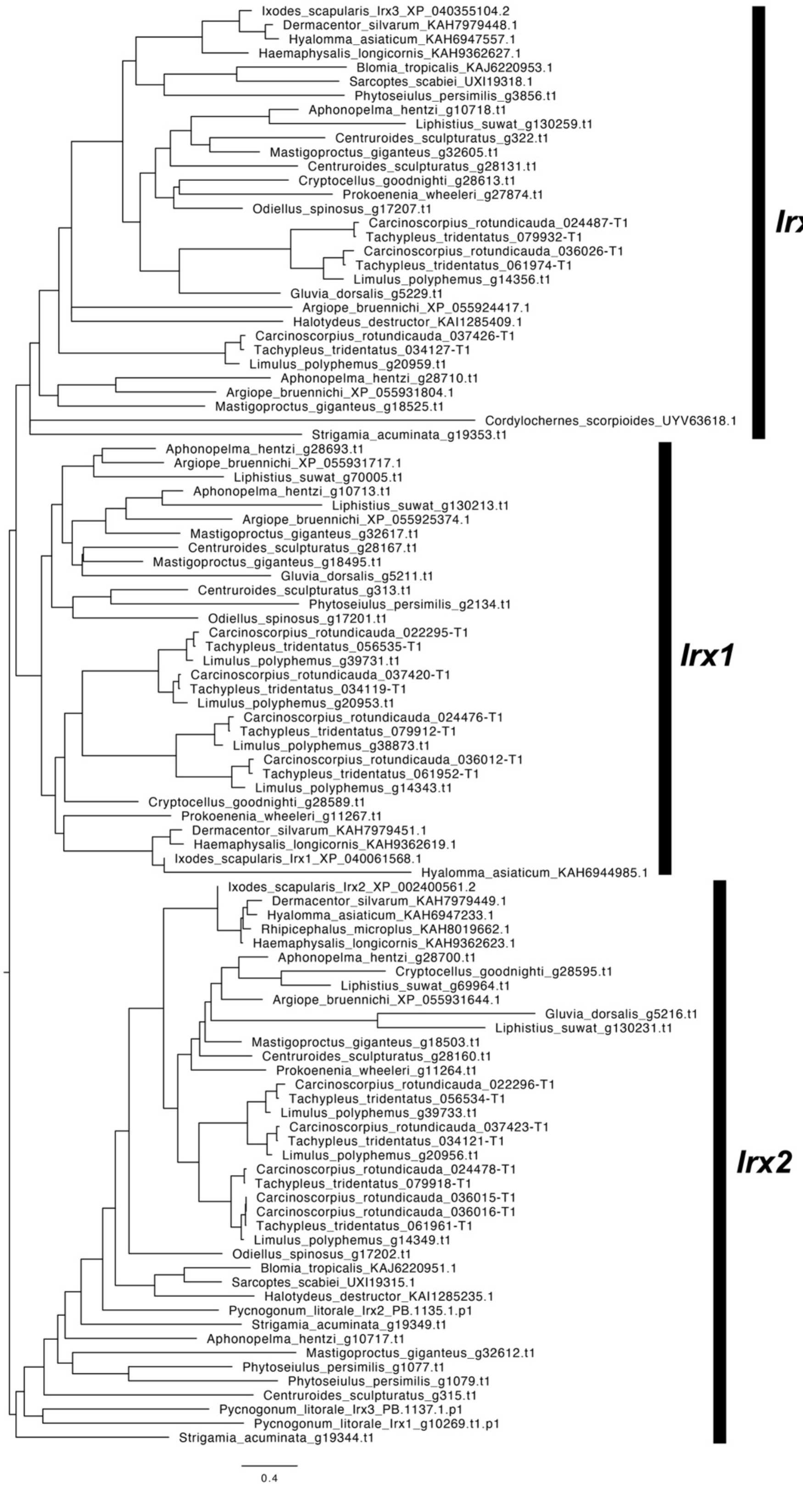

**Supplementary Figure S7.** Maximum likelihood tree topology of SINE cluster genes.

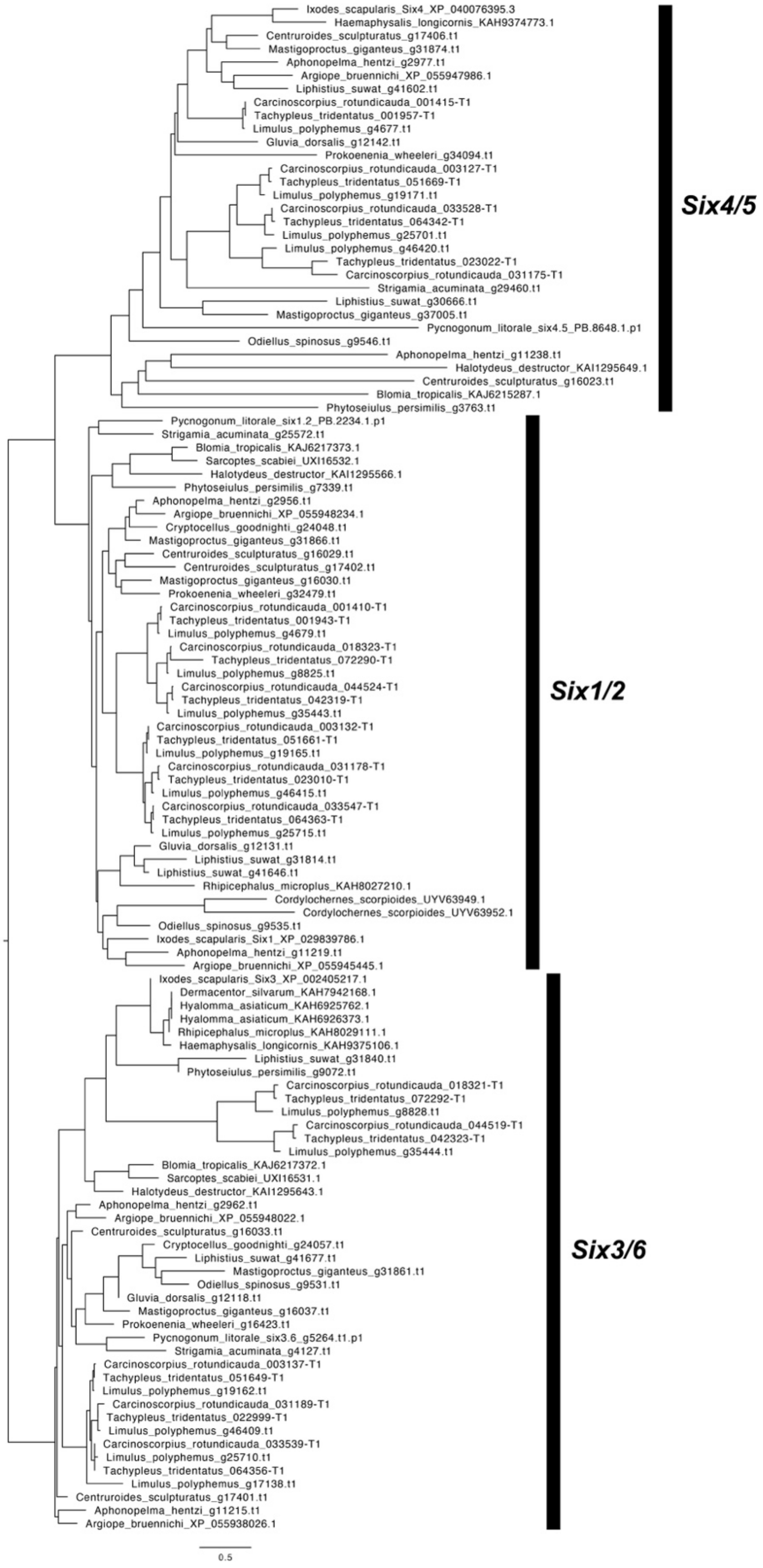

**Supplementary Figure S8.** Maximum likelihood tree topology of Nk cluster genes.

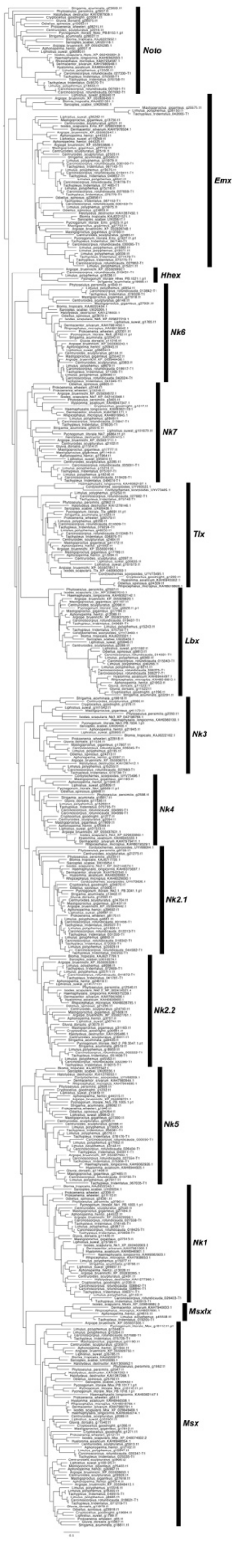

**Supplementary Figure S9.** Oxford dot plot showing the location of apulmonate ancestral linkage group (ALG) homologs in the genome of the mygalomorph spider *Aphonopelma hentzi*. Colors correspond to individual ALGs, with solid color indicating significant hits ($p < 0.05$) and transparent color indicating insignificant hits.

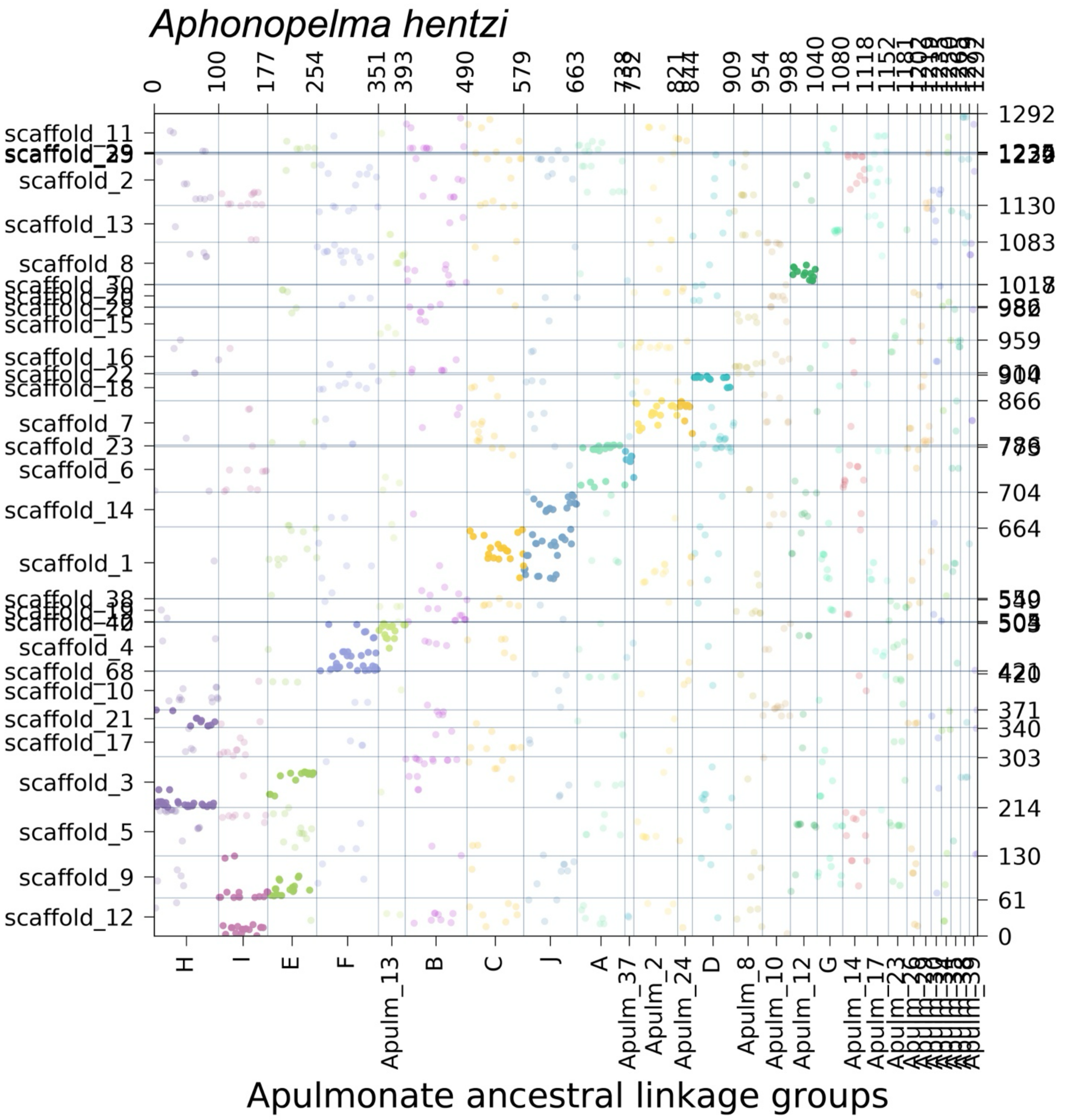

**Supplementary Figure S10.** Oxford dot plot showing the location of apulmonate ancestral linkage group (ALG) homologs in the genome of the araneomorph spider *Argiope bruennichi*. Colors correspond to individual ALGs, with solid color indicating significant hits ($p < 0.05$) and transparent color indicating insignificant hits.

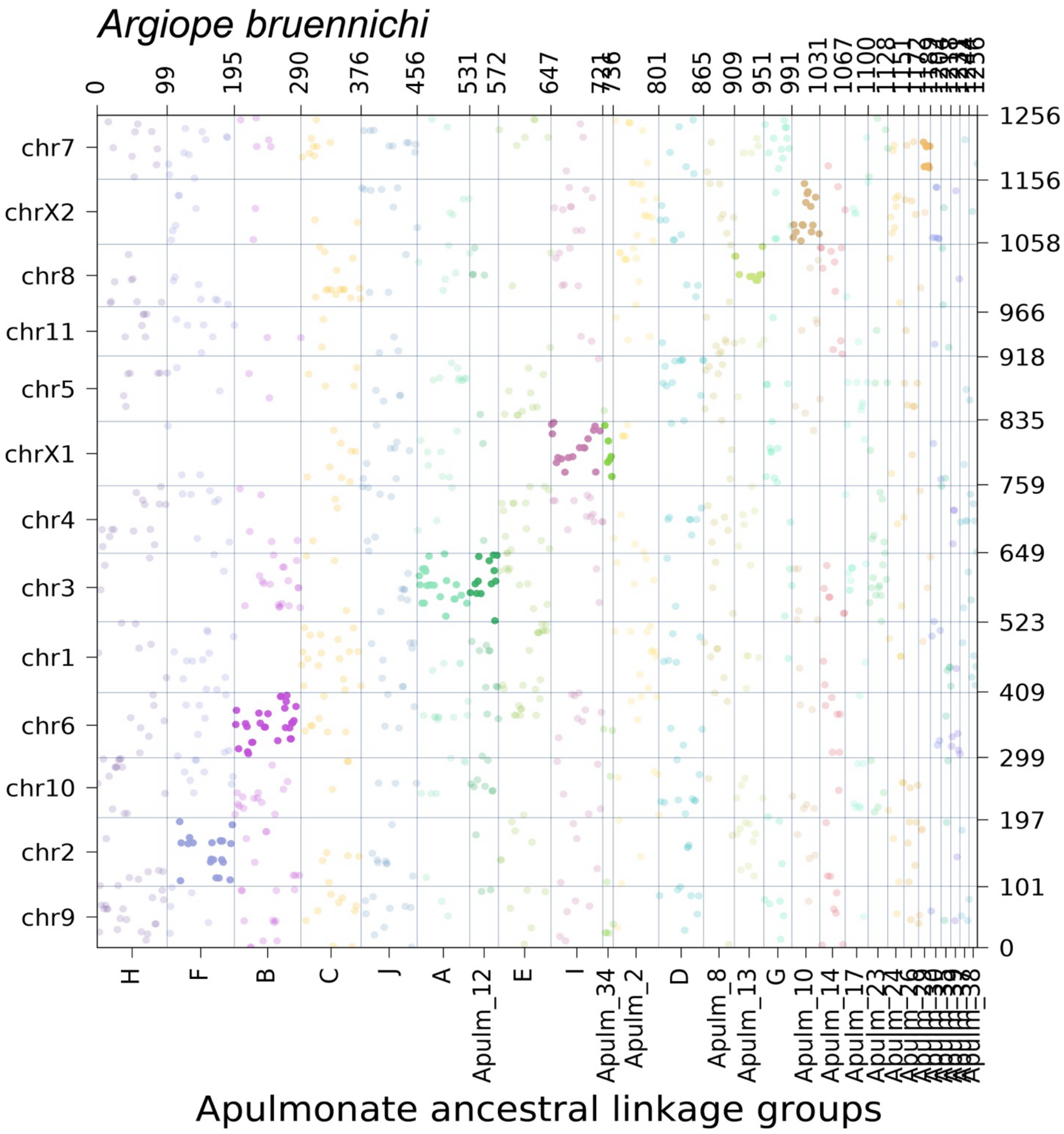

**Supplementary Figure S11.** Oxford dot plot showing the location of apulmonate ancestral linkage group (ALG) homologs in the genome of the acariform mite *Blomia tropicalis*. Colors correspond to individual ALGs, with solid color indicating significant hits ($p < 0.05$) and transparent color indicating insignificant hits.

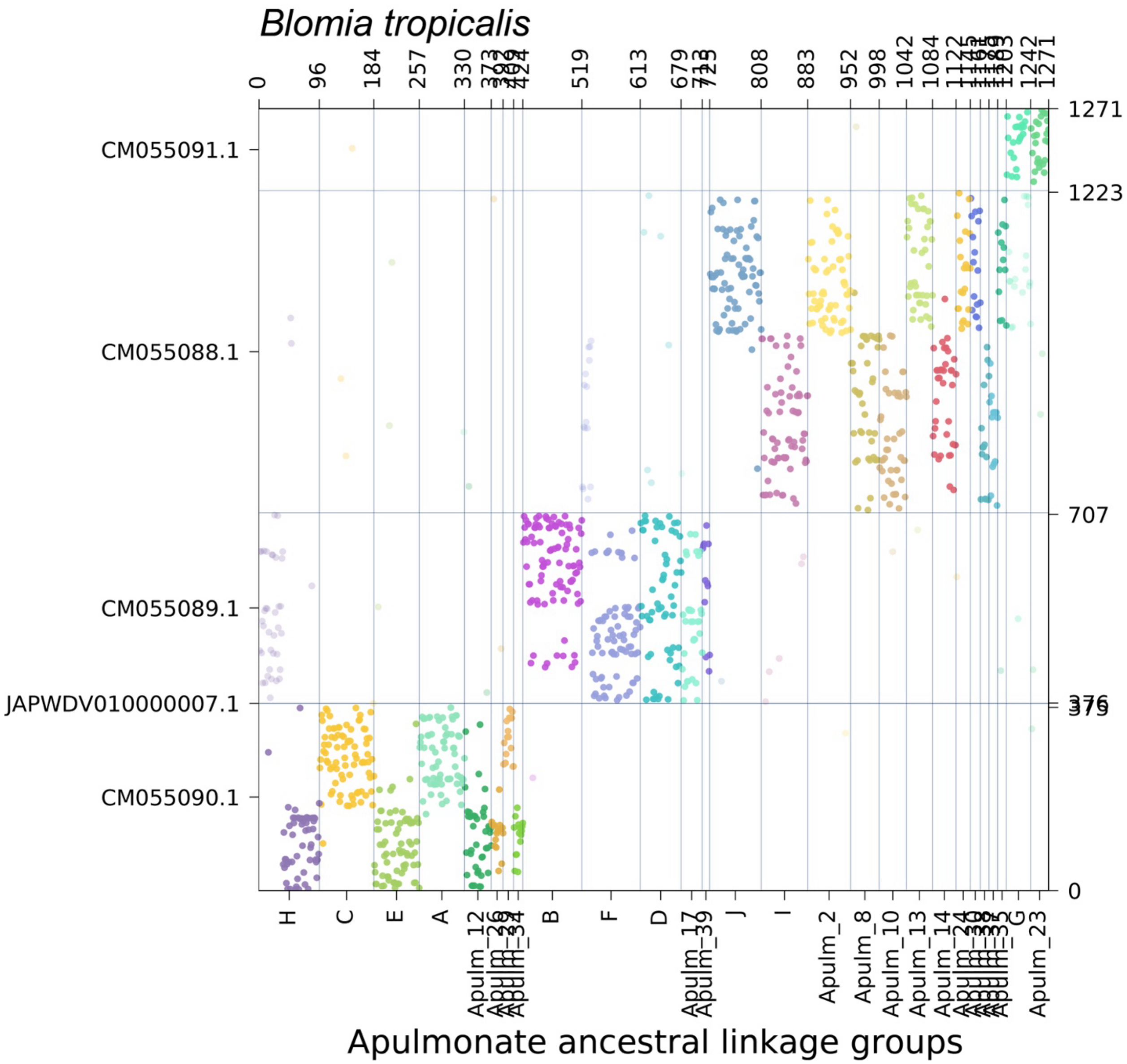

**Supplementary Figure S12.** Oxford dot plot showing the location of apulmonate ancestral linkage group (ALG) homologs in the genome of the horseshoe crab *Carcinoscorpius rotundicauda*. Colors correspond to individual ALGs, with solid color indicating significant hits ($p < 0.05$) and transparent color indicating insignificant hits.

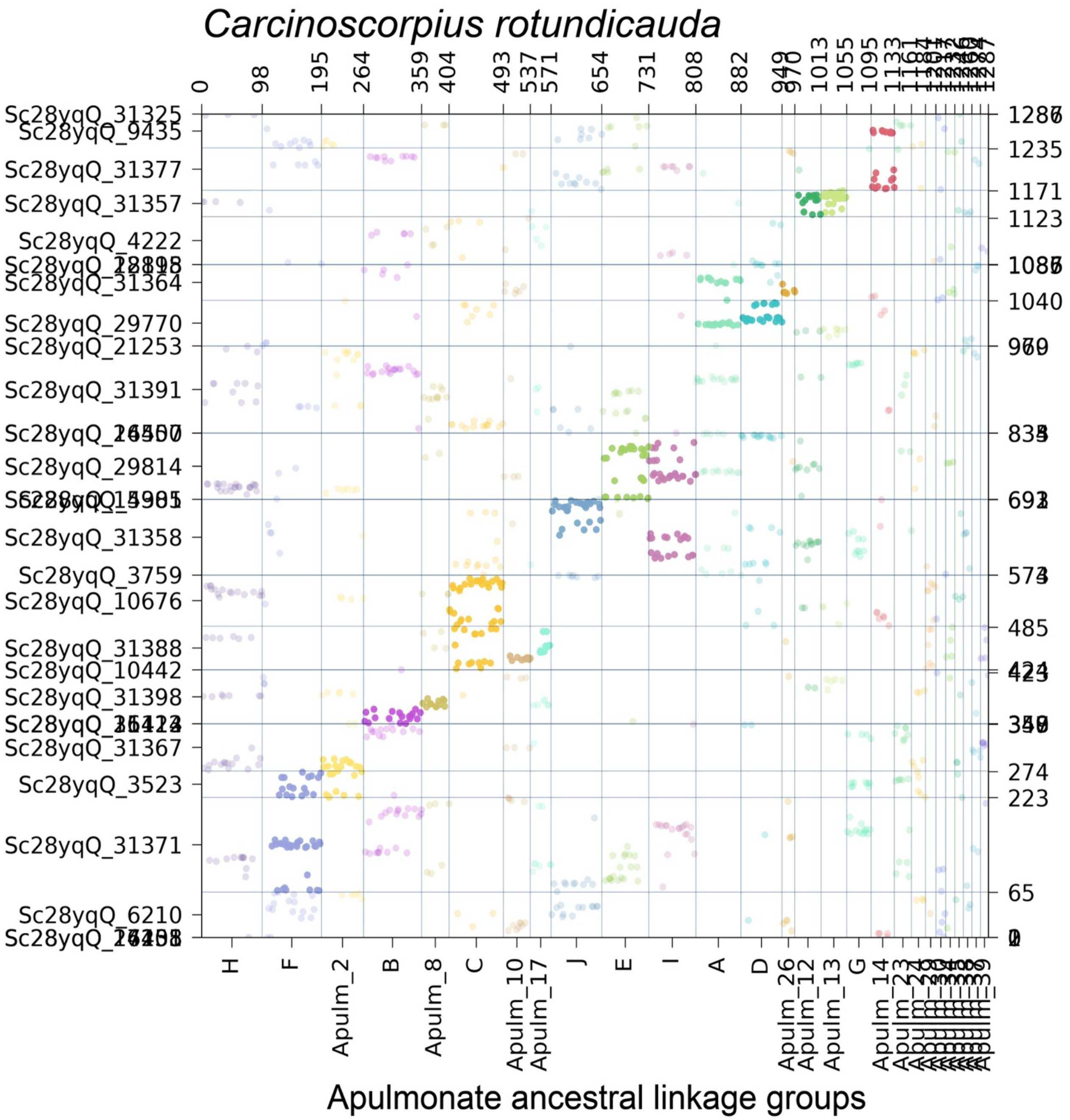

**Supplementary Figure S13.** Oxford dot plot showing the location of apulmonate ancestral linkage group (ALG) homologs in the genome of the scorpion *Centruroides sculpturatus*. Colors correspond to individual ALGs, with solid color indicating significant hits ($p < 0.05$) and transparent color indicating insignificant hits.

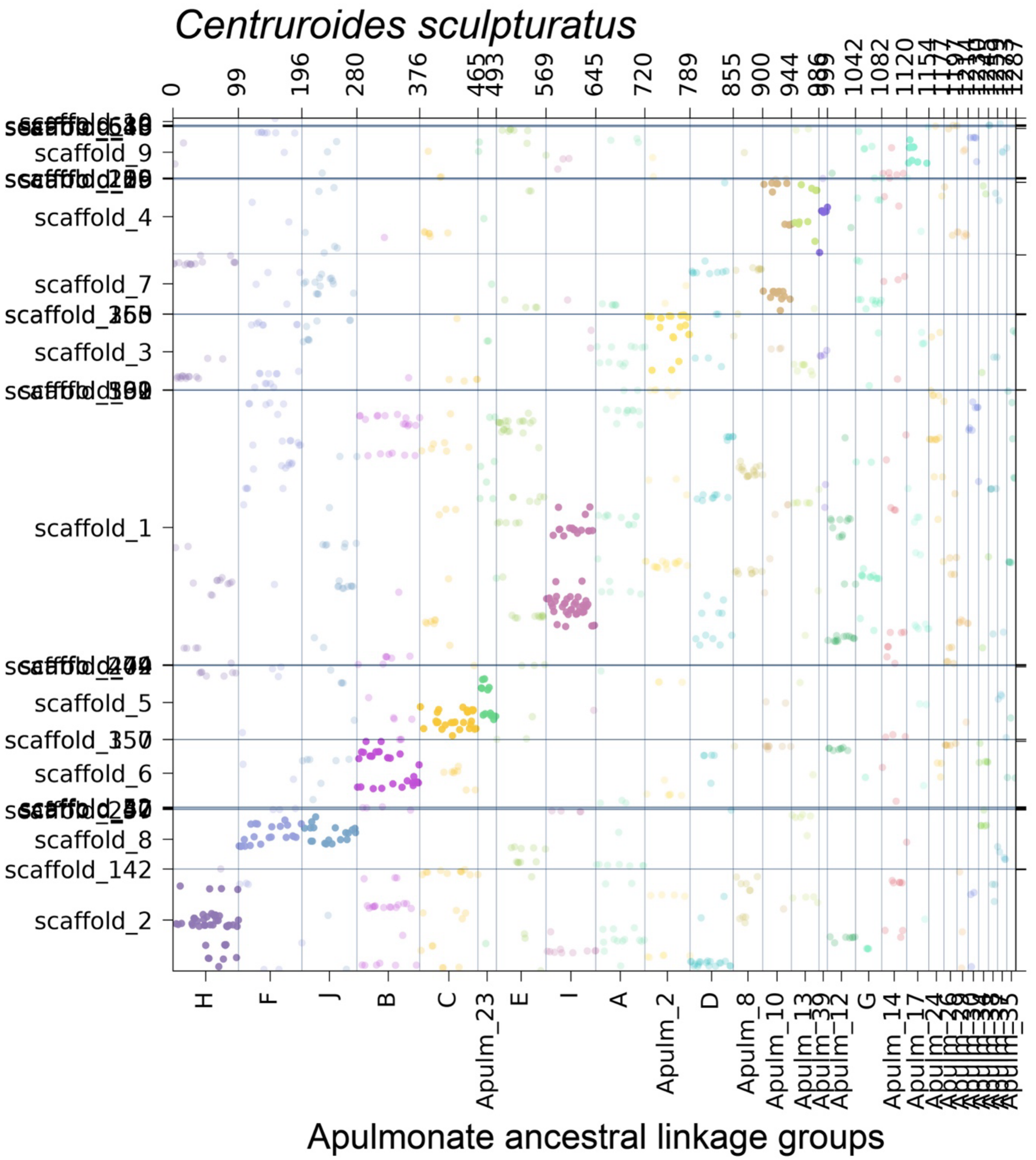

**Supplementary Figure S14.** Oxford dot plot showing the location of apulmonate ancestral linkage group (ALG) homologs in the genome of the pseudoscorpion *Cordylochernes scorpioides*. Colors correspond to individual ALGs, with solid color indicating significant hits ($p < 0.05$) and transparent color indicating insignificant hits.

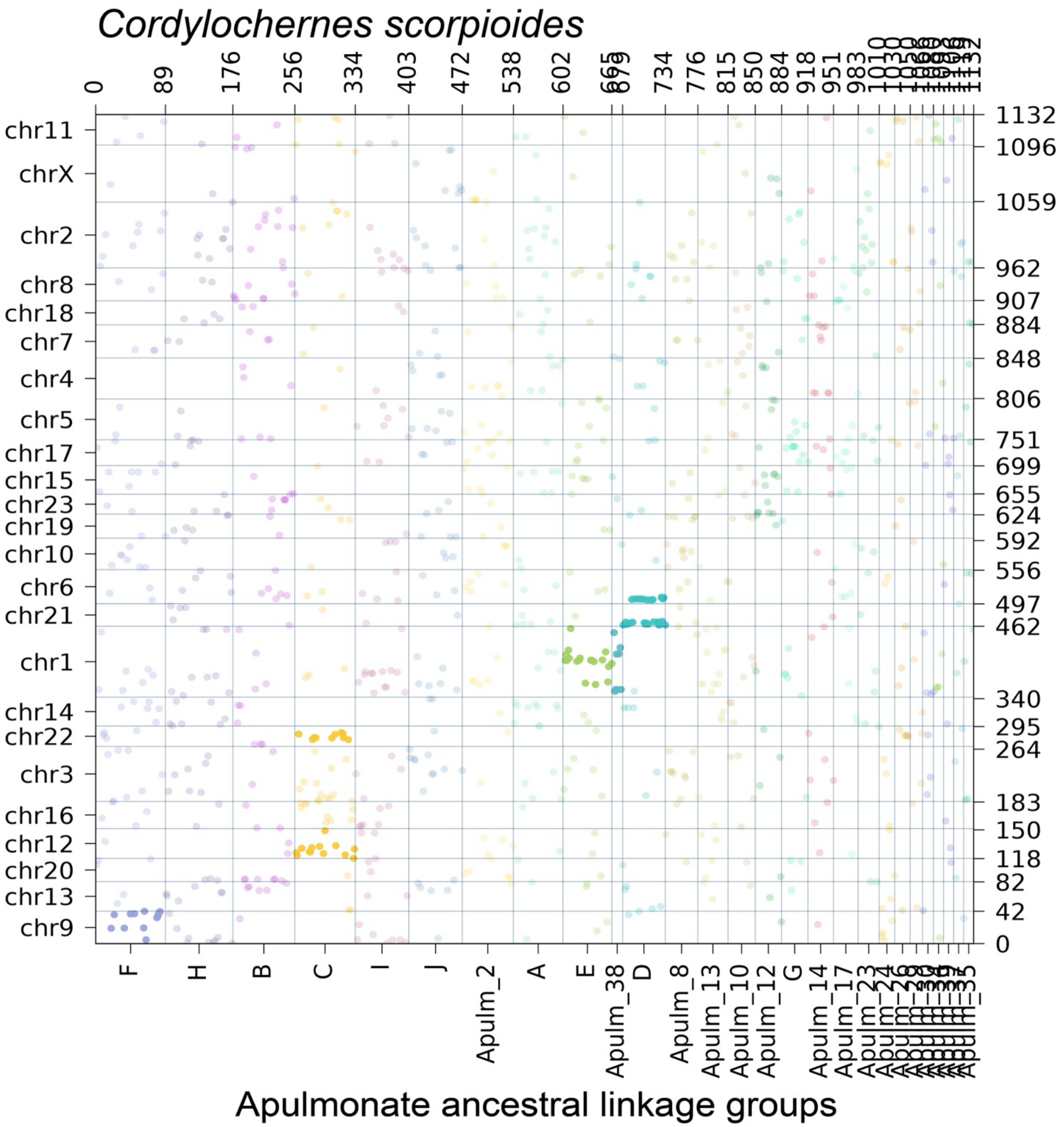

**Supplementary Figure S15.** Oxford dot plot showing the location of apulmonate ancestral linkage group (ALG) homologs in the genome of the ricinuleid *Cryptocellus* cf. *goodnighti*. Colors correspond to individual ALGs, with solid color indicating significant hits ($p < 0.05$) and transparent color indicating insignificant hits.

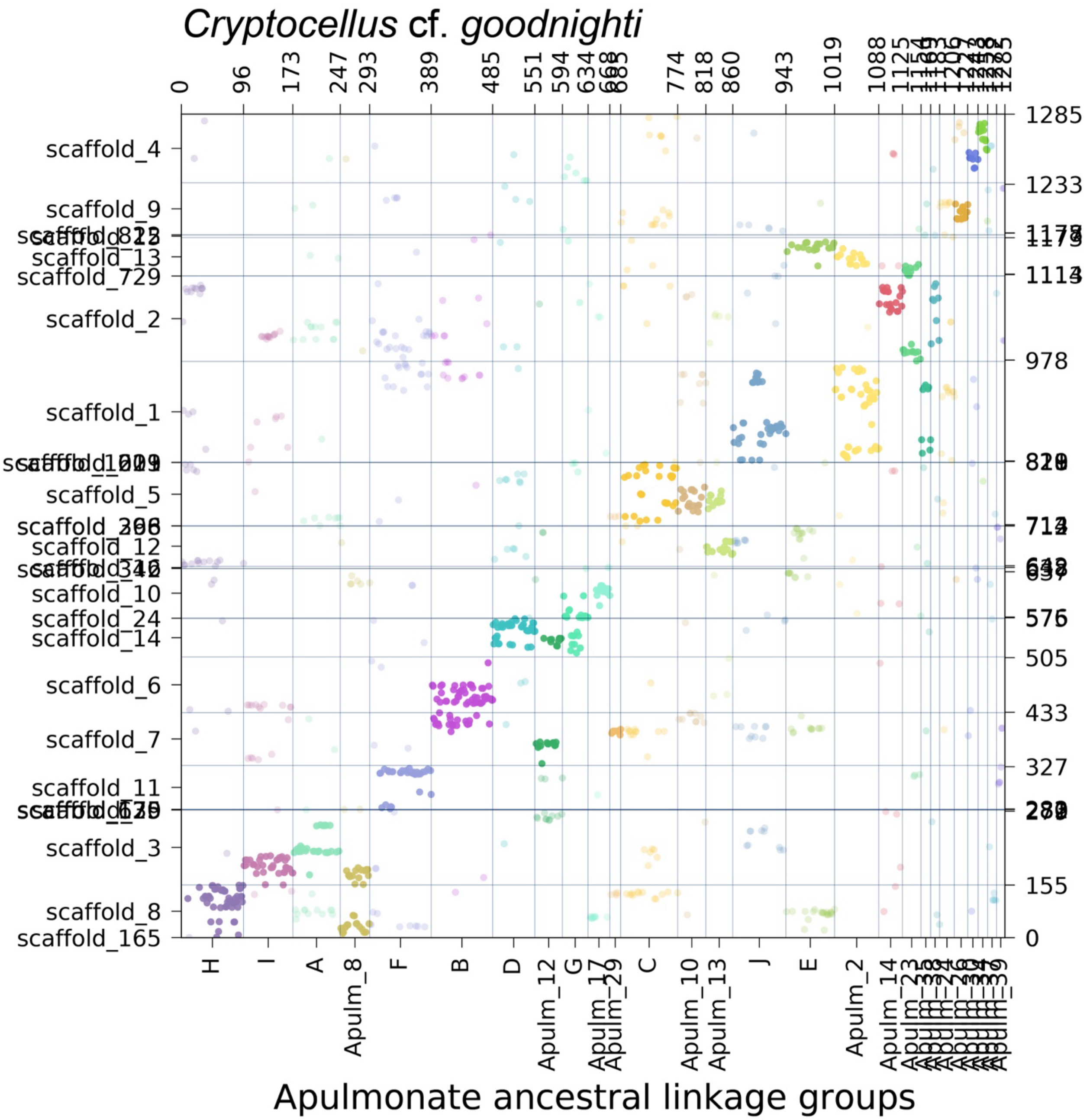

**Supplementary Figure S16.** Oxford dot plot showing the location of apulmonate ancestral linkage group (ALG) homologs in the genome of the parasitiform tick *Dermacentor silvarum*. Colors correspond to individual ALGs, with solid color indicating significant hits ($p < 0.05$) and transparent color indicating insignificant hits.

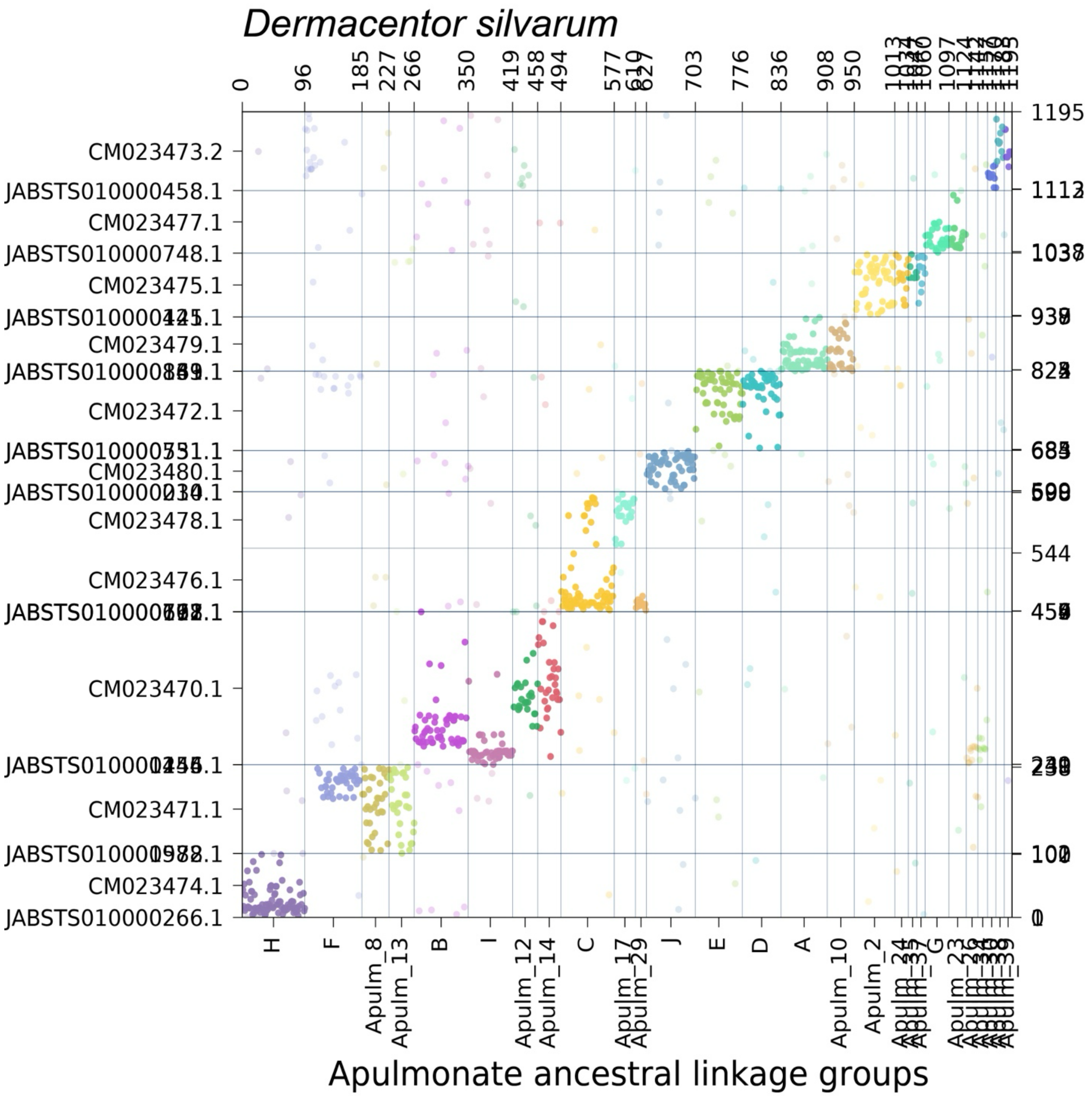

**Supplementary Figure S17.** Oxford dot plot showing the location of apulmonate ancestral linkage group (ALG) homologs in the genome of the solifuge *Gluvia dorsalis*. Colors correspond to individual ALGs, with solid color indicating significant hits ($p < 0.05$) and transparent color indicating insignificant hits.

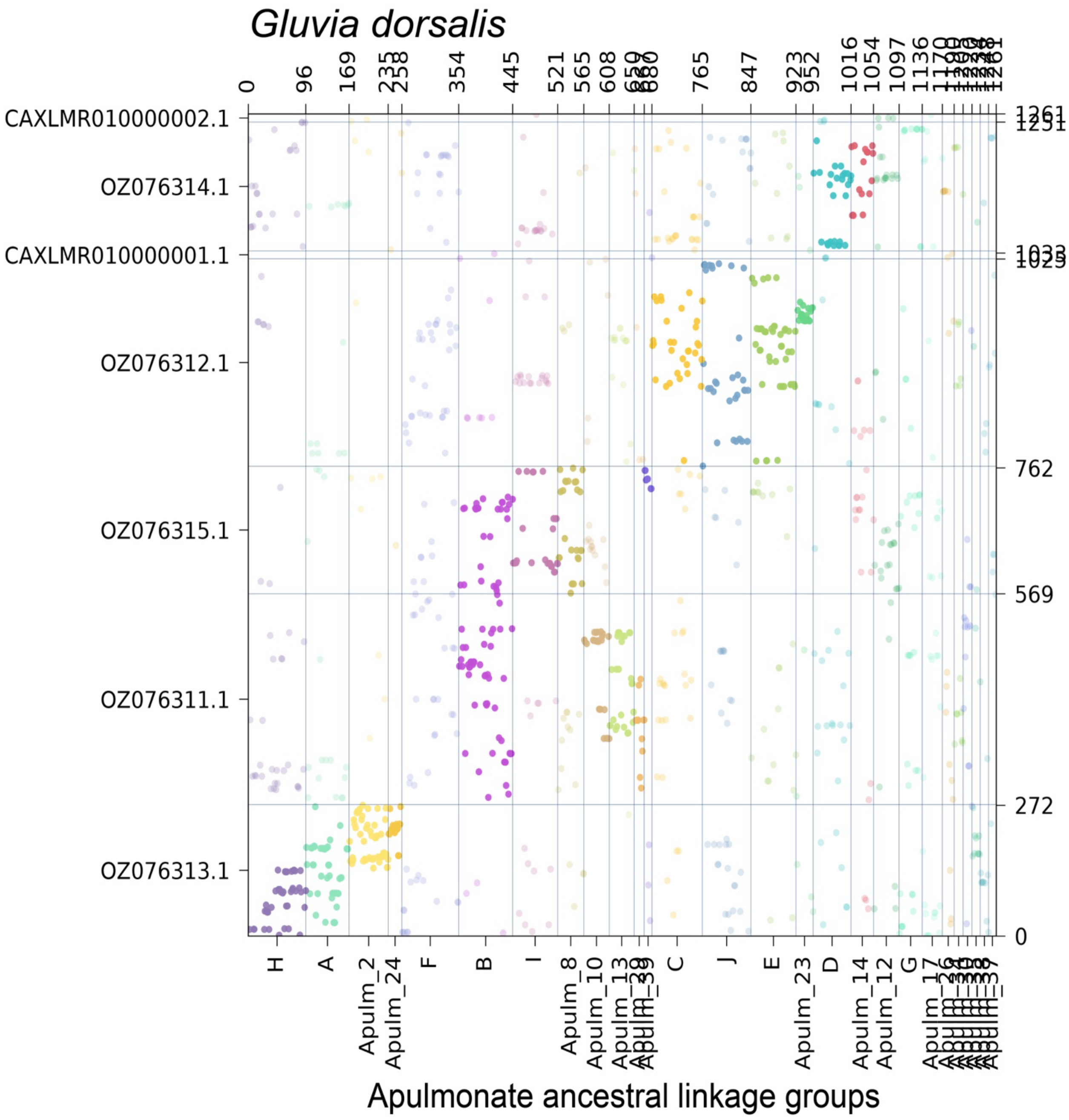

**Supplementary Figure S18.** Oxford dot plot showing the location of apulmonate ancestral linkage group (ALG) homologs in the genome of the scorpion *Hadrurus arizonensis*. Colors correspond to individual ALGs, with solid color indicating significant hits ($p < 0.05$) and transparent color indicating insignificant hits.

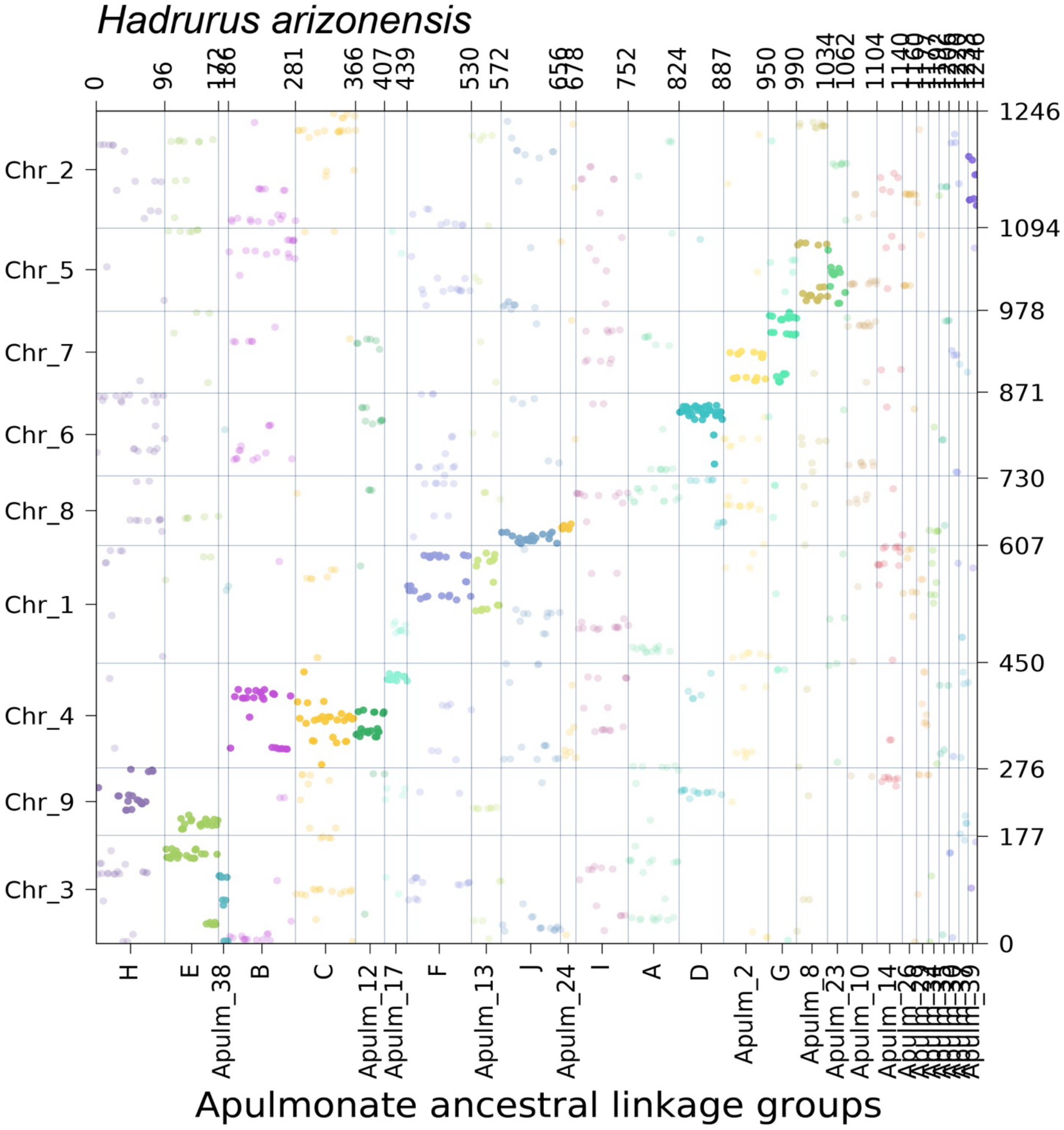

**Supplementary Figure S19.** Oxford dot plot showing the location of apulmonate ancestral linkage group (ALG) homologs in the genome of the parasitiform tick *Haemaphysalis longicornis*. Colors correspond to individual ALGs, with solid color indicating significant hits ($p < 0.05$) and transparent color indicating insignificant hits.

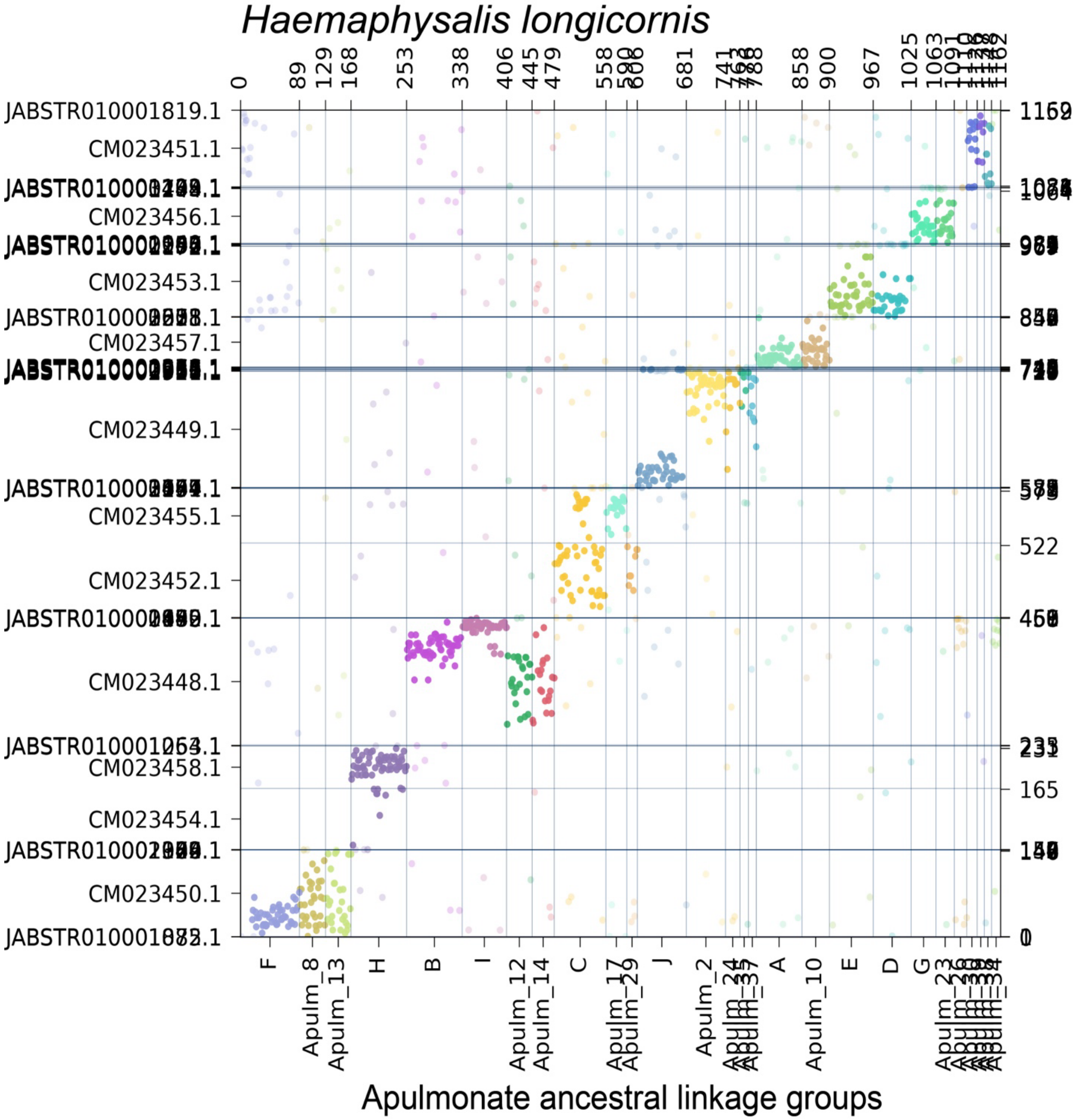

**Supplementary Figure S20.** Oxford dot plot showing the location of apulmonate ancestral linkage group (ALG) homologs in the genome of the acariform mite *Halotydeus destructor*. Colors correspond to individual ALGs, with solid color indicating significant hits ($p < 0.05$) and transparent color indicating insignificant hits.

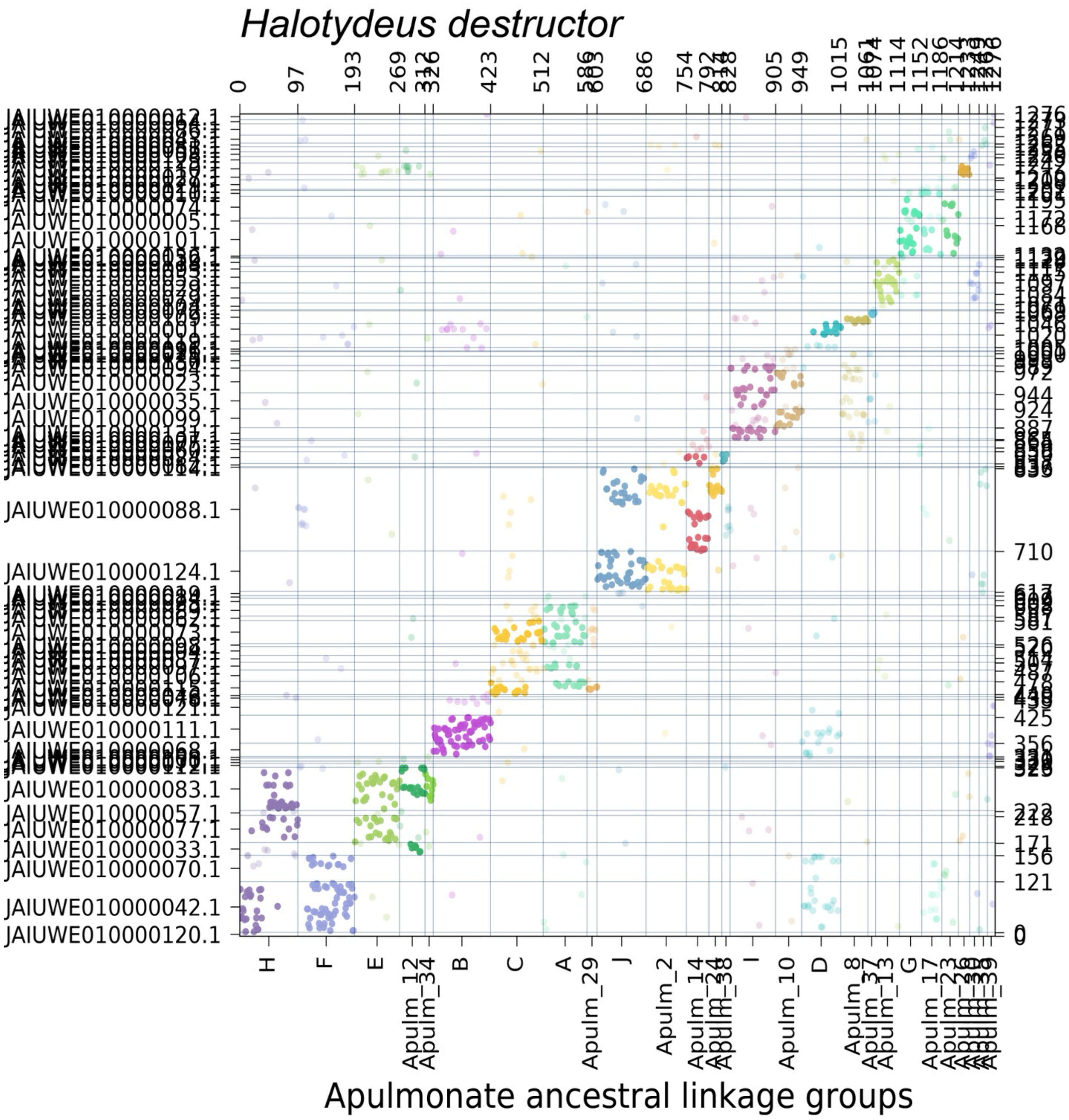

**Supplementary Figure S21.** Oxford dot plot showing the location of apulmonate ancestral linkage group (ALG) homologs in the genome of the parasitiform tick *Hyalomma asiaticum*. Colors correspond to individual ALGs, with solid color indicating significant hits ($p < 0.05$) and transparent color indicating insignificant hits.

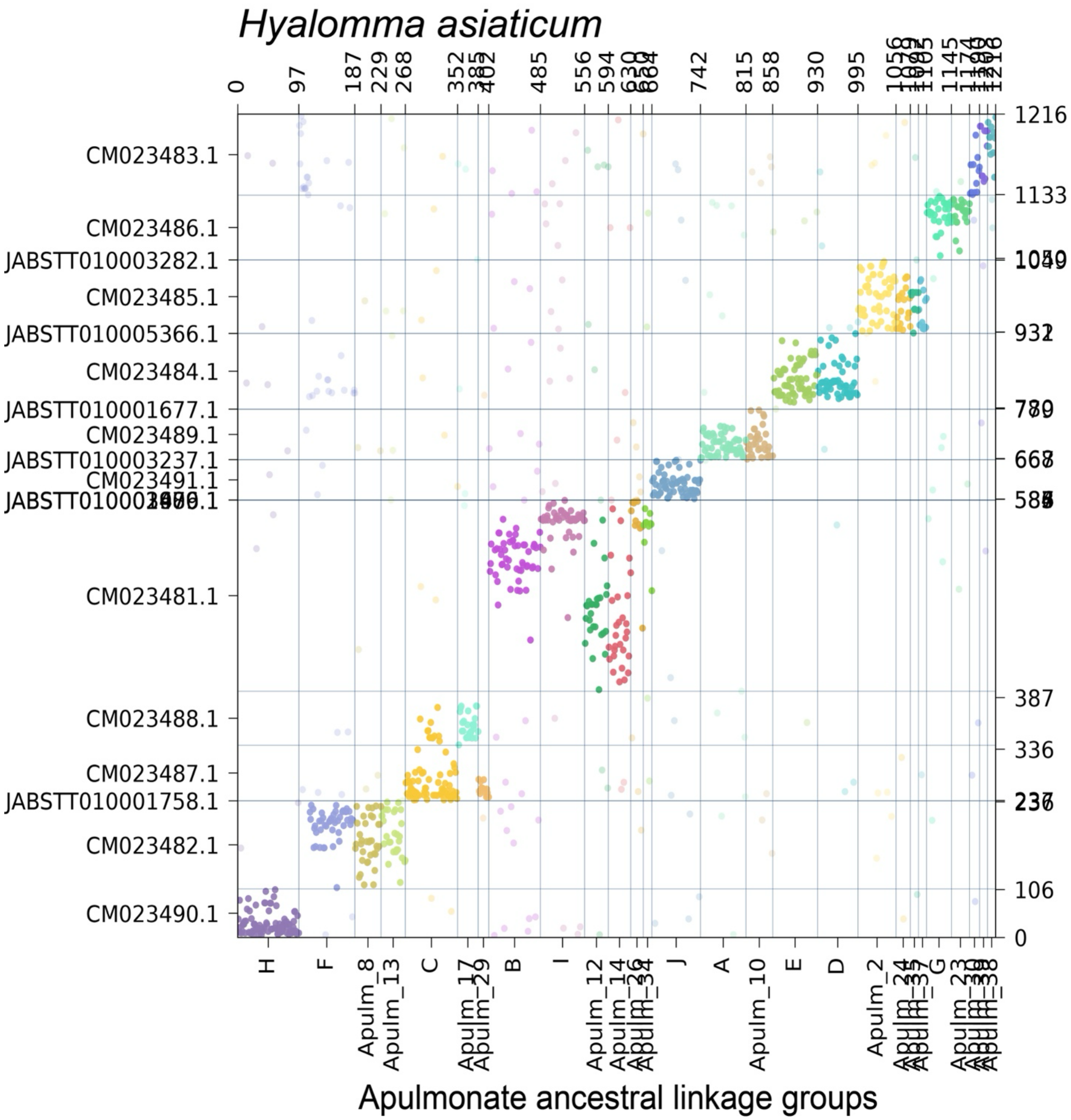

**Supplementary Figure S22.** Oxford dot plot showing the location of apulmonate ancestral linkage group (ALG) homologs in the genome of the parasitiform tick *Ixodes scapularis*. Colors correspond to individual ALGs, with solid color indicating significant hits ($p < 0.05$) and transparent color indicating insignificant hits.

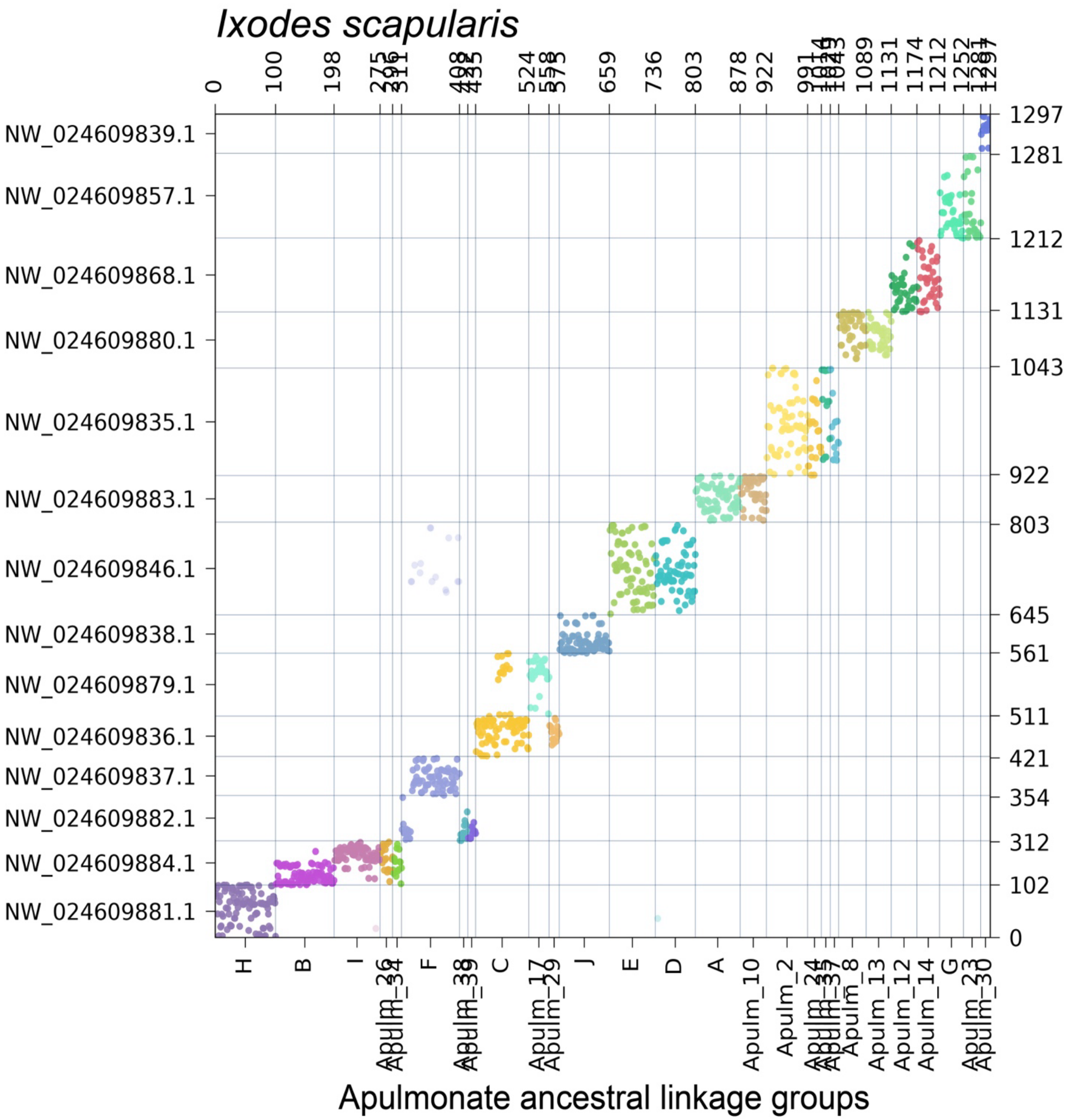

**Supplementary Figure S23.** Oxford dot plot showing the location of apulmonate ancestral linkage group (ALG) homologs in the genome of the harvestman *Leiobunum manubriatum*. Colors correspond to individual ALGs, with solid color indicating significant hits ($p < 0.05$) and transparent color indicating insignificant hits.

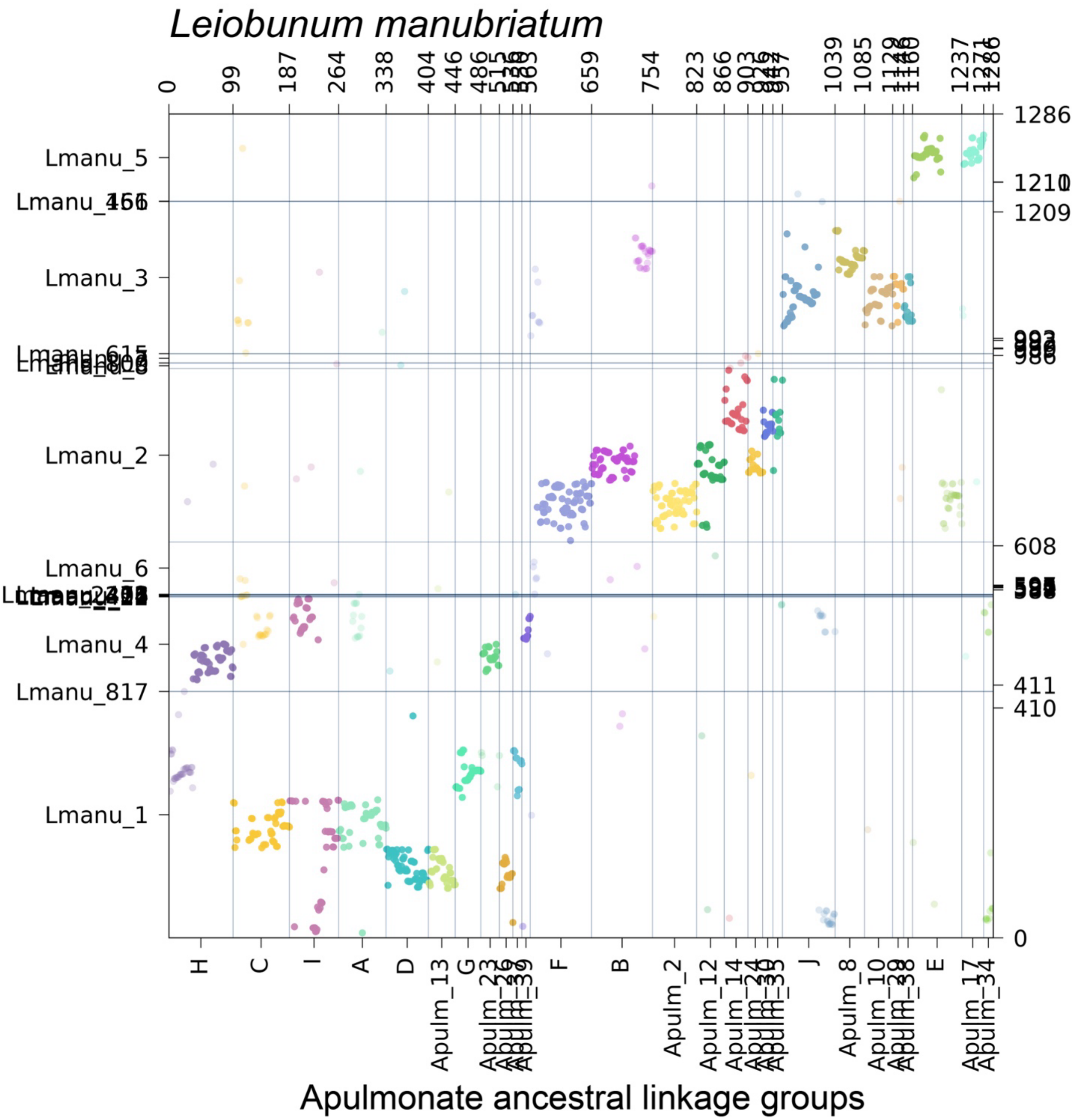

**Supplementary Figure S24.** Oxford dot plot showing the location of apulmonate ancestral linkage group (ALG) homologs in the genome of the horseshoe crab *Limulus polyphemus*. Colors correspond to individual ALGs, with solid color indicating significant hits ($p < 0.05$) and transparent color indicating insignificant hits.

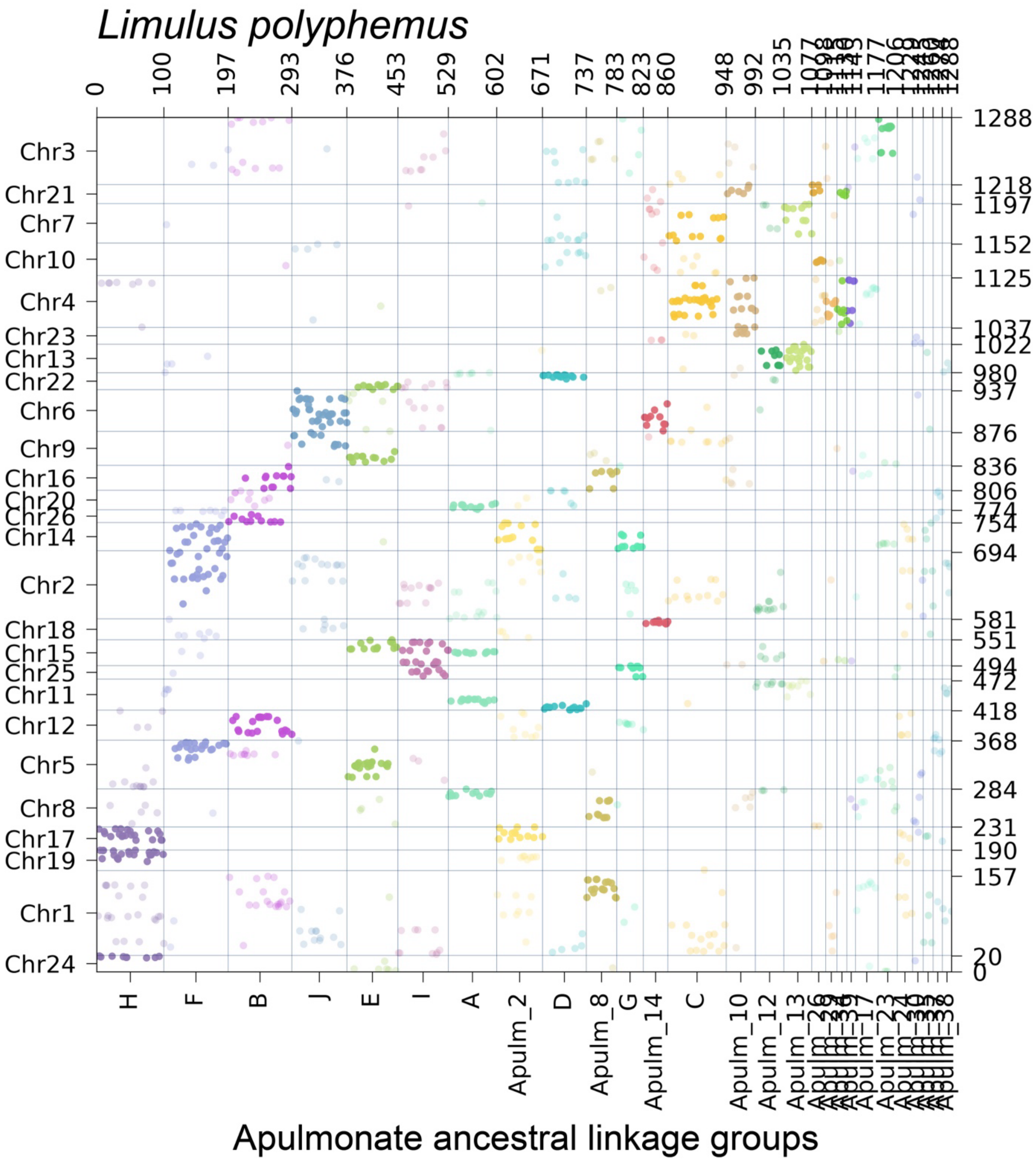

**Supplementary Figure S25.** Oxford dot plot showing the location of apulmonate ancestral linkage group (ALG) homologs in the genome of the mesothele spider *Liphistius suwat*. Colors correspond to individual ALGs, with solid color indicating significant hits ($p < 0.05$) and transparent color indicating insignificant hits.

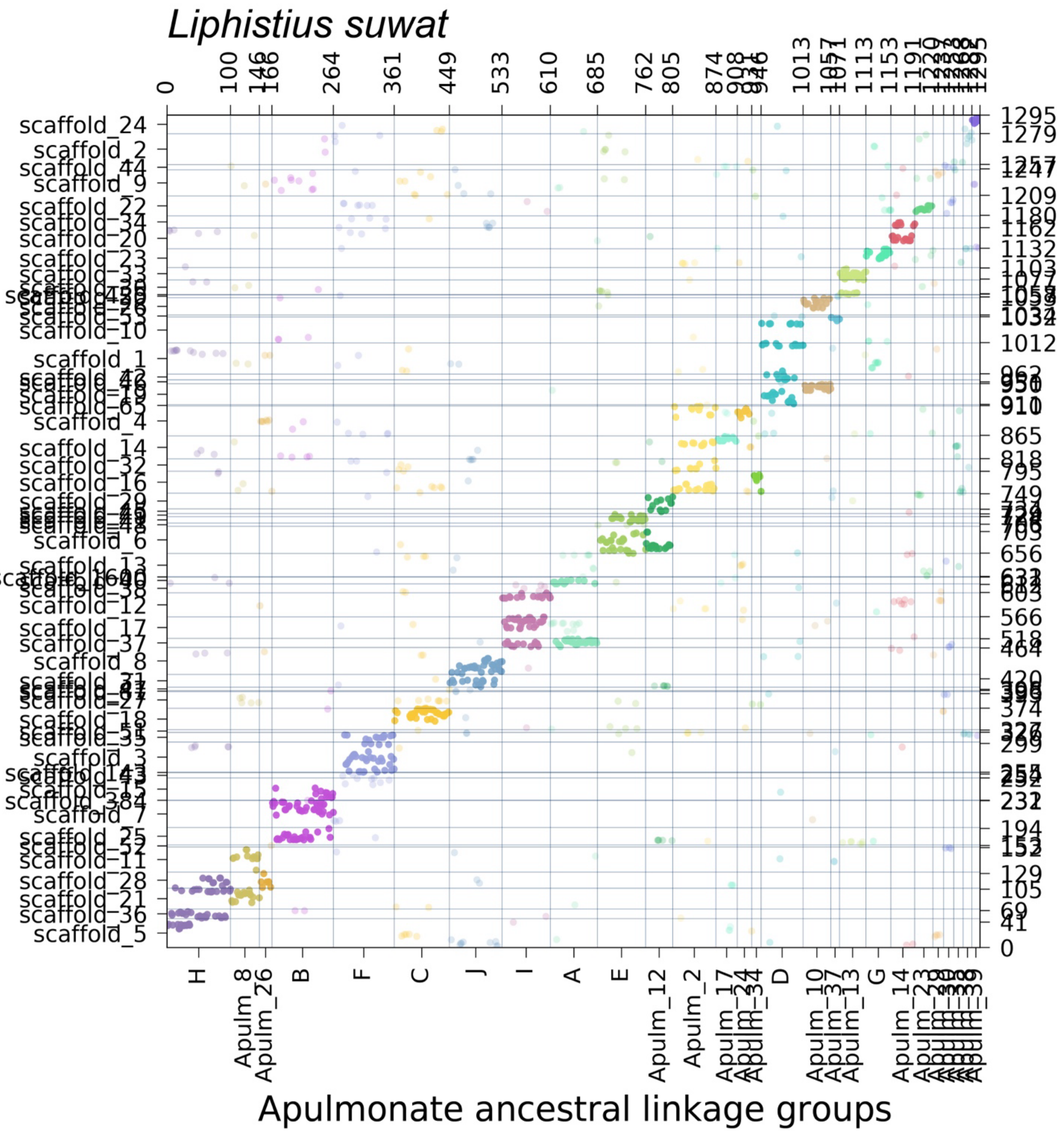

**Supplementary Figure S26.** Oxford dot plot showing the location of apulmonate ancestral linkage group (ALG) homologs in the genome of the vinegaroon *Mastigoproctus giganteus*. Colors correspond to individual ALGs, with solid color indicating significant hits ($p < 0.05$) and transparent color indicating insignificant hits.

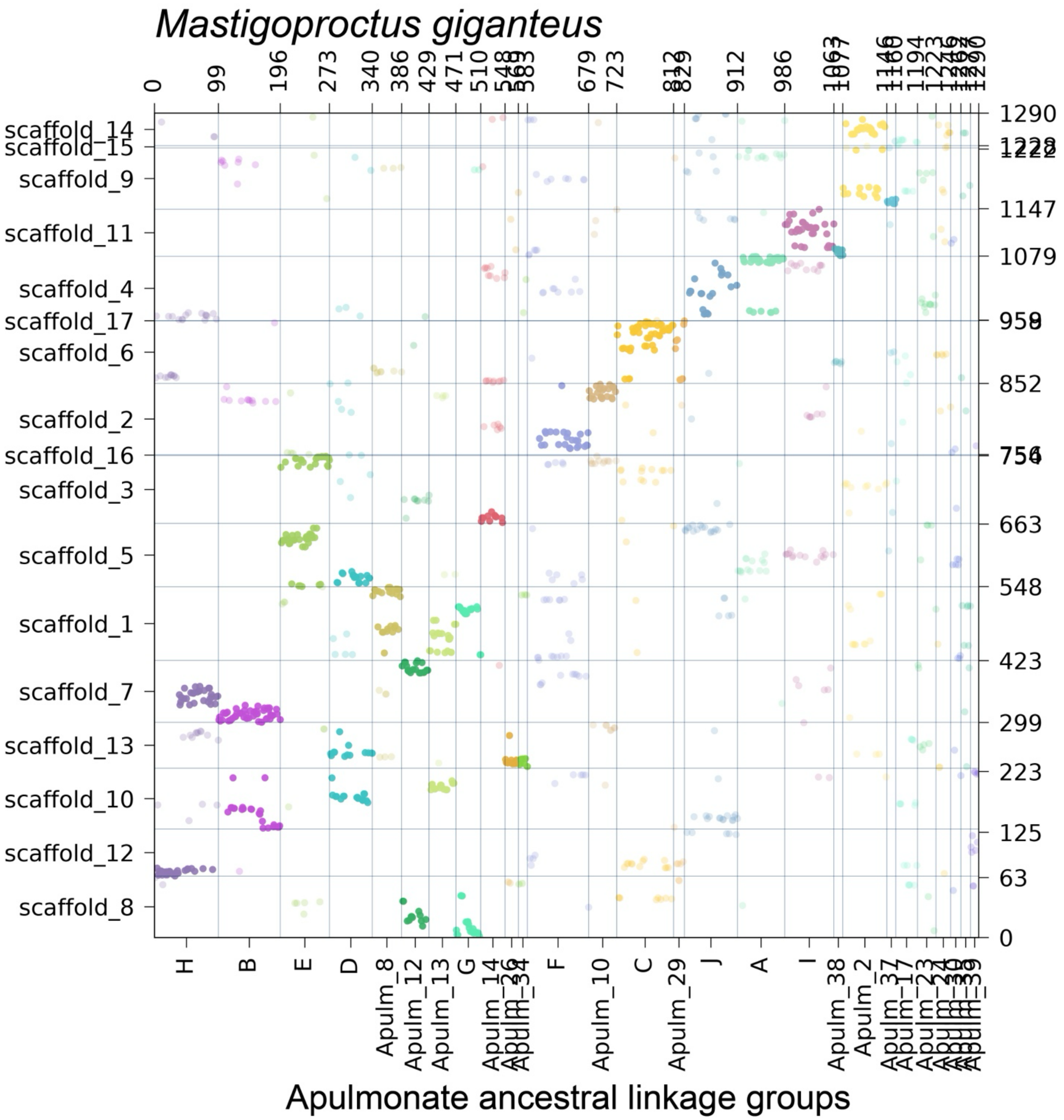

**Supplementary Figure S27.** Oxford dot plot showing the location of apulmonate ancestral linkage group (ALG) homologs in the genome of the harvestman *Odiellus spinosus*. Colors correspond to individual ALGs, with solid color indicating significant hits ($p < 0.05$) and transparent color indicating insignificant hits.

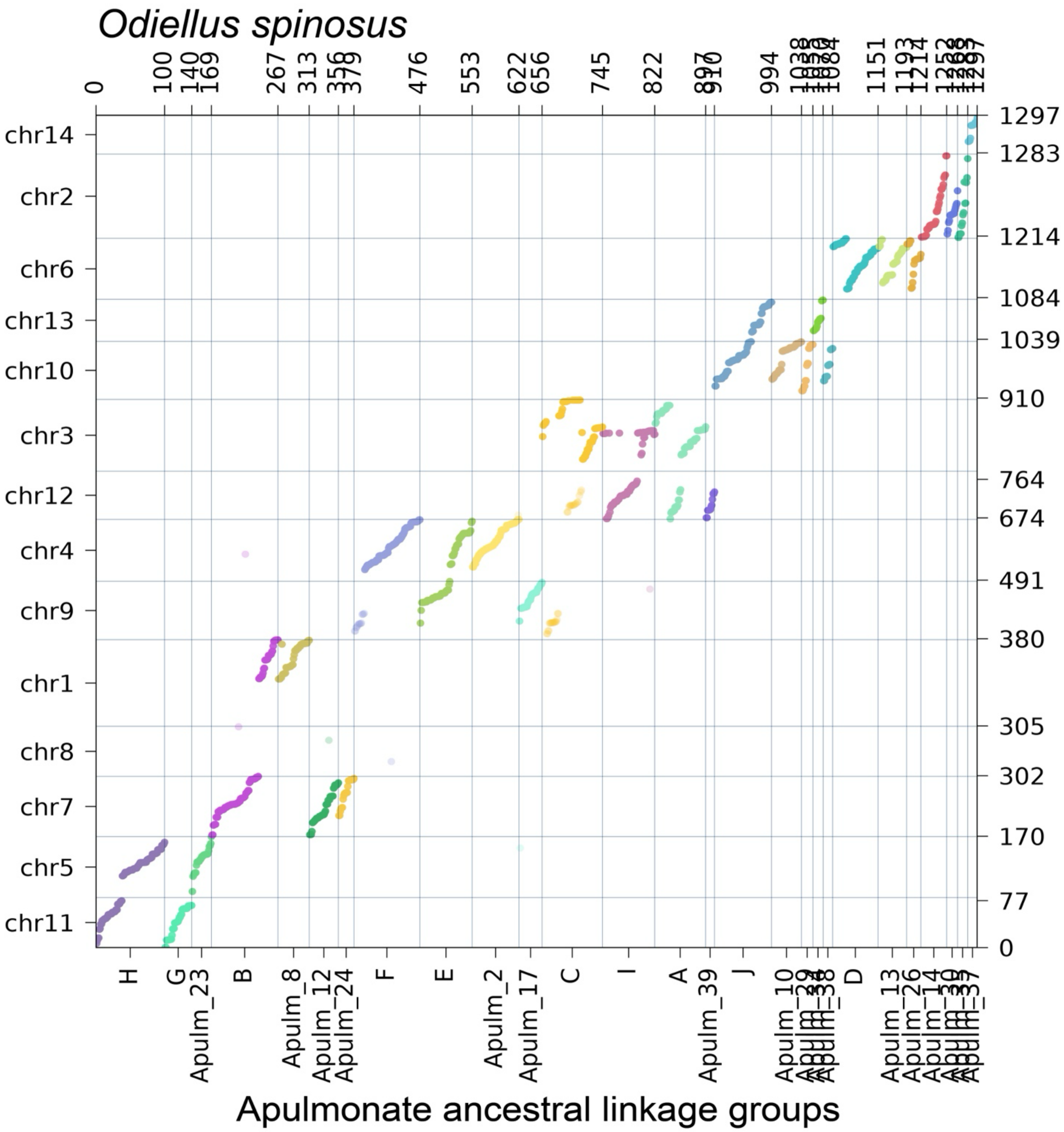

**Supplementary Figure S28.** Oxford dot plot showing the location of apulmonate ancestral linkage group (ALG) homologs in the genome of the acariform mite *Oppia nitens*. Colors correspond to individual ALGs, with solid color indicating significant hits ($p < 0.05$) and transparent color indicating insignificant hits.

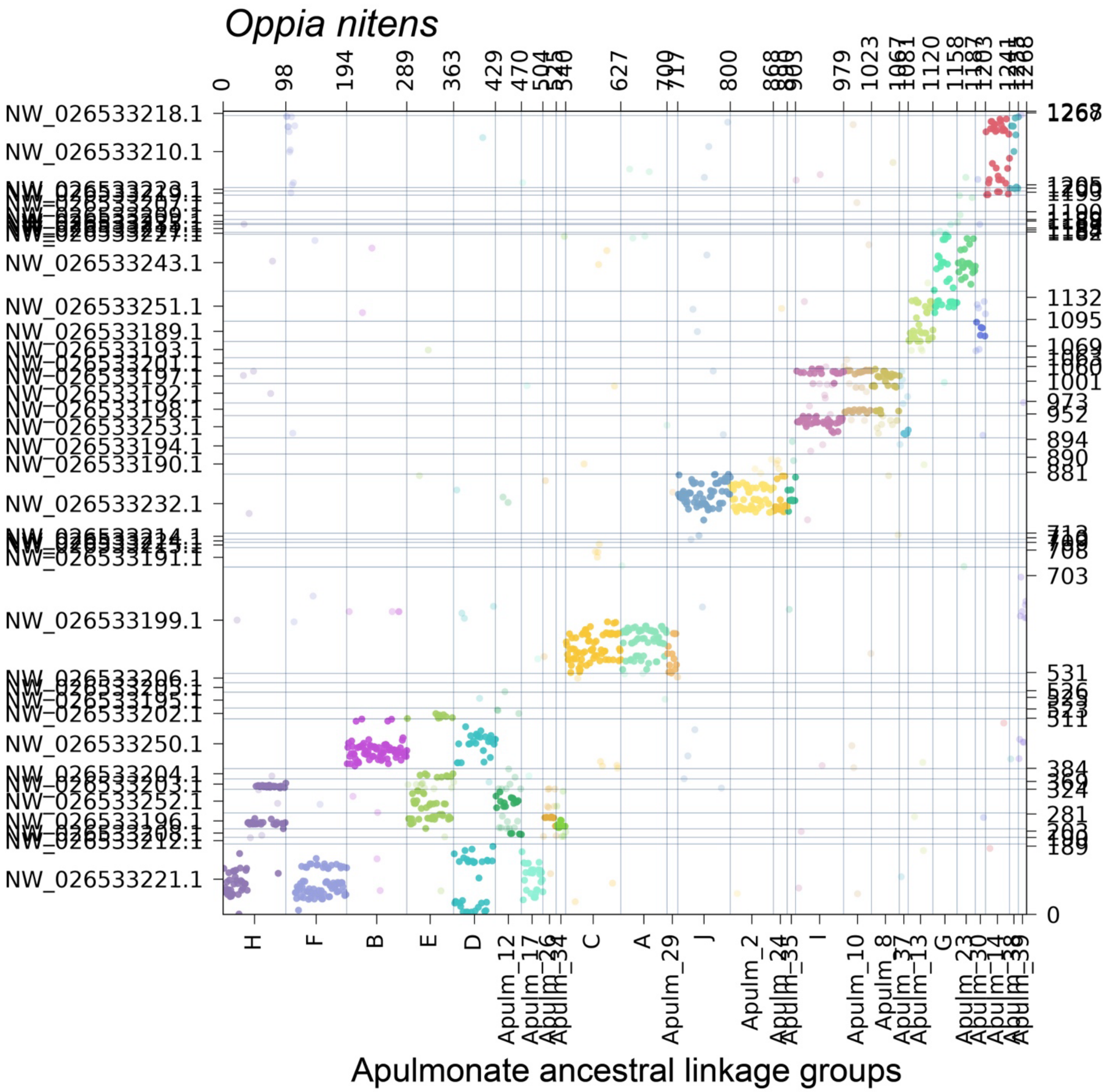

**Supplementary Figure S29.** Oxford dot plot showing the location of apulmonate ancestral linkage group (ALG) homologs in the genome of the sea spider *Pycnogonum litorale*. Colors correspond to individual ALGs, with solid color indicating significant hits ($p < 0.05$) and transparent color indicating insignificant hits.

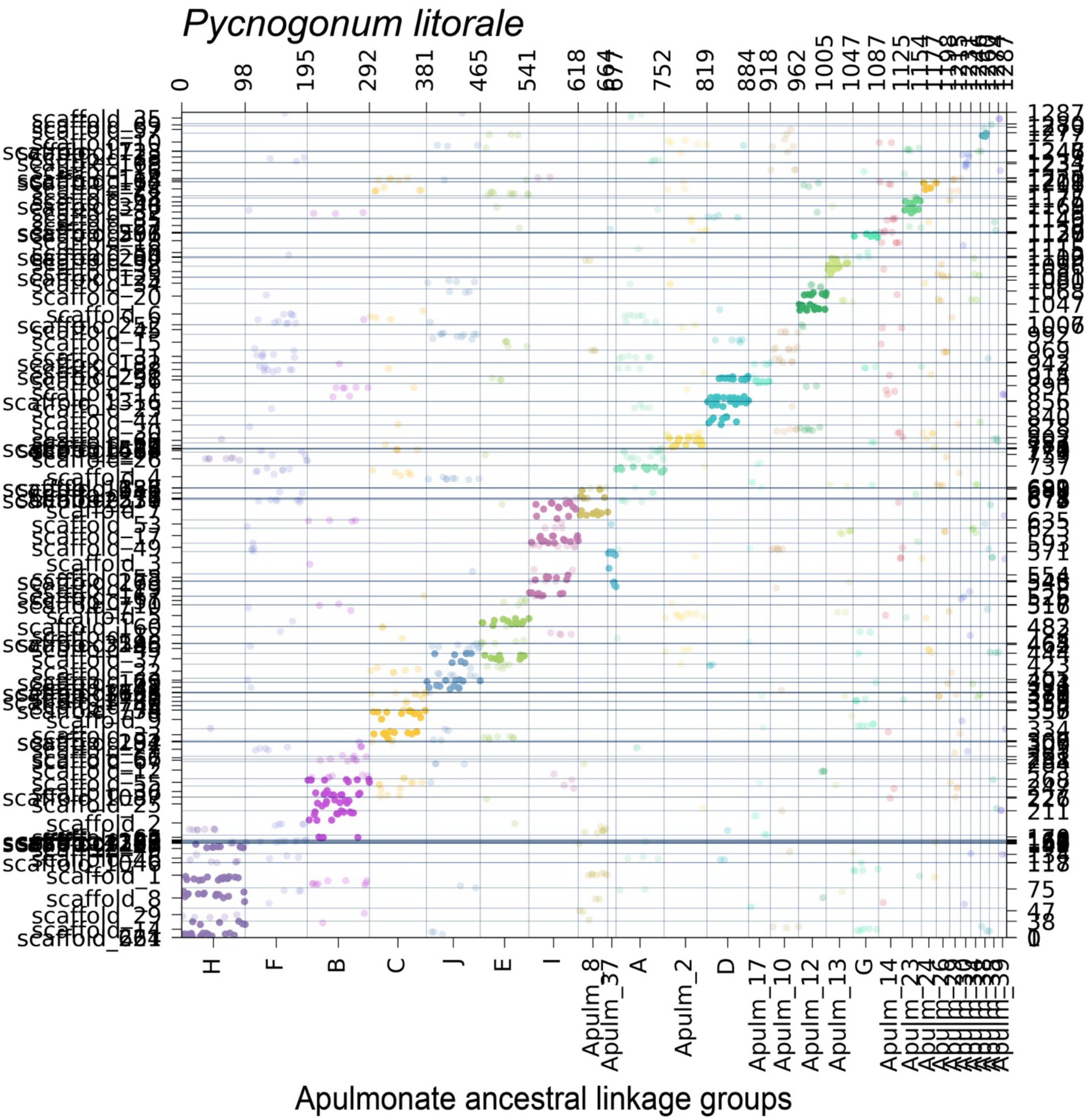

**Supplementary Figure S30.** Oxford dot plot showing the location of apulmonate ancestral linkage group (ALG) homologs in the genome of the parasitiform tick *Rhipicephalus microplus*. Colors correspond to individual ALGs, with solid color indicating significant hits ($p < 0.05$) and transparent color indicating insignificant hits.

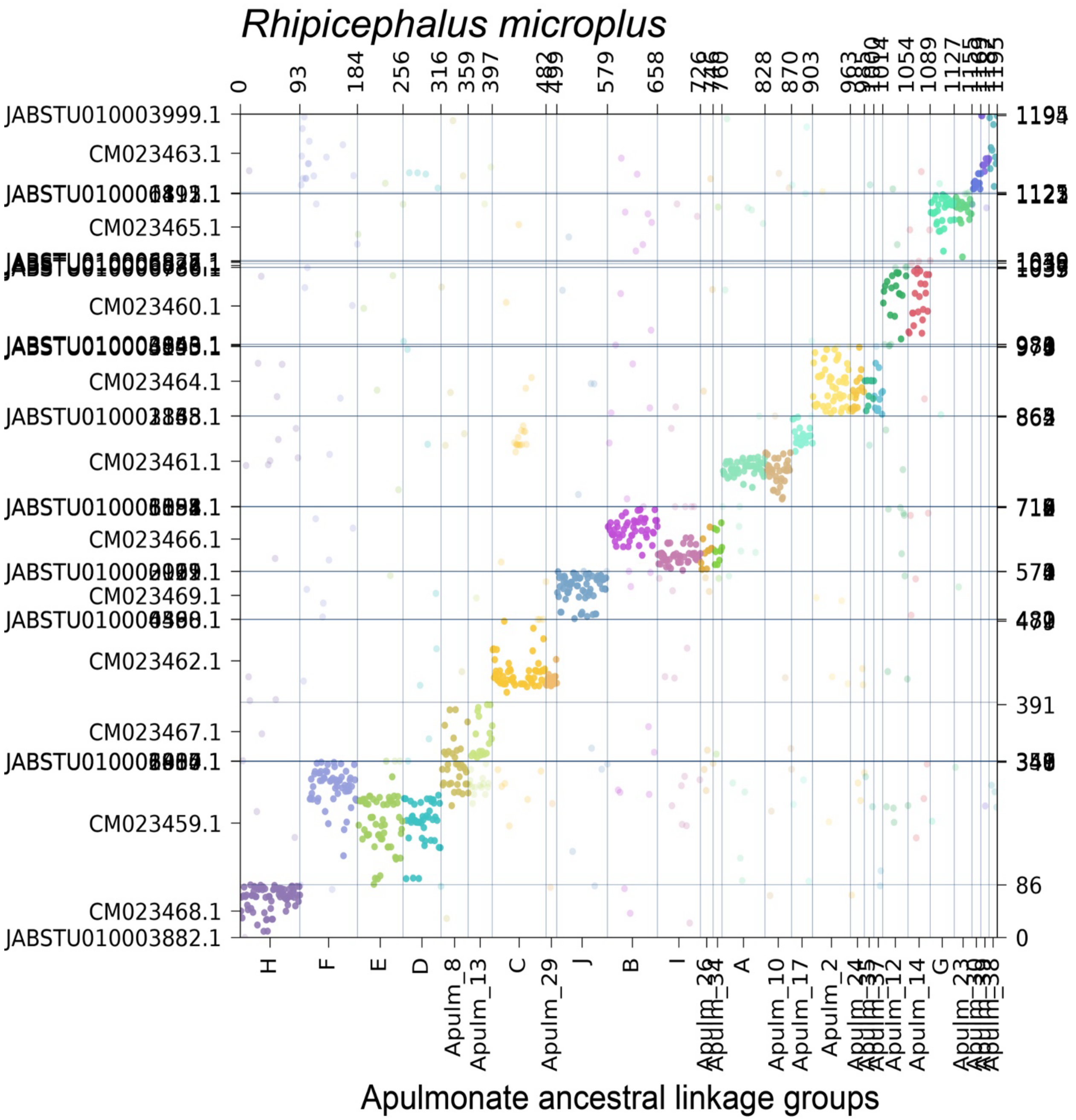

**Supplementary Figure S31.** Oxford dot plot showing the location of apulmonate ancestral linkage group (ALG) homologs in the genome of the acariform mite *Sarcoptes scabiei*. Colors correspond to individual ALGs, with solid color indicating significant hits ($p < 0.05$) and transparent color indicating insignificant hits.

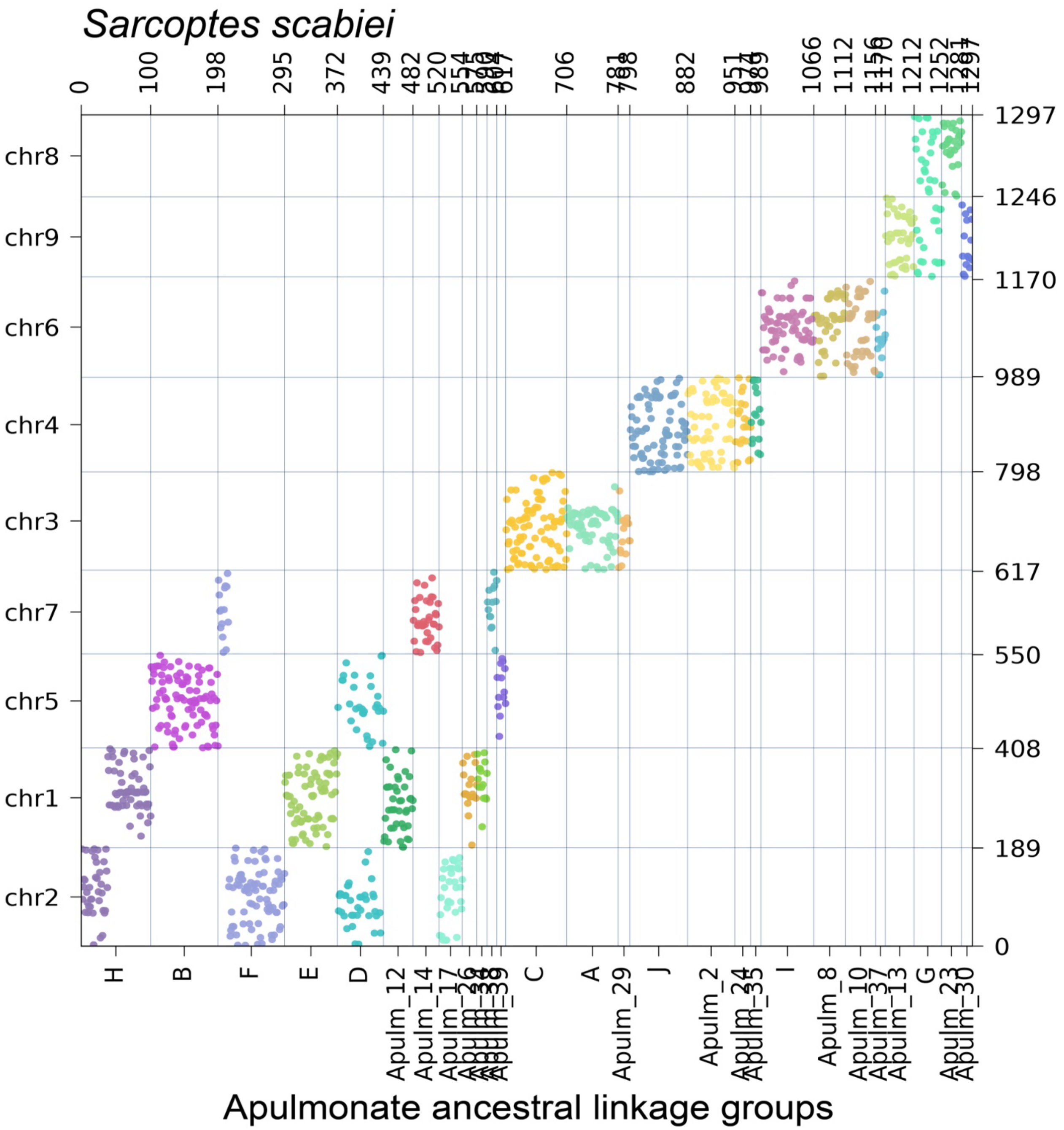

**Supplementary Figure S32.** Oxford dot plot showing the location of apulmonate ancestral linkage group (ALG) homologs in the genome of the centipede *Strigamia acuminata*. Colors correspond to individual ALGs, with solid color indicating significant hits ($p < 0.05$) and transparent color indicating insignificant hits.

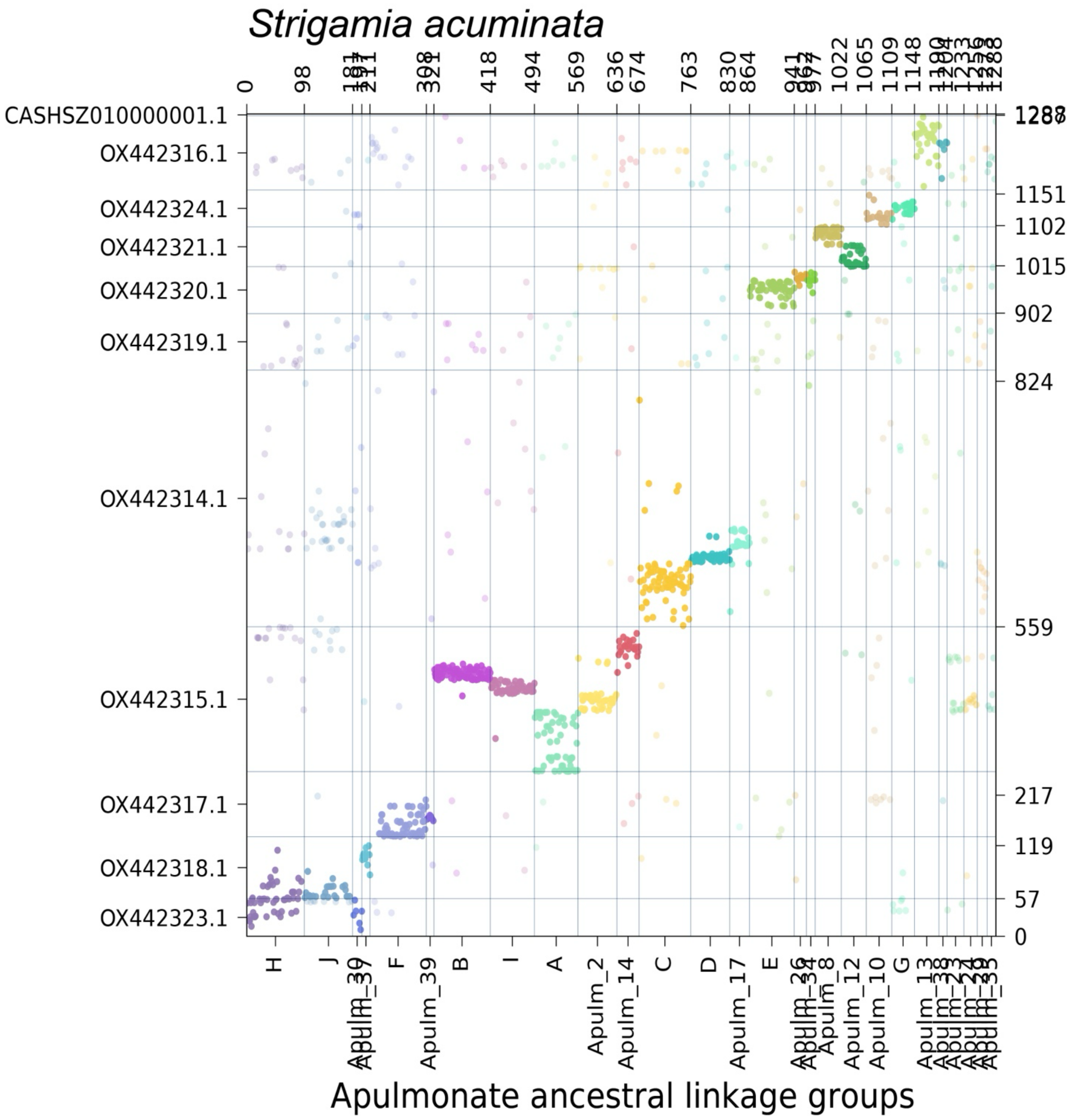

**Supplementary Figure S33.** Oxford dot plot showing the location of apulmonate ancestral linkage group (ALG) homologs in the genome of the horseshoe crab *Tachypleus tridentatus*. Colors correspond to individual ALGs, with solid color indicating significant hits ($p < 0.05$) and transparent color indicating insignificant hits.

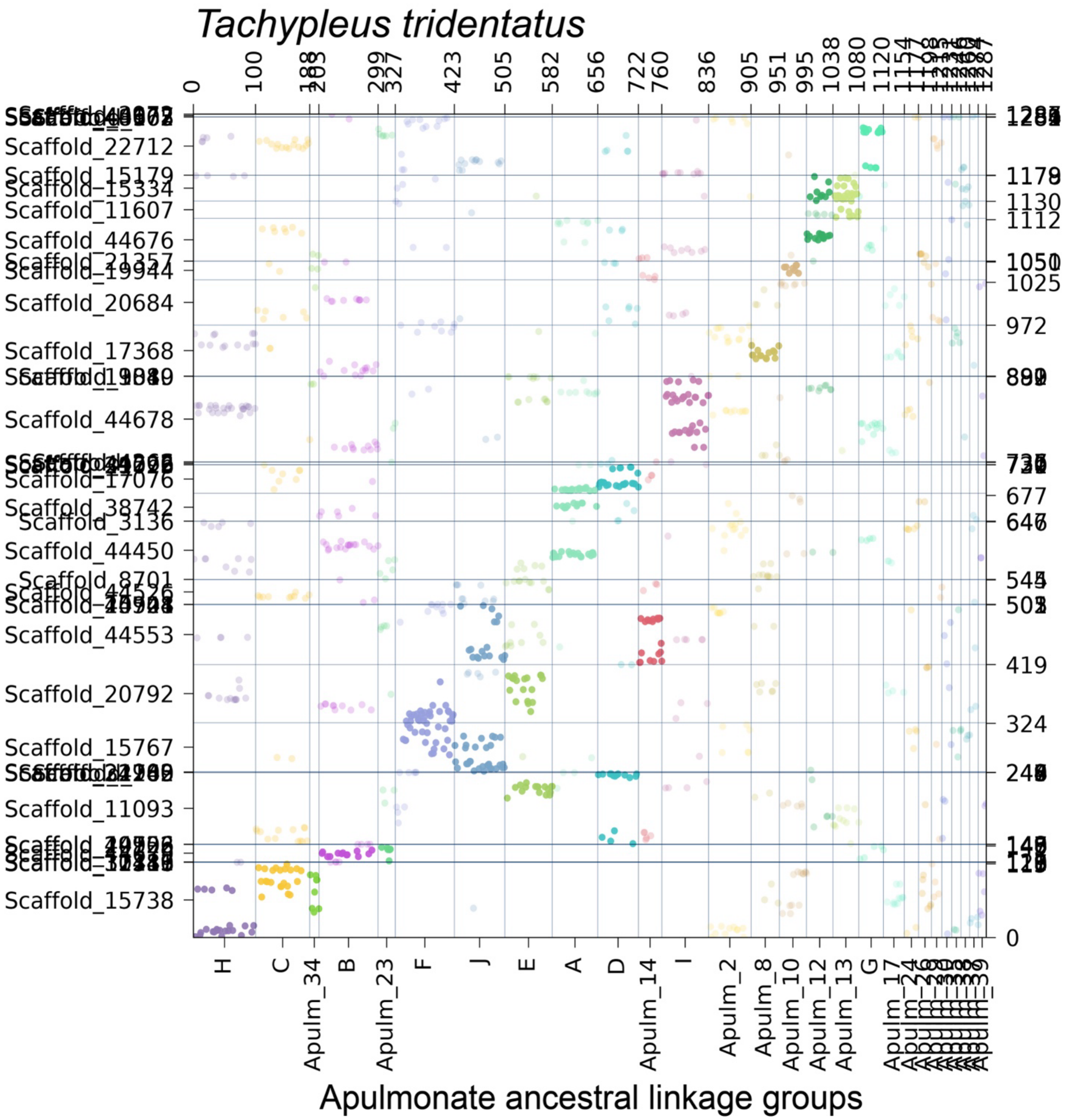

**Supplementary Figure S34.** Oxford dot plot showing the location of apulmonate ancestral linkage group (ALG) homologs in the genome of the parasitiform mite *Varroa destructor*. Colors correspond to individual ALGs, with solid color indicating significant hits ($p < 0.05$) and transparent color indicating insignificant hits.

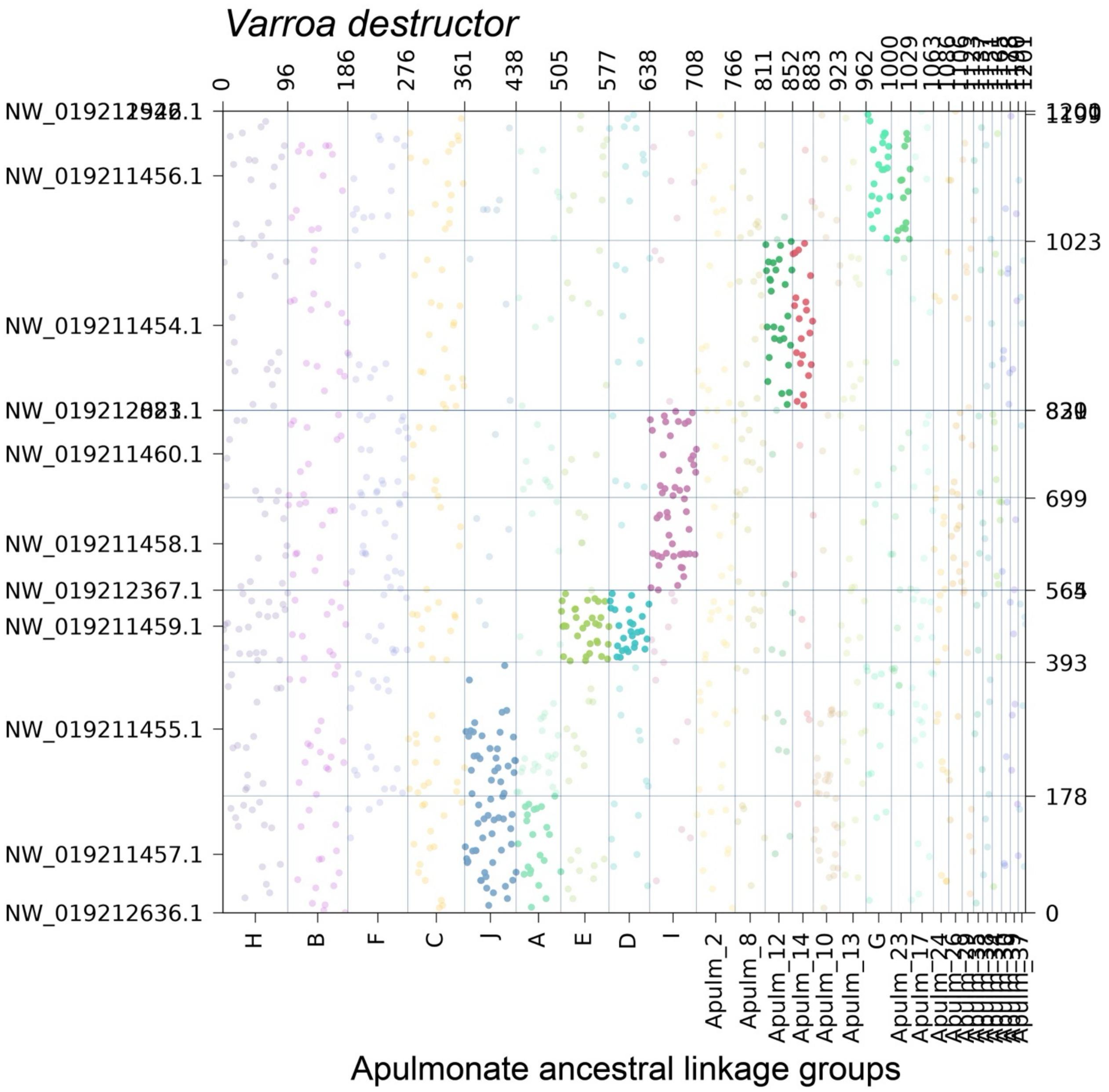

**Supplementary Figure S35.** Oxford dot plots showing the location of apulmonate ancestral linkage group (ALG) homologs in the genomes of two solifuges for the apulmonate (top row) and the BCnS (bottom row) linkage groups. Left column: *Gluvia dorsalis.* Right column: *Paragaleodes pallidus*. Colors correspond to individual ALGs, with solid color indicating significant hits ($p < 0.05$) and transparent color indicating insignificant hits.

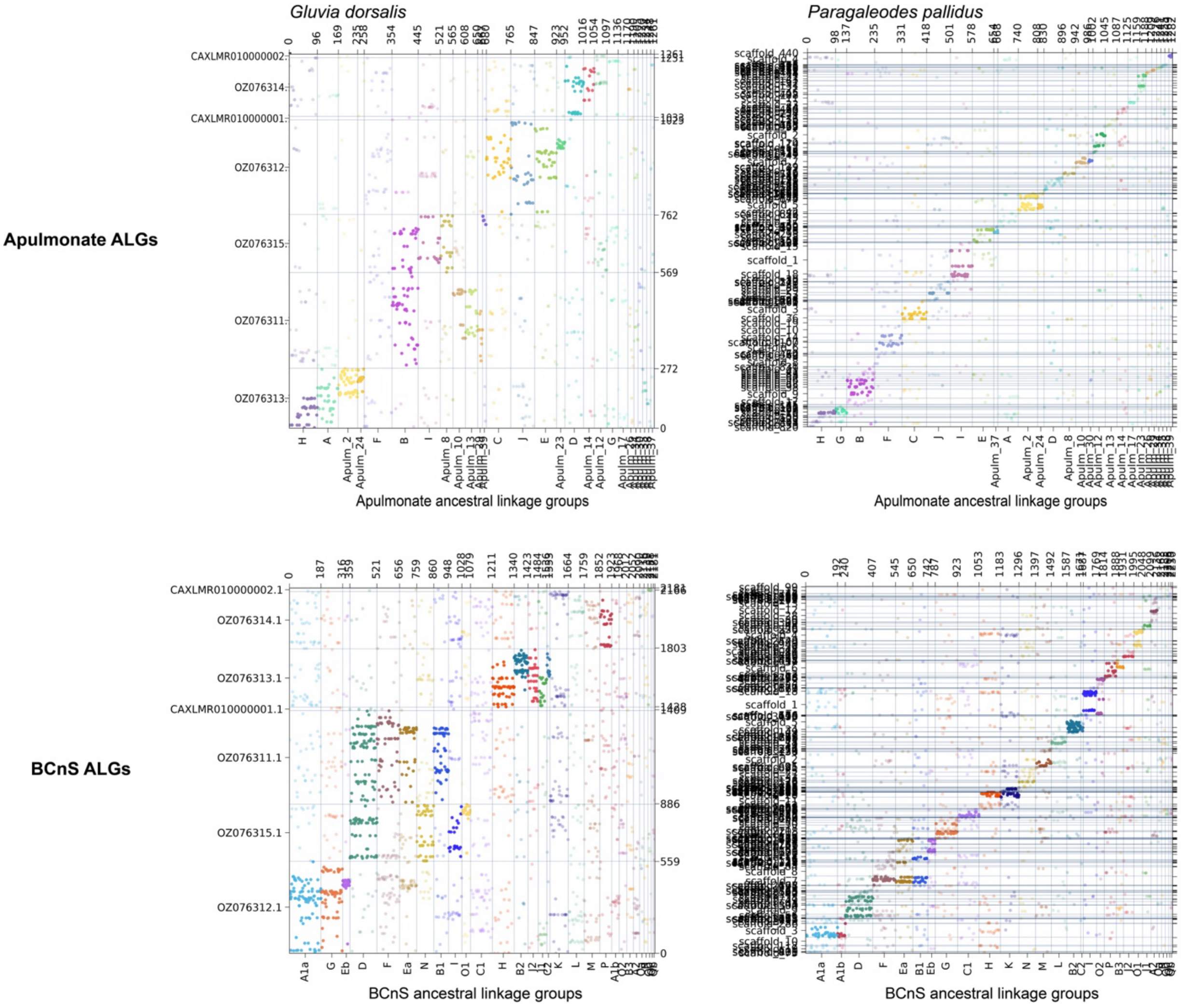

**Supplementary Figure S36.** Insignificant cases of fusion-with-mixing in Acariformes + Ricinulei. **a**, Oxford dot plot of the hooded tick spider *Cryptocellus* cf. *goodnighti* showing fusion-with-mixing events that are putatively shared with Acariformes (blue outlines). Dotted lines indicate mixing events with fewer than 40 genes included and fewer than five minimum swap events. **b**, Oxford dot plot of the acariform mite *Sarcoptes scabiei* showing the same three events as in **a**. **c**, Summary of patterns shown in panels **a** and **b**. Question marks correspond to insignificant mixing. **d**, Ribbon plots showing insignificant mixing for two cases (I ⊗ 8 and 14 ⊗ 38).

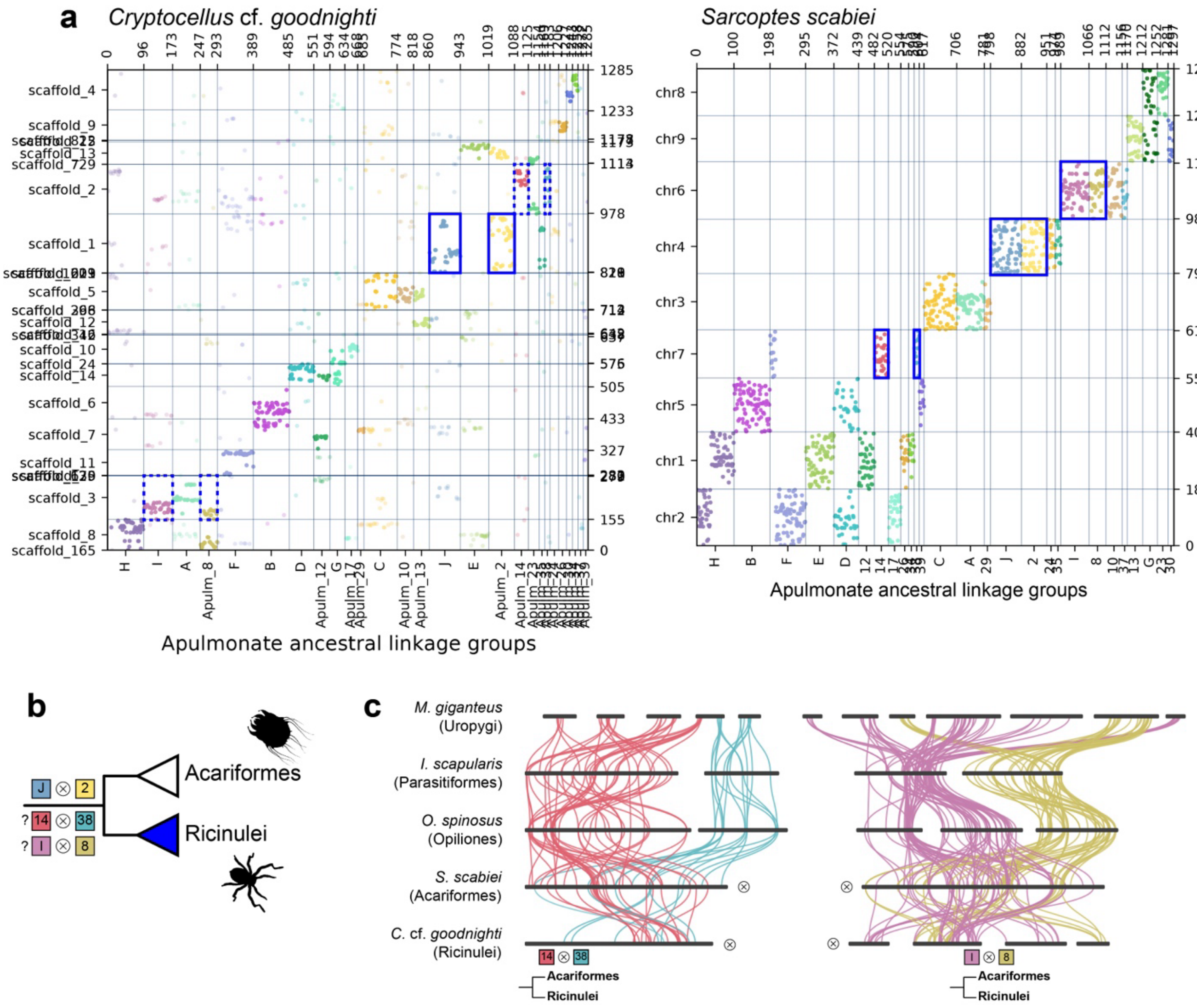

**Supplementary Figure S37.** Oxford dot plot showing the location of BCnS ancestral linkage group (ALG) homologs in the genome of the mygalomorph spider *Aphonopelma hentzi*. Colors correspond to individual ALGs, with solid color indicating significant hits ($p < 0.05$) and transparent color indicating insignificant hits.

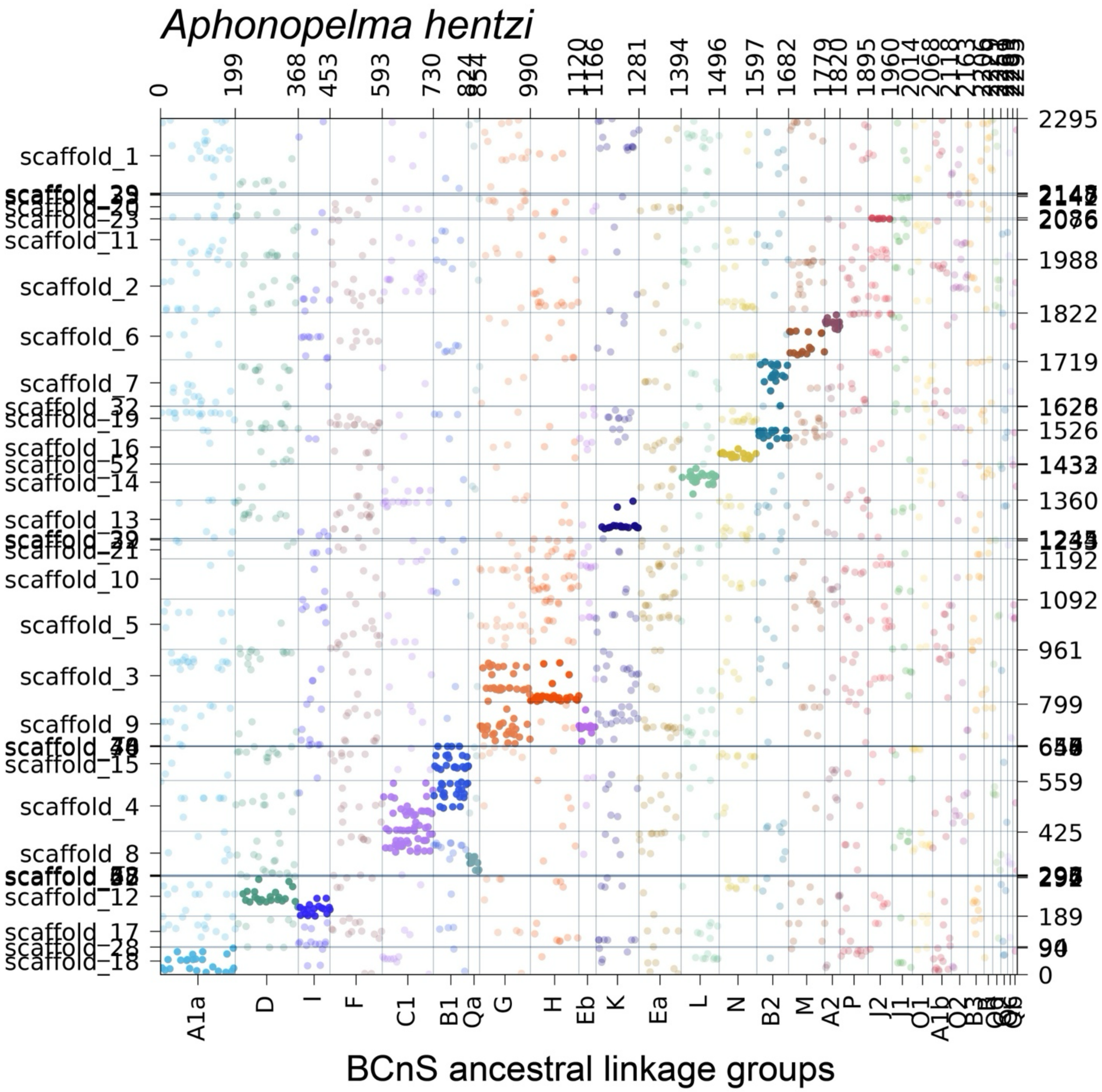

**Supplementary Figure S38.** Oxford dot plot showing the location of BCnS ancestral linkage group (ALG) homologs in the genome of the araneomorph spider *Argiope bruennichi*. Colors correspond to individual ALGs, with solid color indicating significant hits ($p < 0.05$) and transparent color indicating insignificant hits.

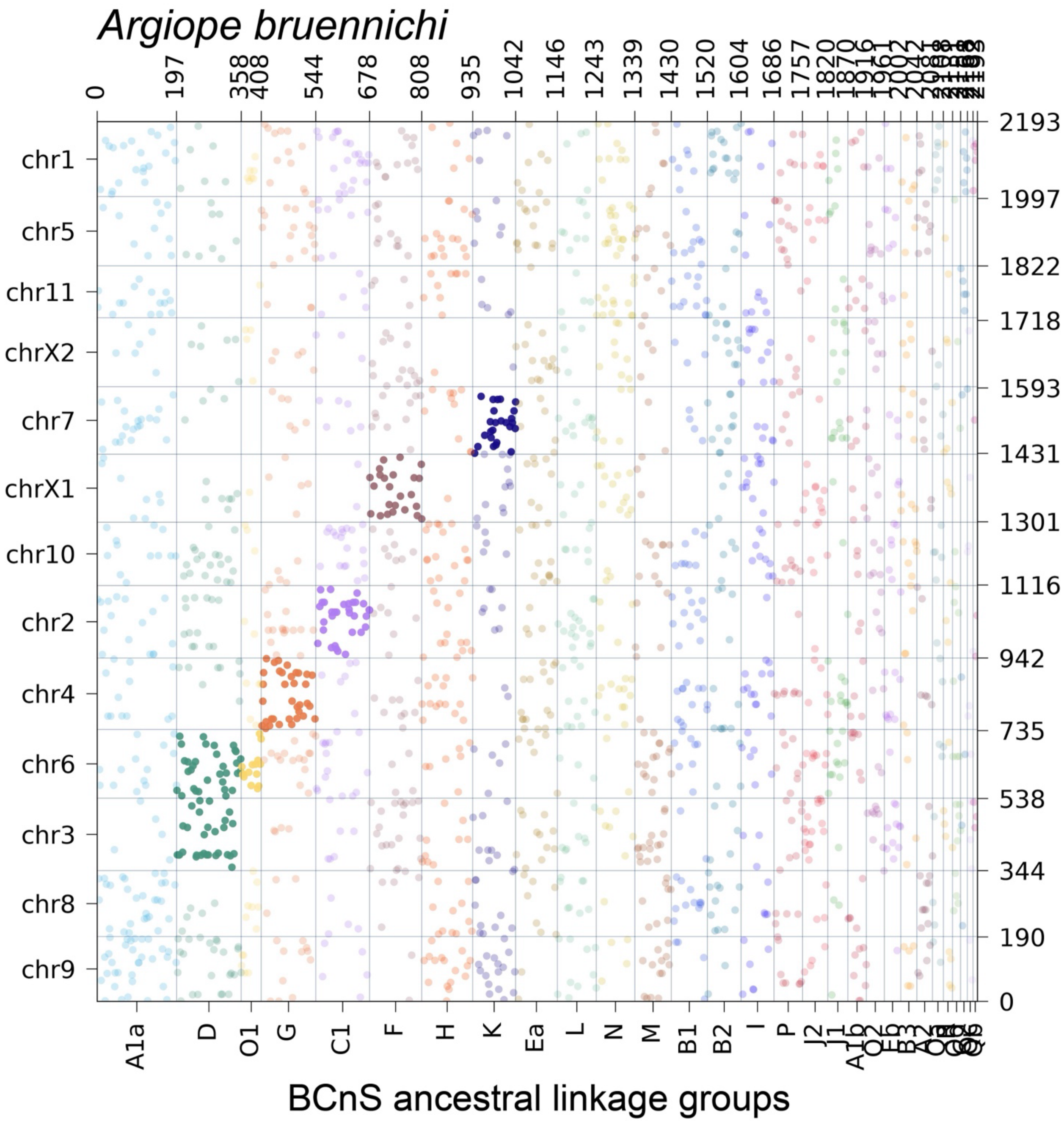

**Supplementary Figure S39.** Oxford dot plot showing the location of BCnS ancestral linkage group (ALG) homologs in the genome of the acariform mite *Blomia tropicalis*. Colors correspond to individual ALGs, with solid color indicating significant hits ($p < 0.05$) and transparent color indicating insignificant hits.

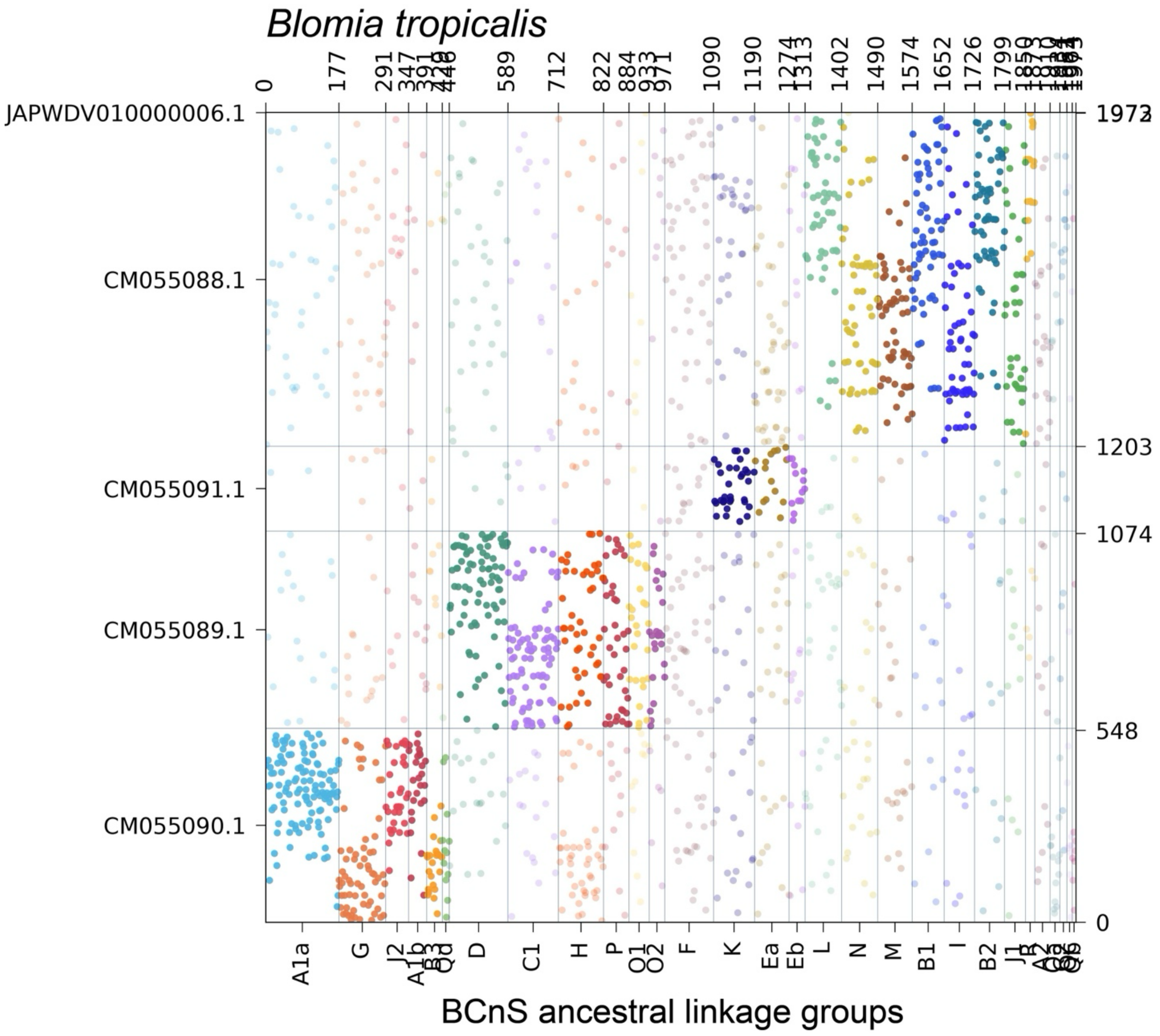

**Supplementary Figure S40.** Oxford dot plot showing the location of BCnS ancestral linkage group (ALG) homologs in the genome of the horseshoe crab *Carcinoscorpius rotundicauda*. Colors correspond to individual ALGs, with solid color indicating significant hits ($p < 0.05$) and transparent color indicating insignificant hits.

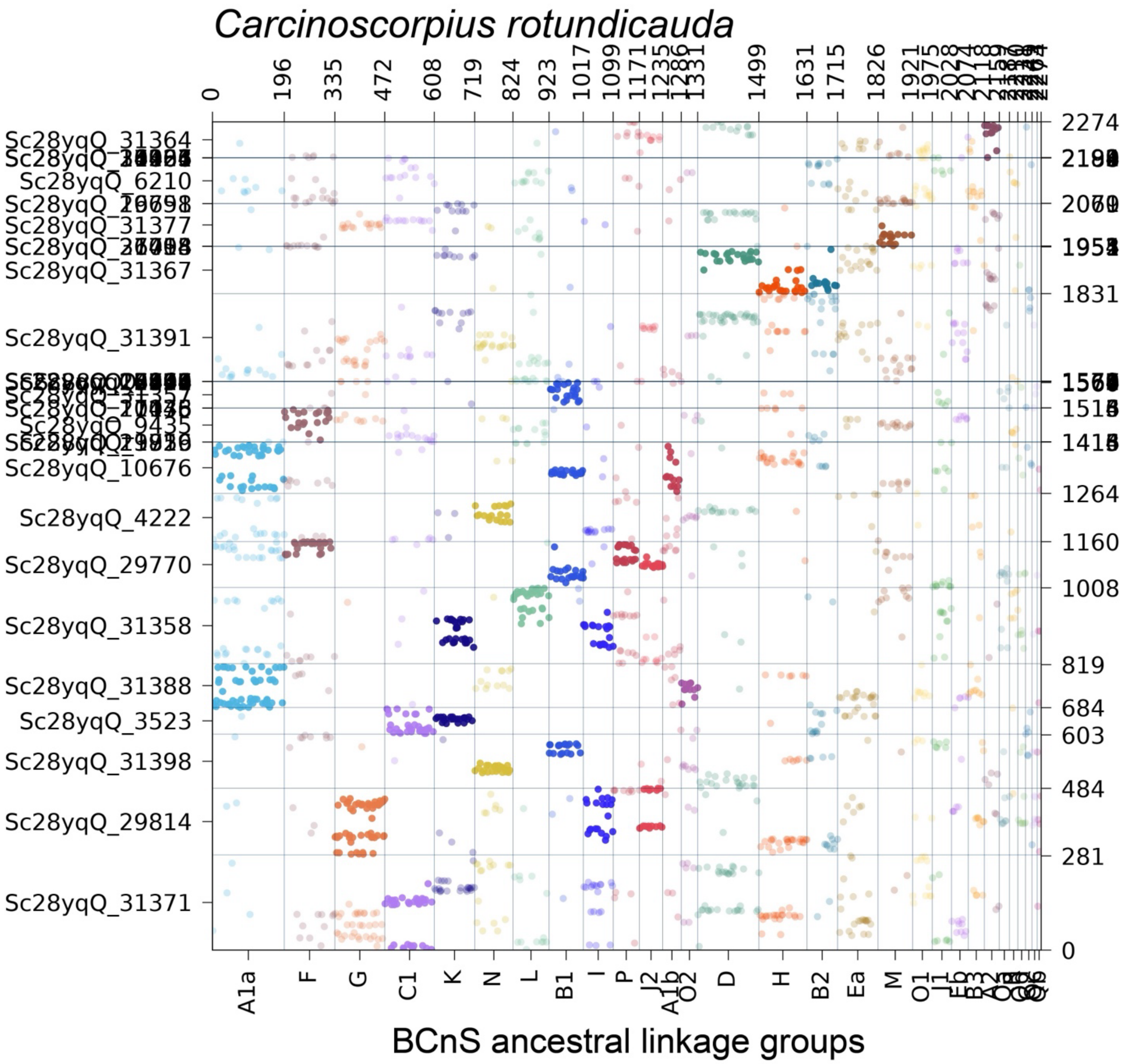

**Supplementary Figure S41.** Oxford dot plot showing the location of BCnS ancestral linkage group (ALG) homologs in the genome of the scorpion *Centruroides sculpturatus*. Colors correspond to individual ALGs, with solid color indicating significant hits ($p < 0.05$) and transparent color indicating insignificant hits.

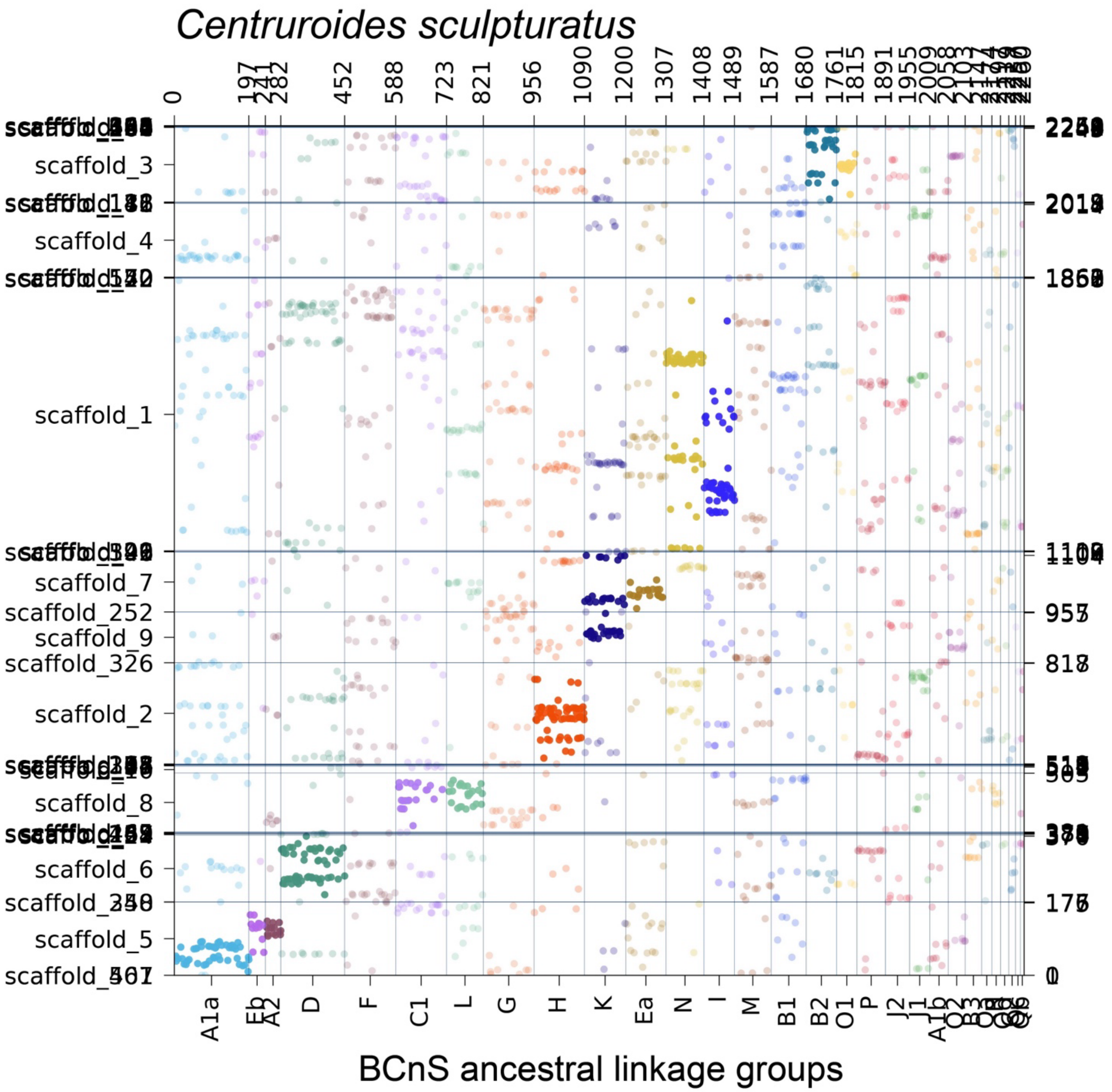

**Supplementary Figure S42.** Oxford dot plot showing the location of BCnS ancestral linkage group (ALG) homologs in the genome of the pseudoscorpion *Cordylochernes scorpioides*. Colors correspond to individual ALGs, with solid color indicating significant hits ($p < 0.05$) and transparent color indicating insignificant hits.

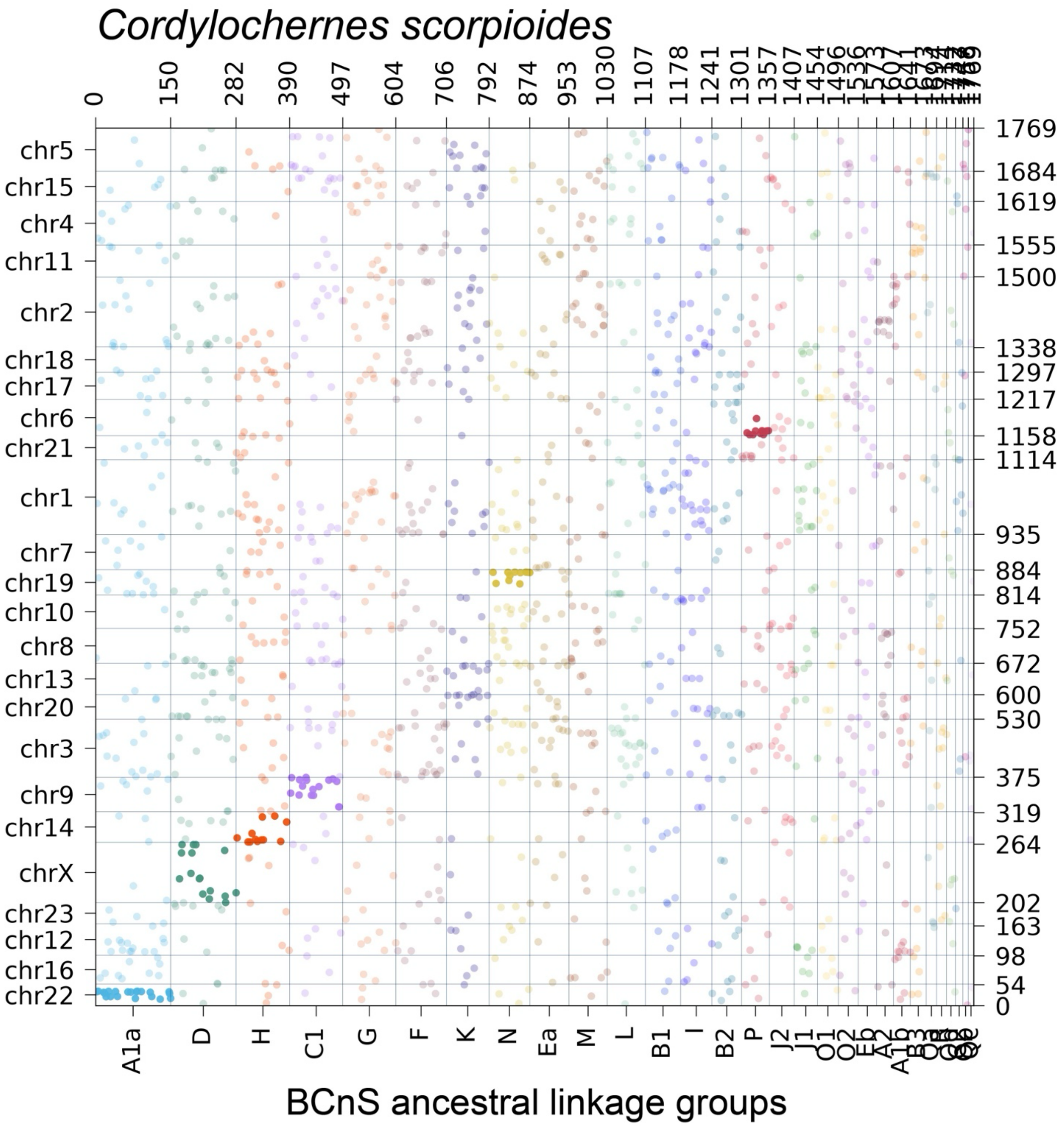

**Supplementary Figure S43.** Oxford dot plot showing the location of BCnS ancestral linkage group (ALG) homologs in the genome of the ricinuleid *Cryptocellus* cf. *goodnighti*. Colors correspond to individual ALGs, with solid color indicating significant hits ($p < 0.05$) and transparent color indicating insignificant hits.

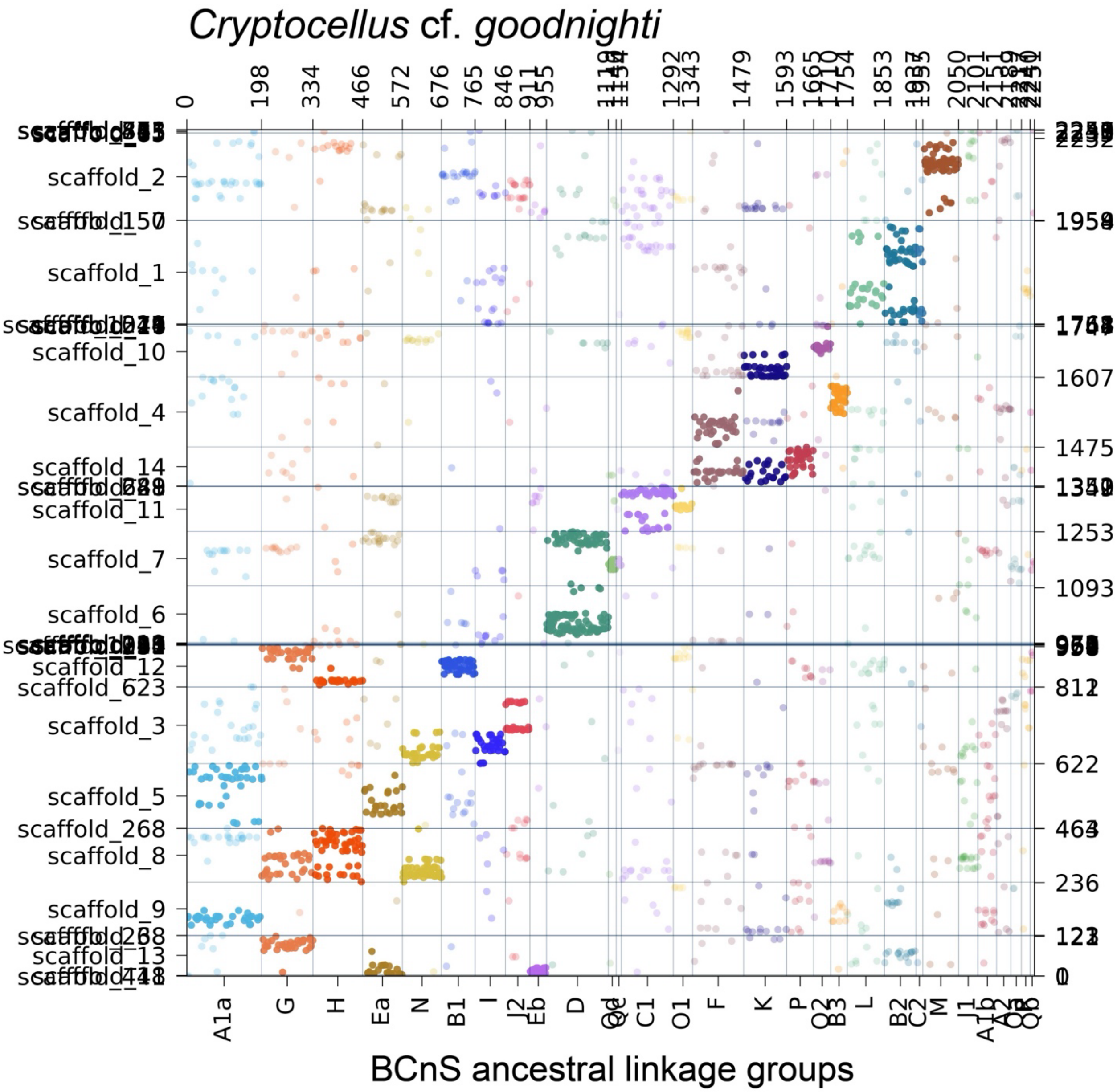

**Supplementary Figure S44.** Oxford dot plot showing the location of BCnS ancestral linkage group (ALG) homologs in the genome of the parasitiform tick *Dermacentor silvarum*. Colors correspond to individual ALGs, with solid color indicating significant hits ($p < 0.05$) and transparent color indicating insignificant hits.

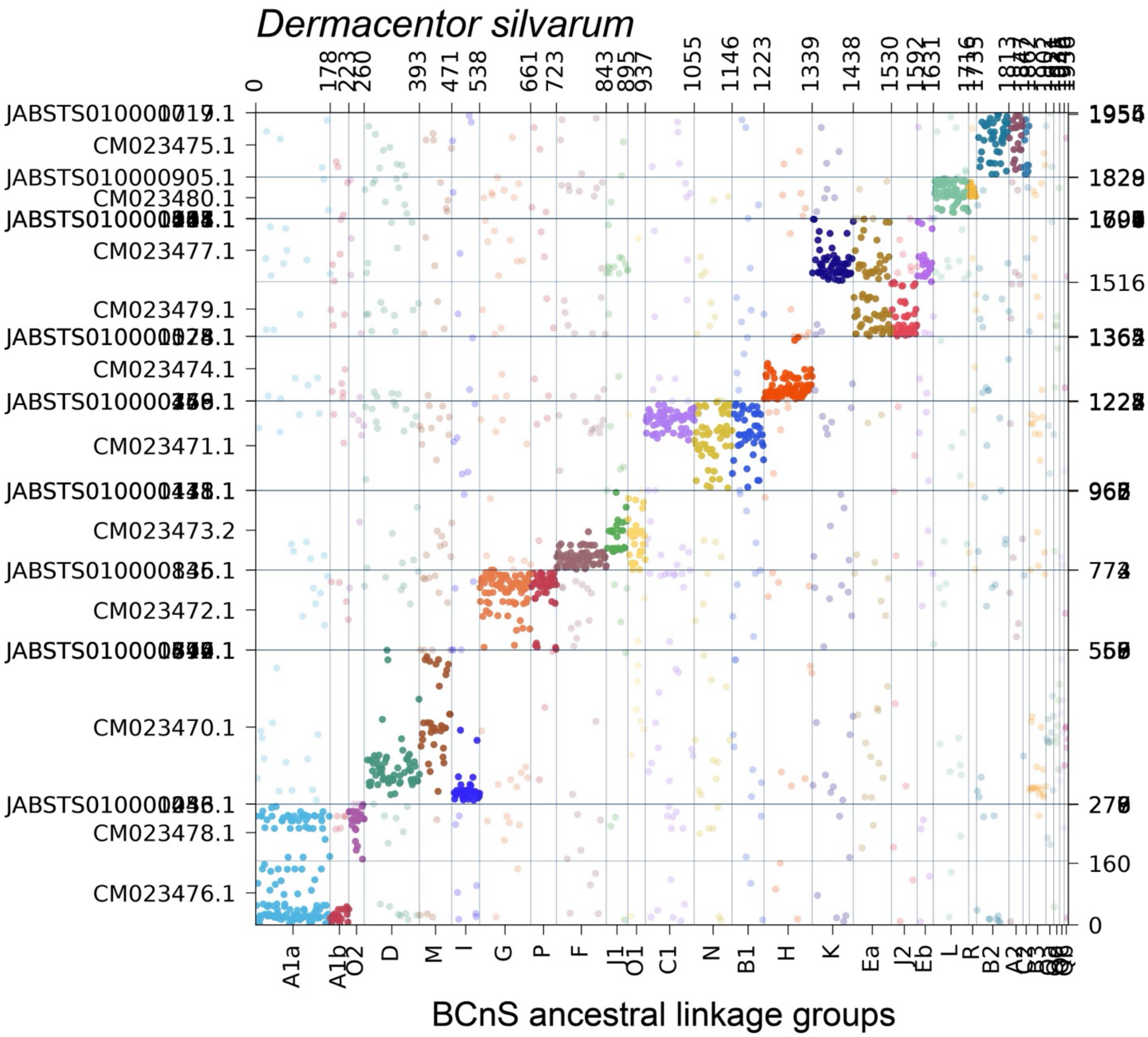

**Supplementary Figure S45.** Oxford dot plot showing the location of BCnS ancestral linkage group (ALG) homologs in the genome of the solifuge *Gluvia dorsalis*. Colors correspond to individual ALGs, with solid color indicating significant hits ($p < 0.05$) and transparent color indicating insignificant hits.

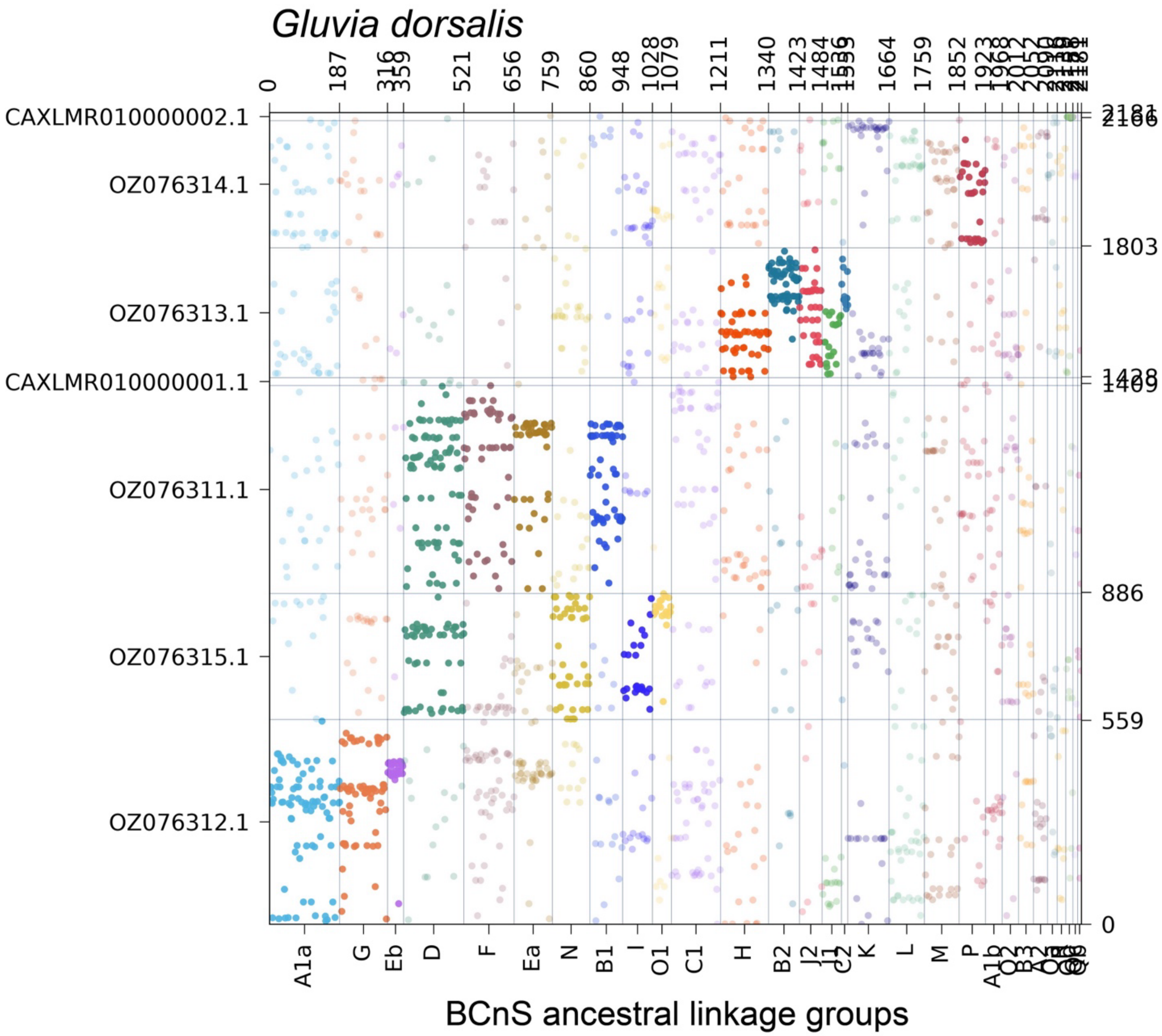

**Supplementary Figure S46.** Oxford dot plot showing the location of BCnS ancestral linkage group (ALG) homologs in the genome of the scorpion *Hadrurus arizonensis*. Colors correspond to individual ALGs, with solid color indicating significant hits ($p < 0.05$) and transparent color indicating insignificant hits.

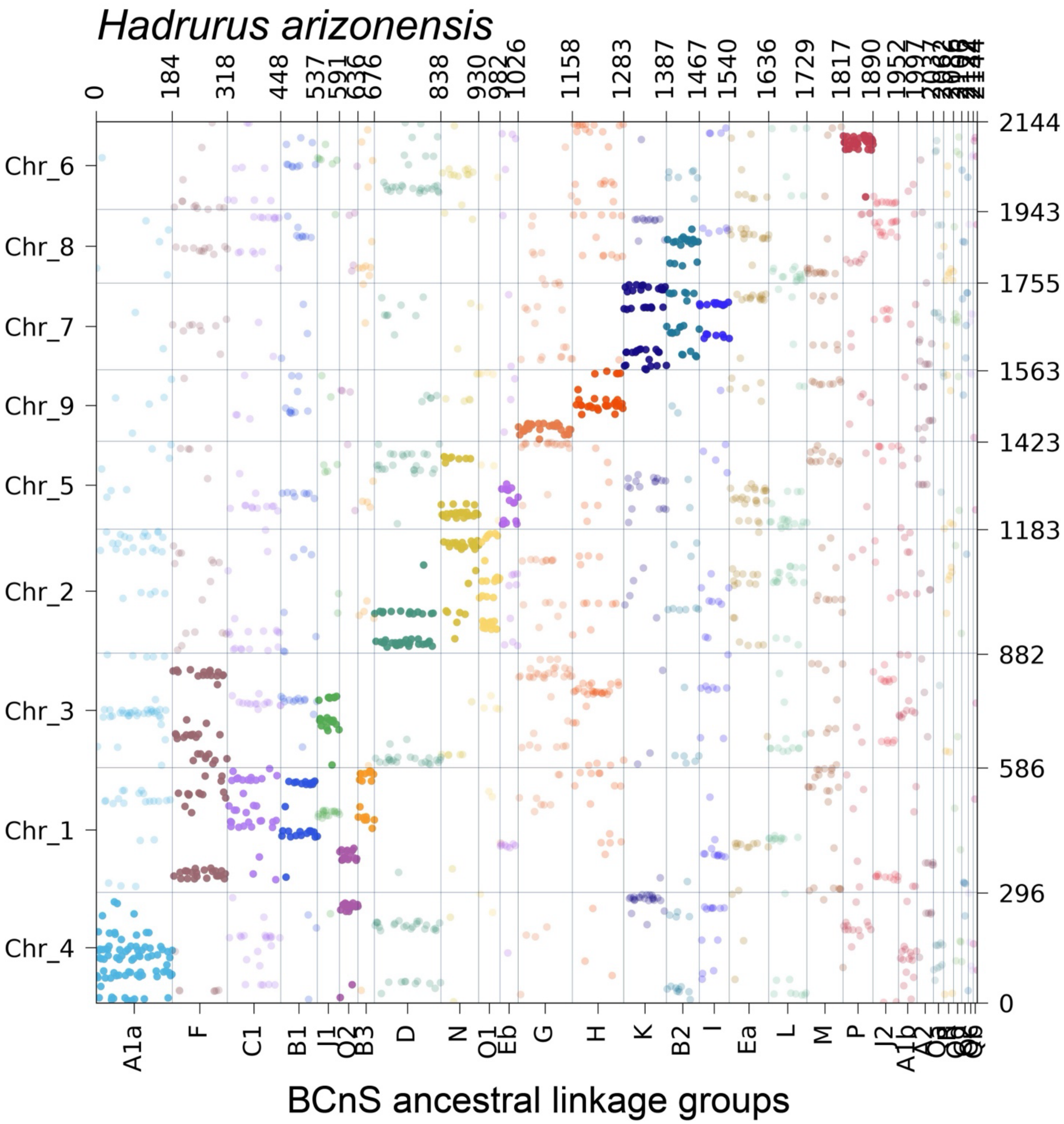

**Supplementary Figure S47.** Oxford dot plot showing the location of BCnS ancestral linkage group (ALG) homologs in the genome of the parasitiform tick *Haemaphysalis longicornis*. Colors correspond to individual ALGs, with solid color indicating significant hits ($p < 0.05$) and transparent color indicating insignificant hits.

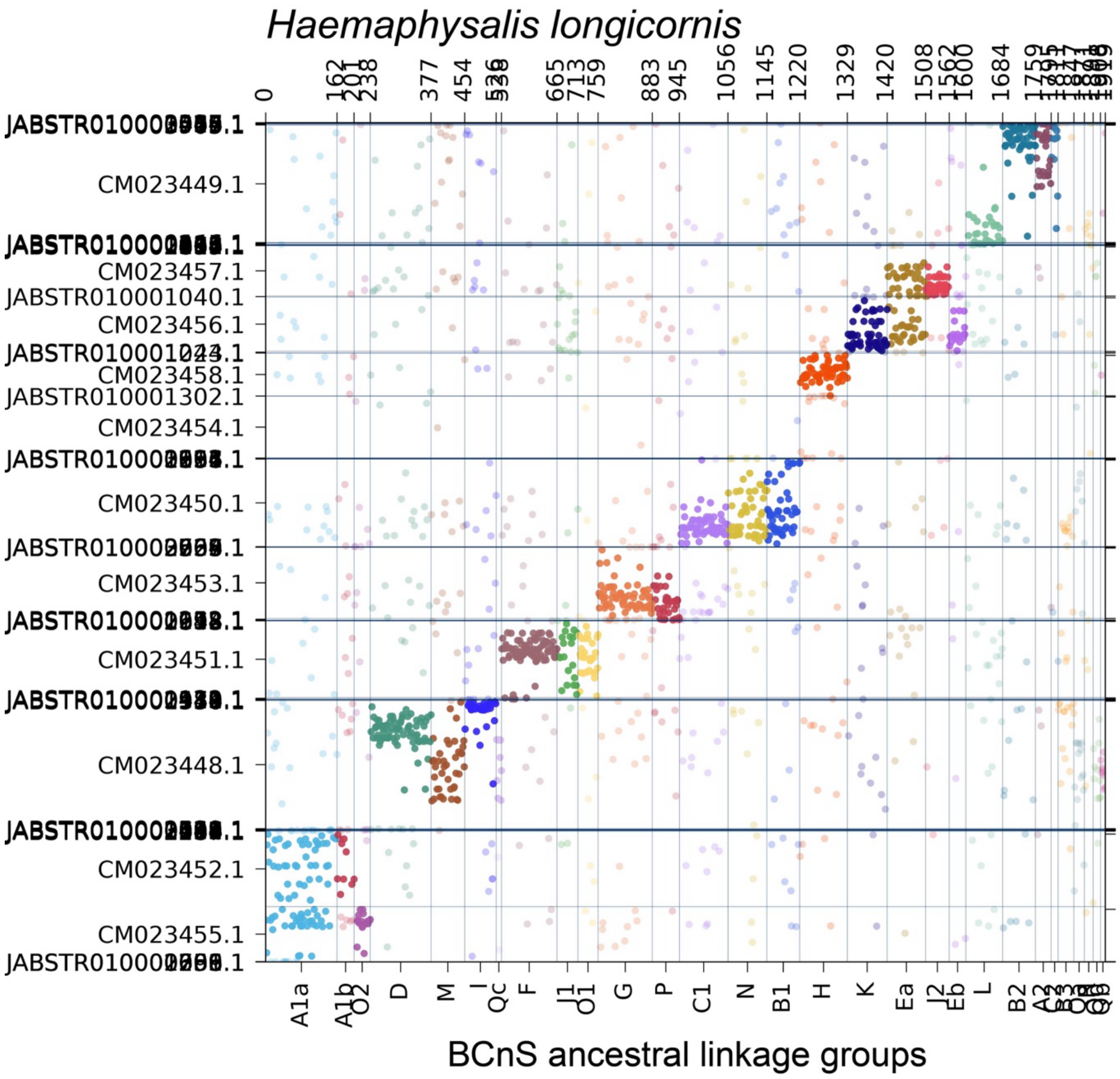

**Supplementary Figure S48.** Oxford dot plot showing the location of BCnS ancestral linkage group (ALG) homologs in the genome of the acariform mite *Halotydeus destructor*. Colors correspond to individual ALGs, with solid color indicating significant hits ($p < 0.05$) and transparent color indicating insignificant hits.

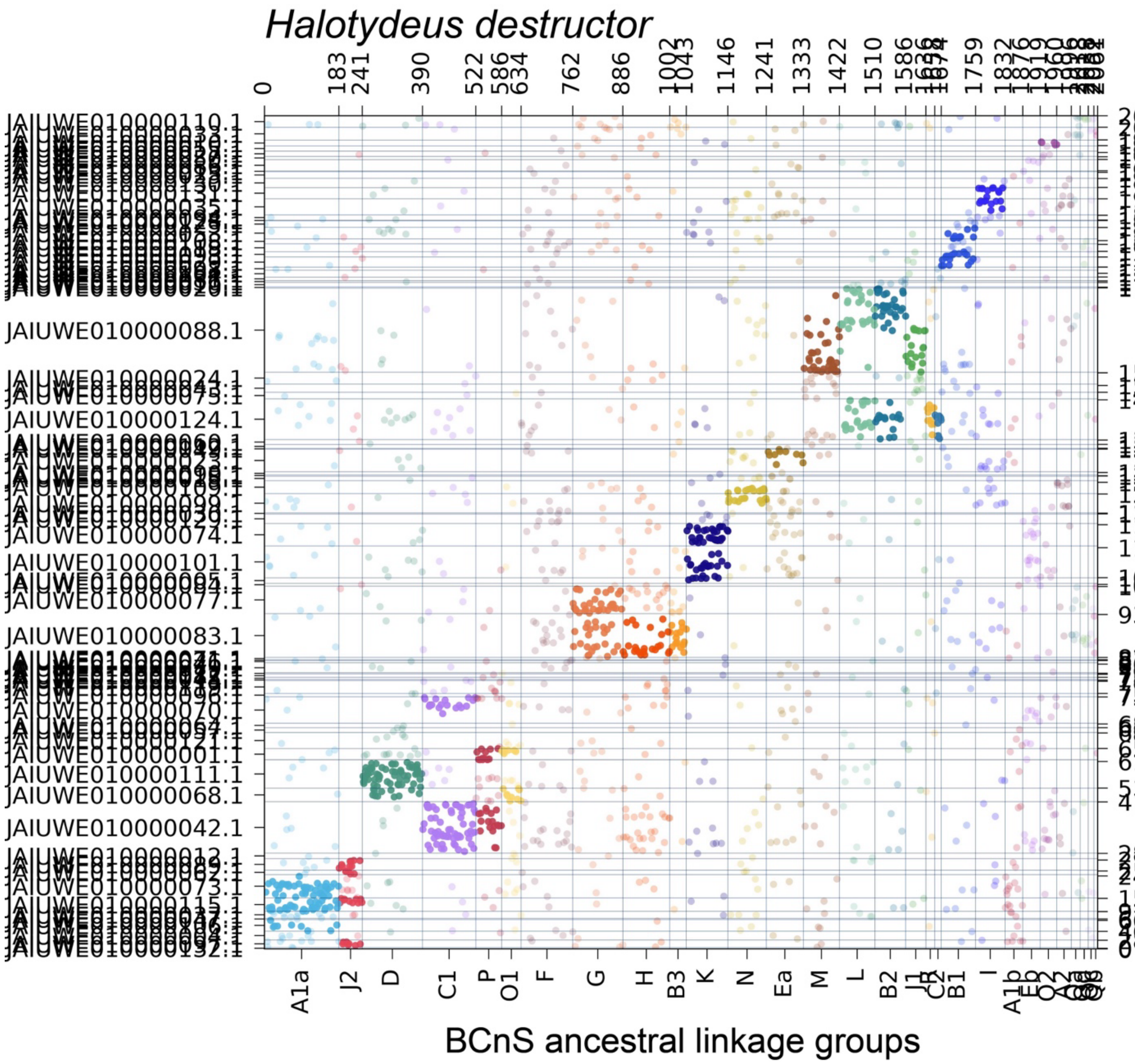

**Supplementary Figure S49.** Oxford dot plot showing the location of BCnS ancestral linkage group (ALG) homologs in the genome of the parasitiform tick *Hyalomma asiaticum*. Colors correspond to individual ALGs, with solid color indicating significant hits ($p < 0.05$) and transparent color indicating insignificant hits.

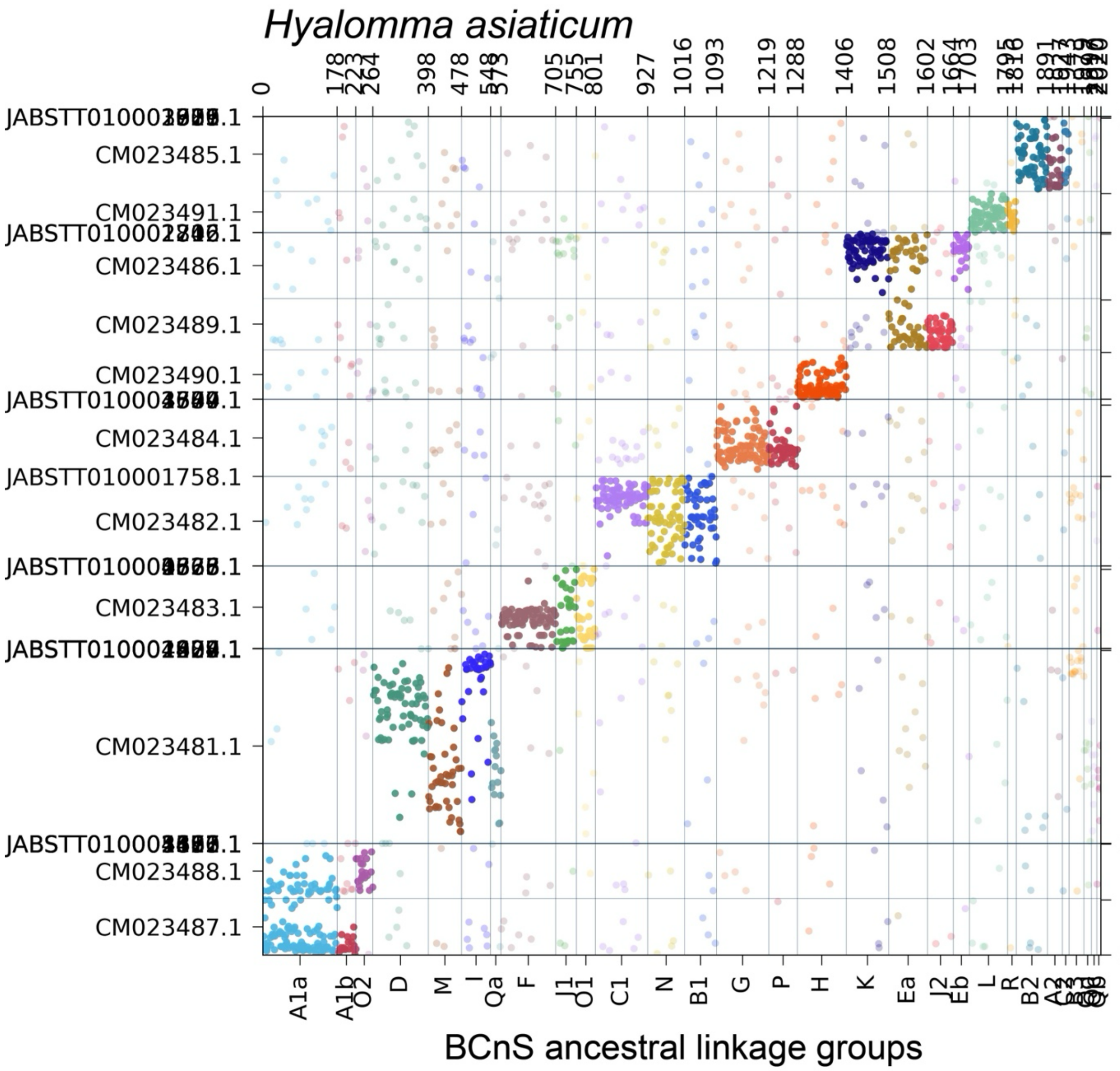

**Supplementary Figure S50.** Oxford dot plot showing the location of BCnS ancestral linkage group (ALG) homologs in the genome of the parasitiform tick *Ixodes scapularis*. Colors correspond to individual ALGs, with solid color indicating significant hits ($p < 0.05$) and transparent color indicating insignificant hits.

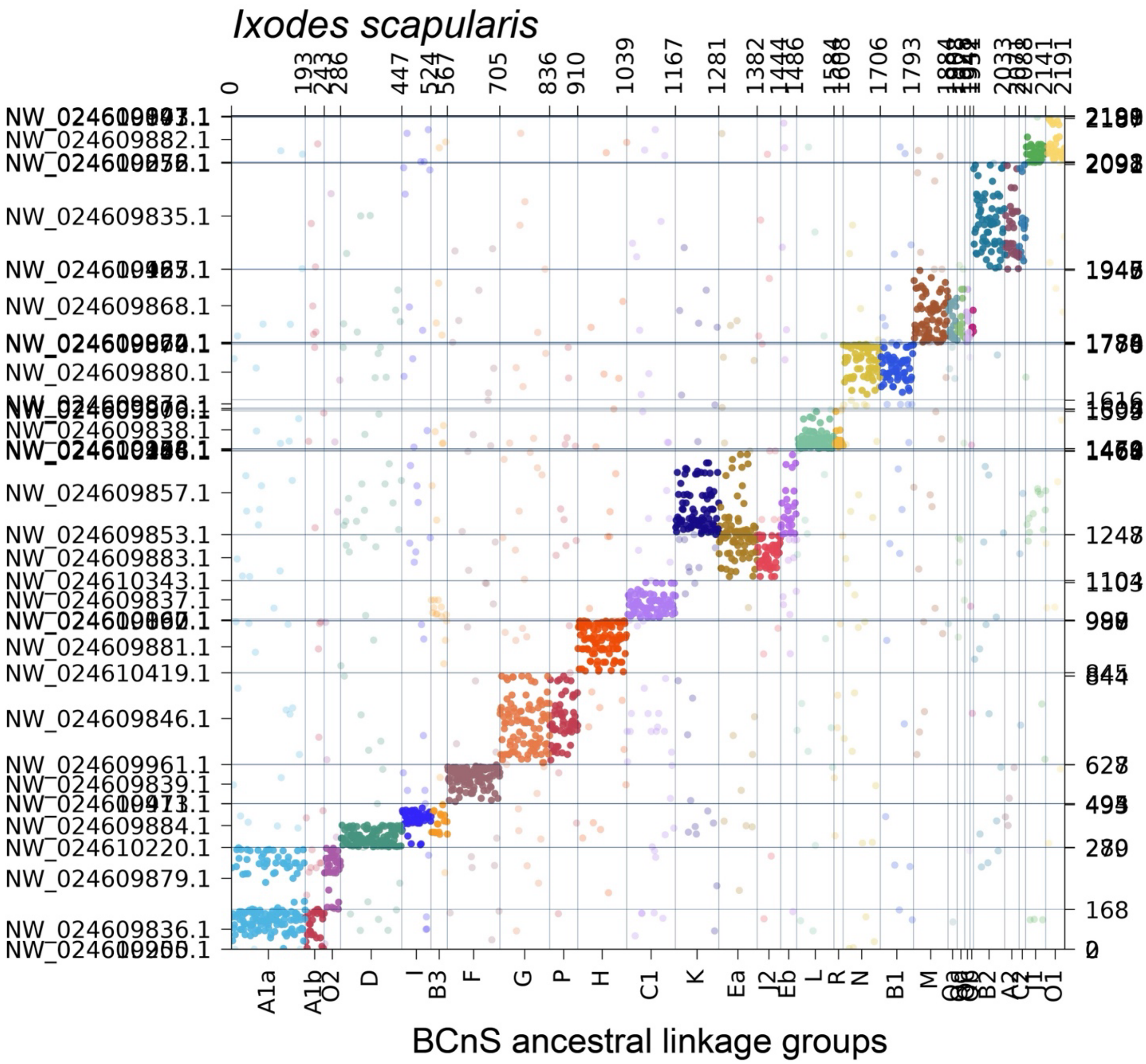

**Supplementary Figure S51.** Oxford dot plot showing the location of BCnS ancestral linkage group (ALG) homologs in the genome of the harvestman *Leiobunum manubriatum*. Colors correspond to individual ALGs, with solid color indicating significant hits ($p < 0.05$) and transparent color indicating insignificant hits.

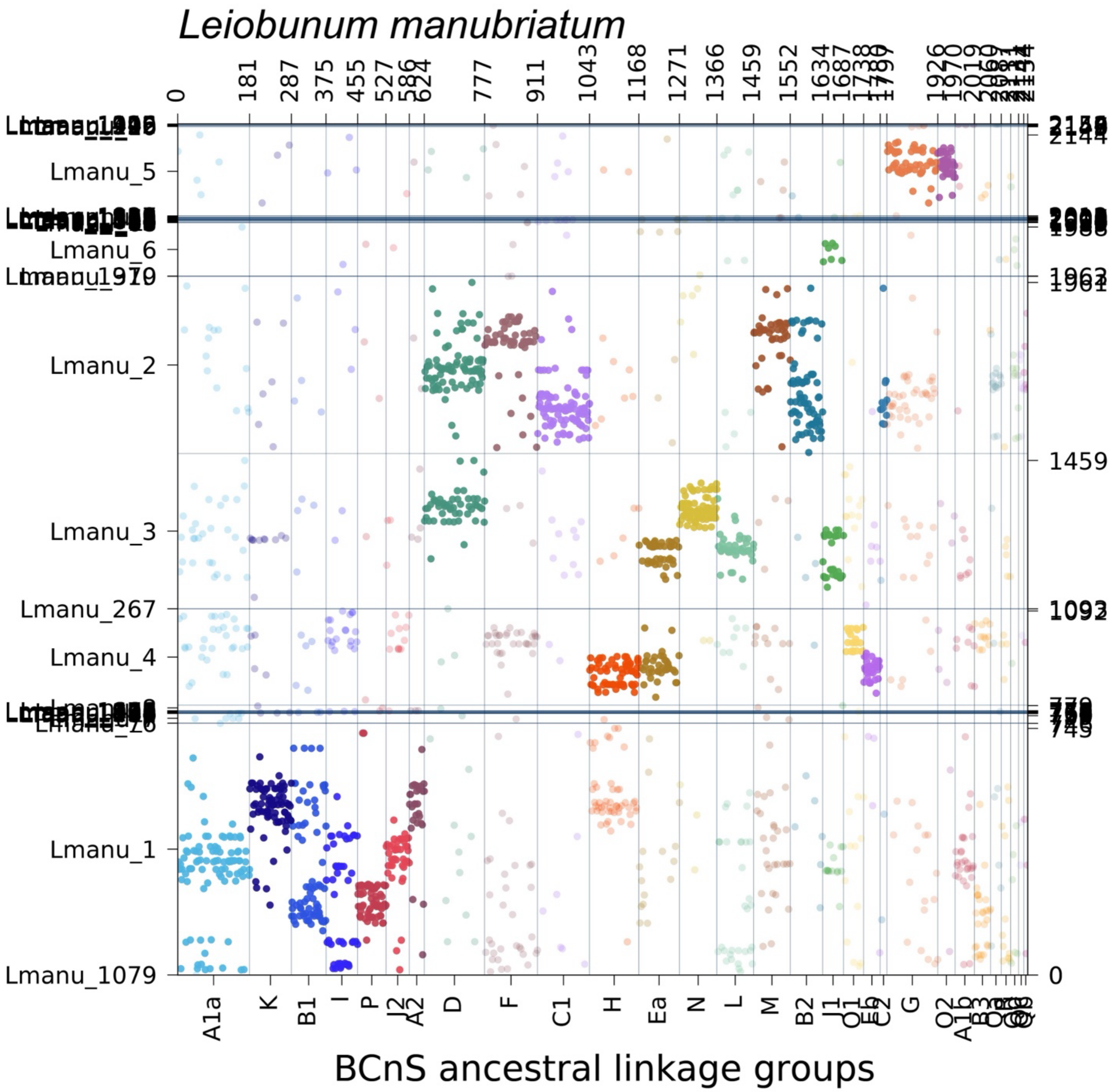

**Supplementary Figure S52.** Oxford dot plot showing the location of BCnS ancestral linkage group (ALG) homologs in the genome of the horseshoe crab *Limulus polyphemus*. Colors correspond to individual ALGs, with solid color indicating significant hits ($p < 0.05$) and transparent color indicating insignificant hits.

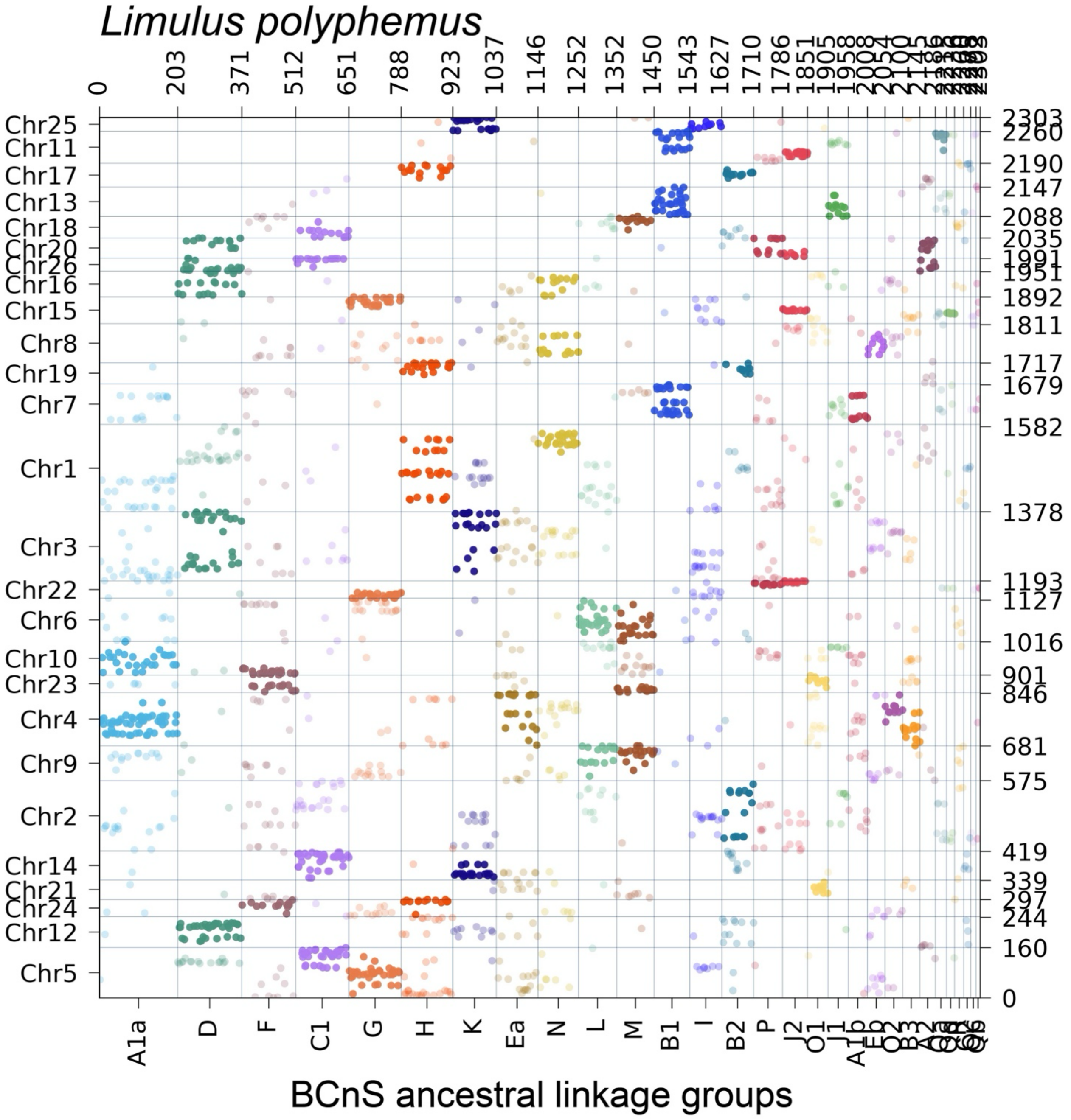

**Supplementary Figure S53.** Oxford dot plot showing the location of BCnS ancestral linkage group (ALG) homologs in the genome of the mesothele spider *Liphistius suwat*. Colors correspond to individual ALGs, with solid color indicating significant hits ($p < 0.05$) and transparent color indicating insignificant hits.

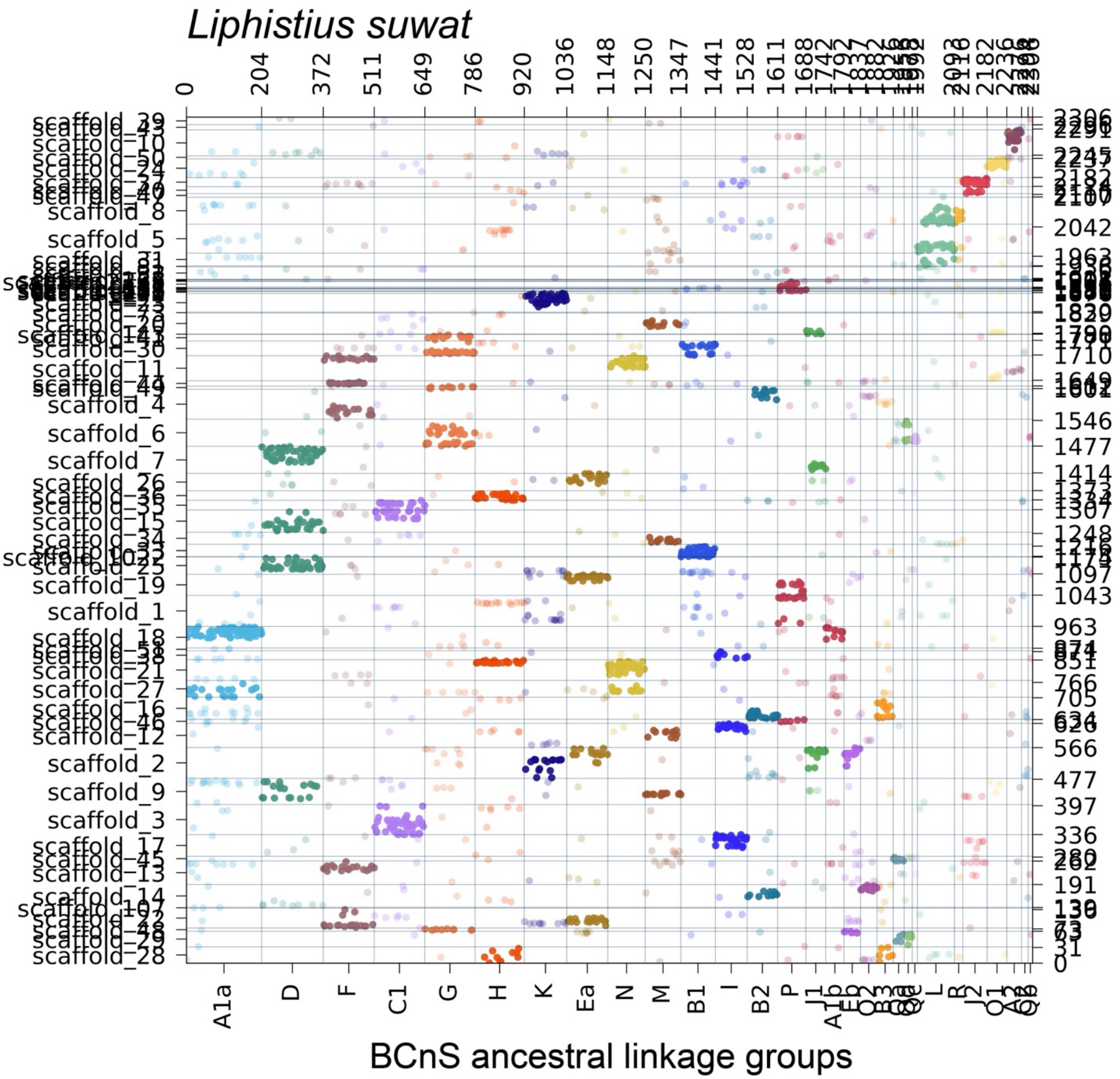

**Supplementary Figure S54.** Oxford dot plot showing the location of BCnS ancestral linkage group (ALG) homologs in the genome of the vinegaroon *Mastigoproctus giganteus*. Colors correspond to individual ALGs, with solid color indicating significant hits ($p < 0.05$) and transparent color indicating insignificant hits.

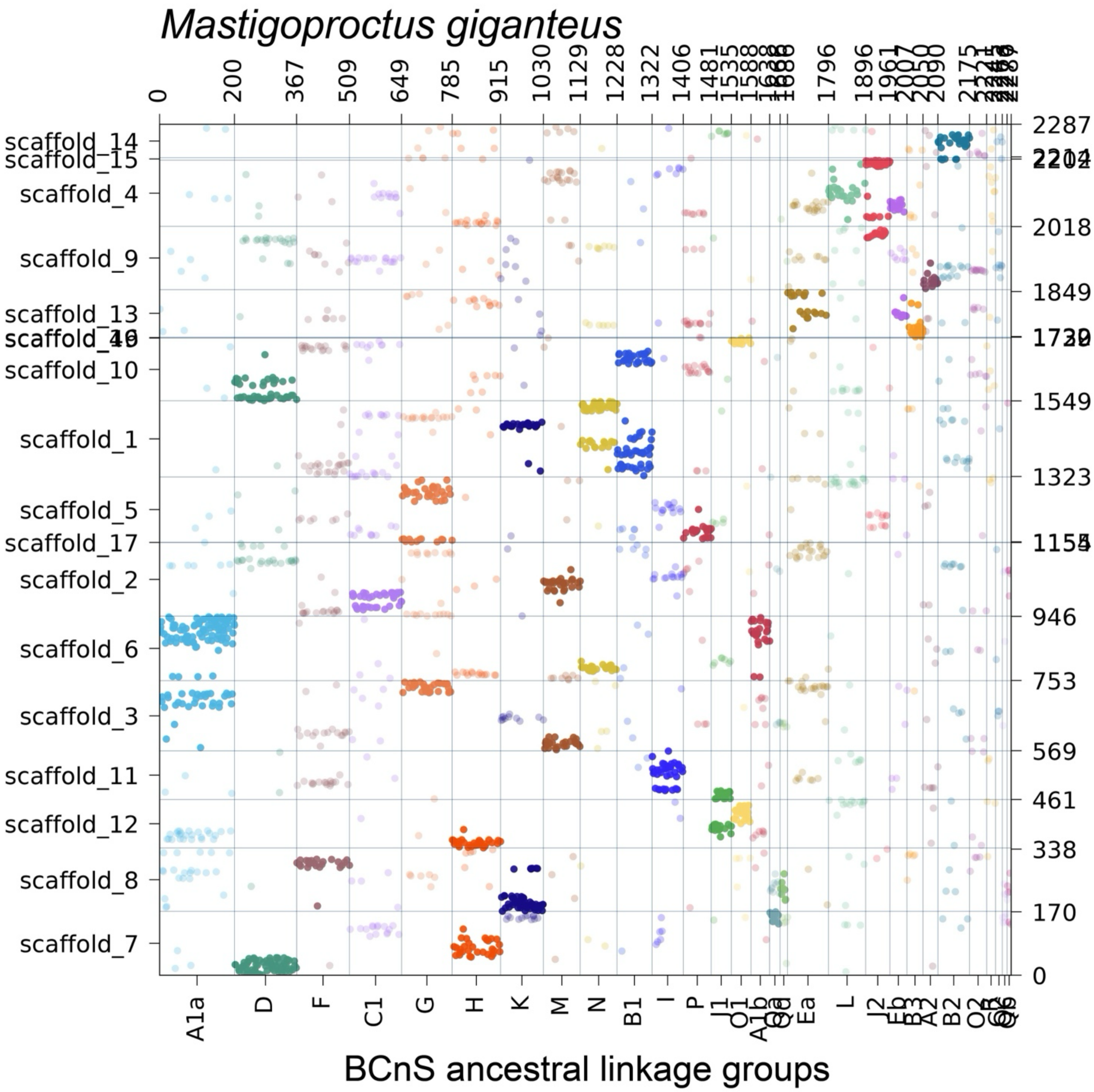

**Supplementary Figure S55.** Oxford dot plot showing the location of BCnS ancestral linkage group (ALG) homologs in the genome of the harvestman *Odiellus spinosus*. Colors correspond to individual ALGs, with solid color indicating significant hits ($p < 0.05$) and transparent color indicating insignificant hits.

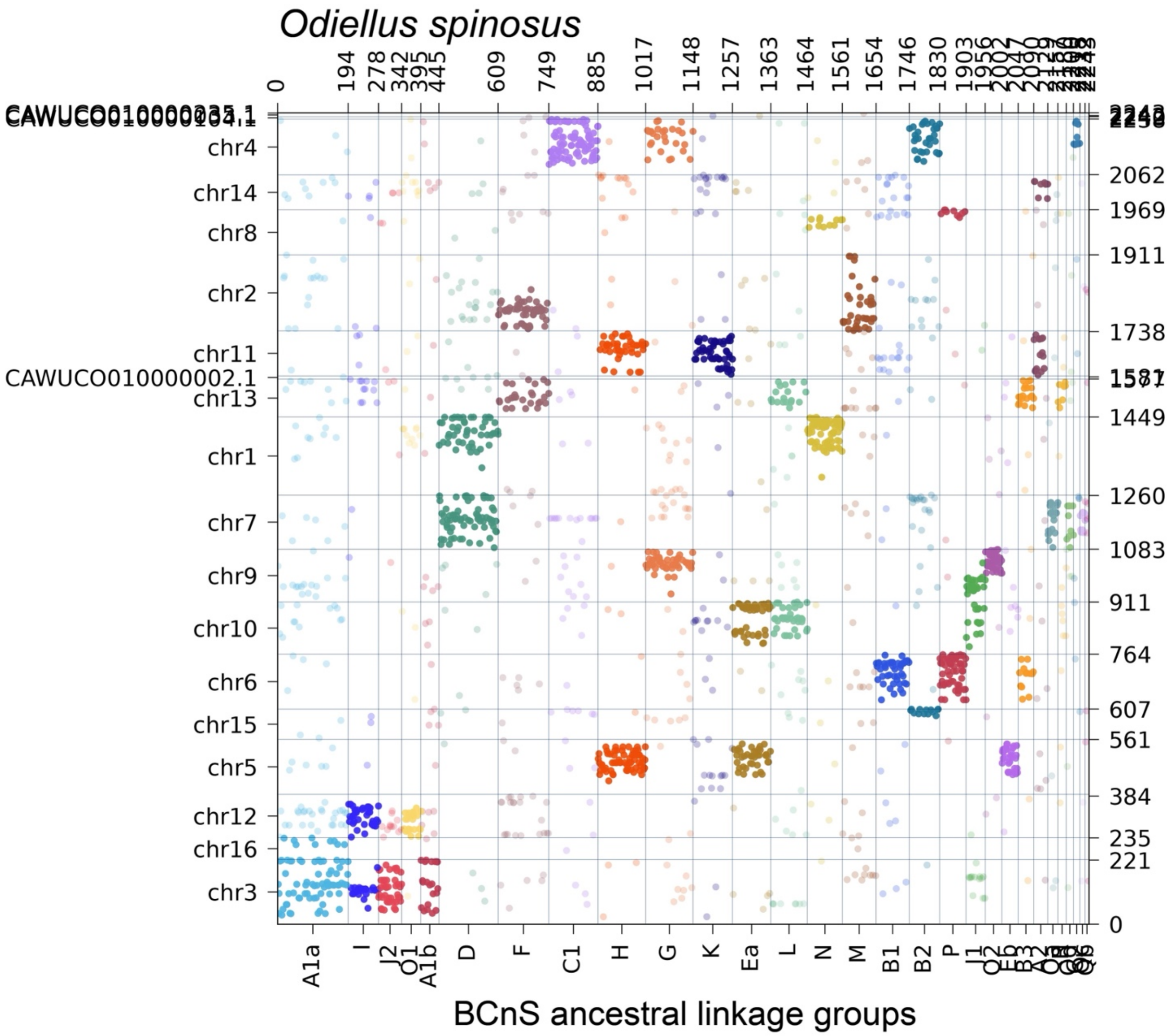

**Supplementary Figure S56.** Oxford dot plot showing the location of BCnS ancestral linkage group (ALG) homologs in the genome of the acariform mite *Oppia nitens*. Colors correspond to individual ALGs, with solid color indicating significant hits ($p < 0.05$) and transparent color indicating insignificant hits.

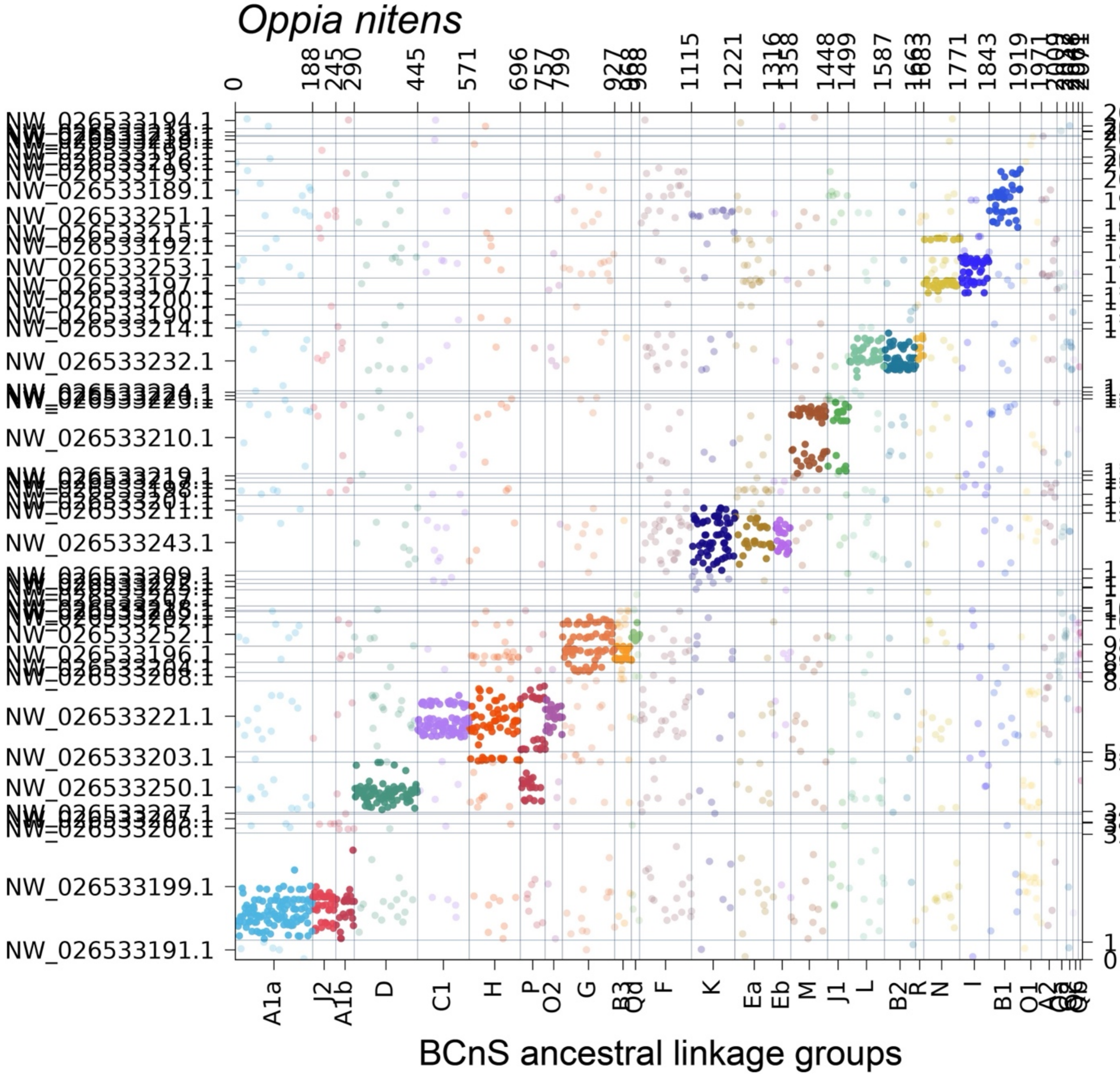

**Supplementary Figure S57.** Oxford dot plot showing the location of BCnS ancestral linkage group (ALG) homologs in the genome of the sea spider *Pycnogonum litorale*. Colors correspond to individual ALGs, with solid color indicating significant hits ($p < 0.05$) and transparent color indicating insignificant hits.

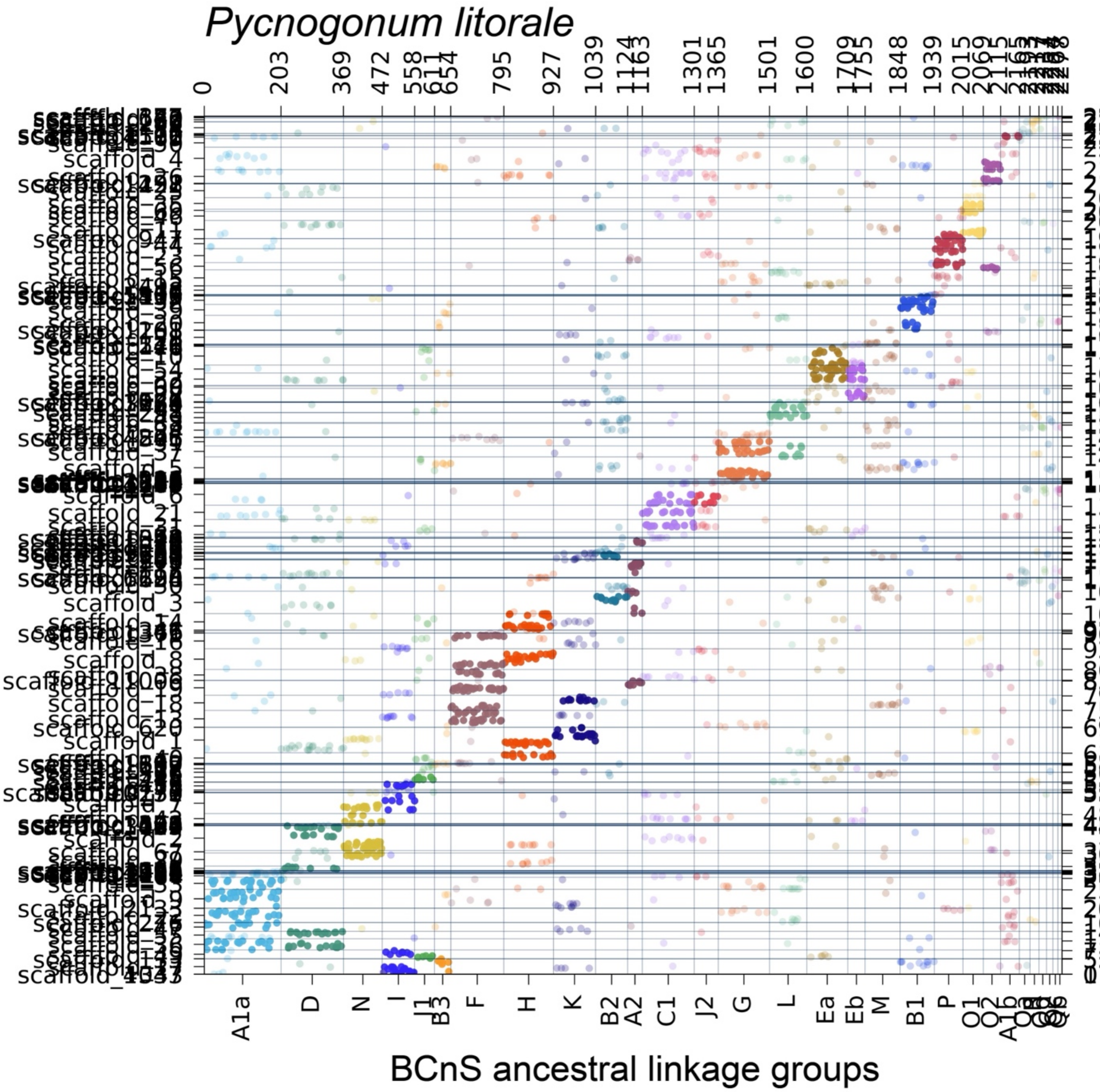

**Supplementary Figure S58.** Oxford dot plot showing the location of BCnS ancestral linkage group (ALG) homologs in the genome of the parasitiform tick *Rhipicephalus microplus*. Colors correspond to individual ALGs, with solid color indicating significant hits ($p < 0.05$) and transparent color indicating insignificant hits.

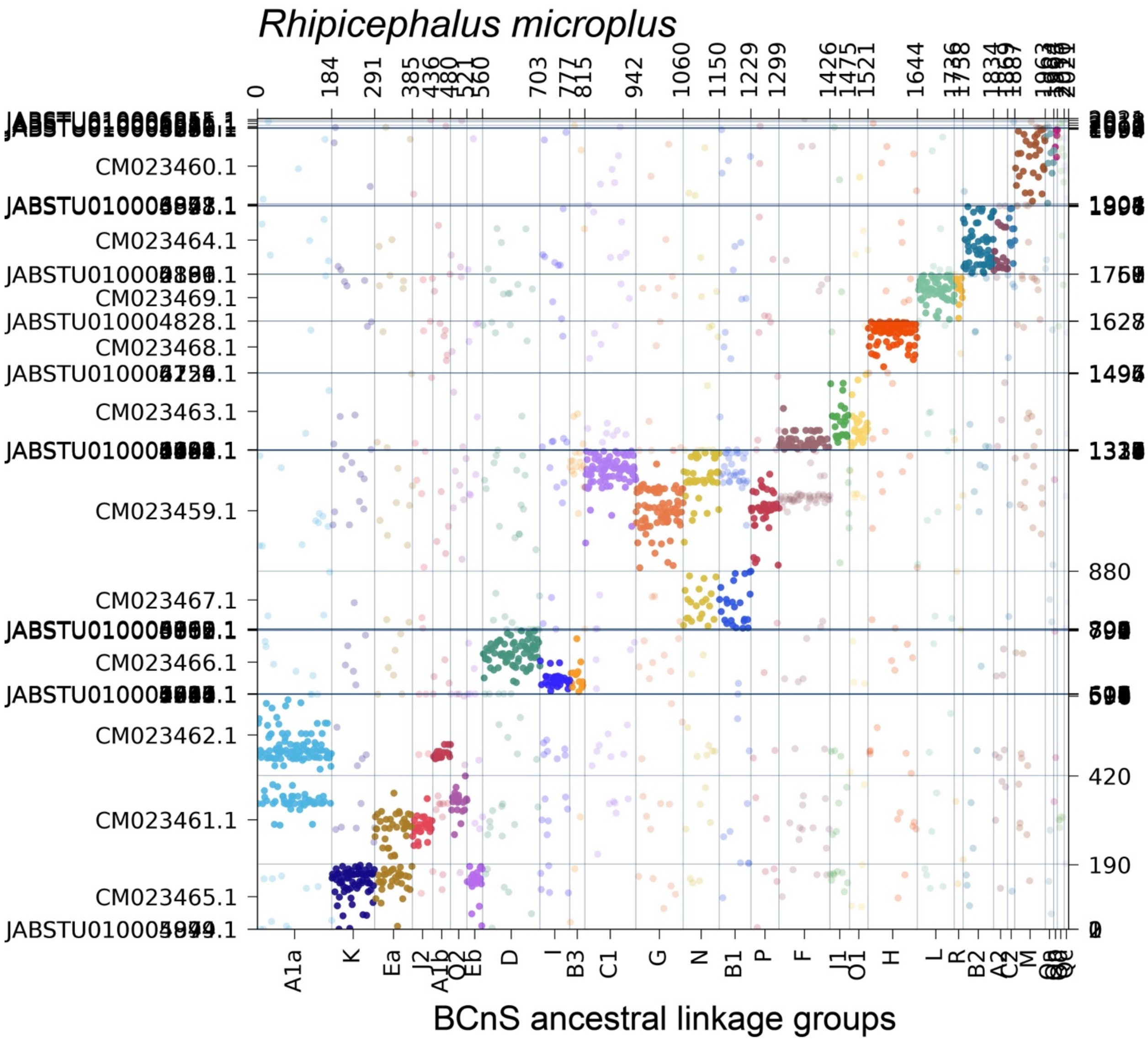

**Supplementary Figure S59.** Oxford dot plot showing the location of BCnS ancestral linkage group (ALG) homologs in the genome of the acariform mite *Sarcoptes scabiei*. Colors correspond to individual ALGs, with solid color indicating significant hits ($p < 0.05$) and transparent color indicating insignificant hits.

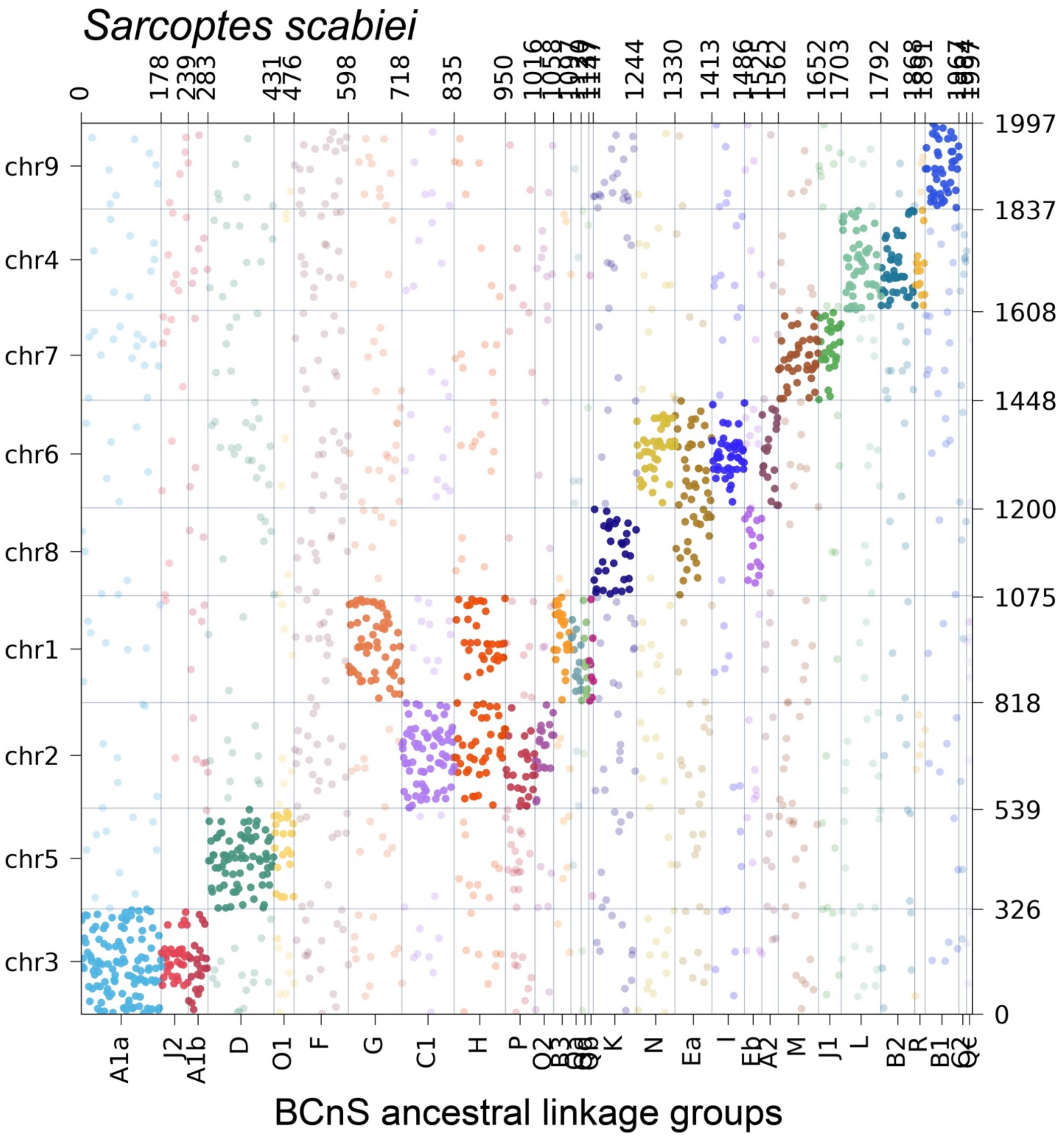

**Supplementary Figure S60.** Oxford dot plot showing the location of BCnS ancestral linkage group (ALG) homologs in the genome of the centipede *Strigamia acuminata*. Colors correspond to individual ALGs, with solid color indicating significant hits ($p < 0.05$) and transparent color indicating insignificant hits.

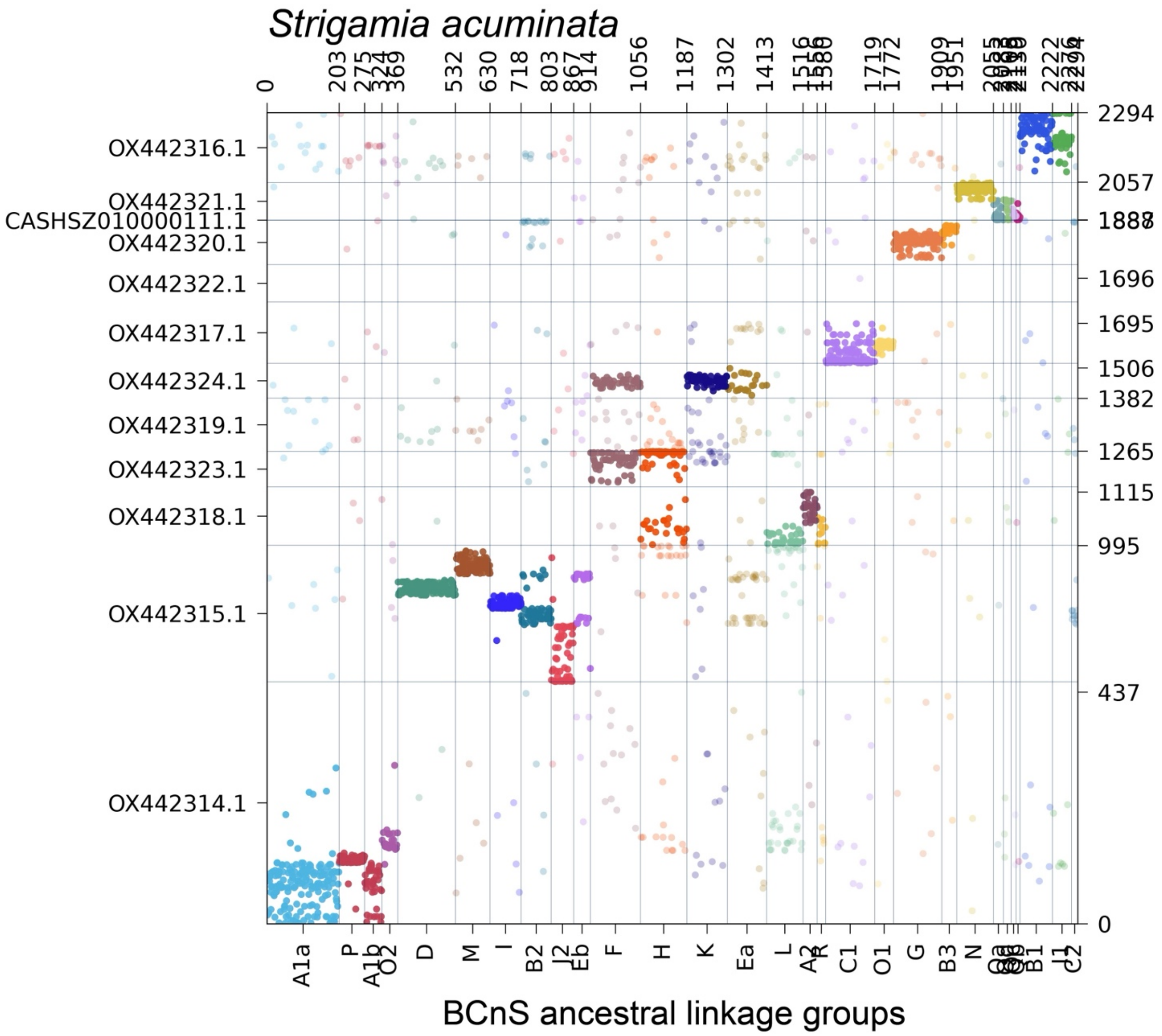

**Supplementary Figure S61.** Oxford dot plot showing the location of BCnS ancestral linkage group (ALG) homologs in the genome of the horseshoe crab *Tachypleus tridentatus*. Colors correspond to individual ALGs, with solid color indicating significant hits ($p < 0.05$) and transparent color indicating insignificant hits.

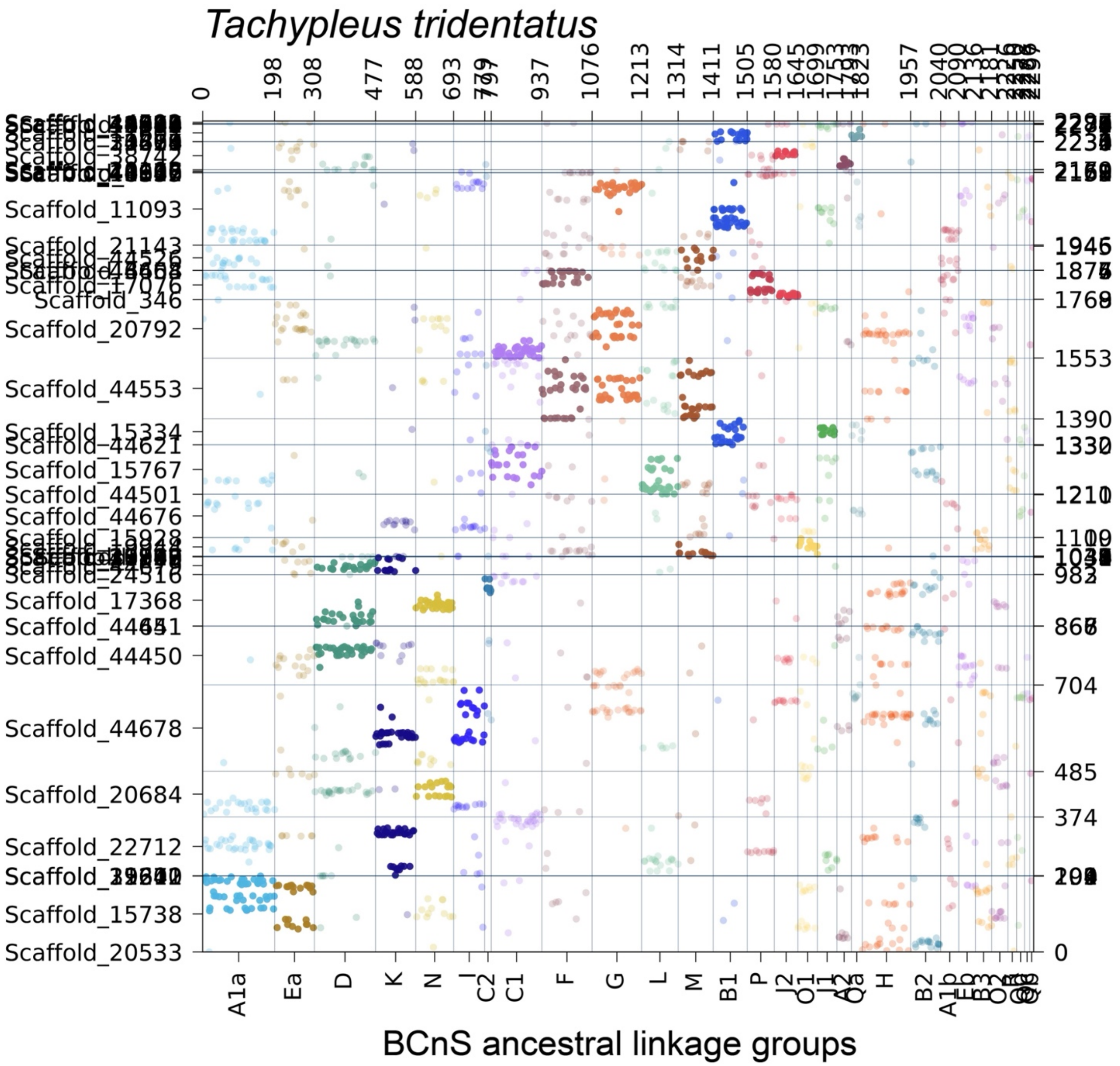

**Supplementary Figure S62.** Oxford dot plot showing the location of apulmonate ancestral linkage group (ALG) homologs in the genome of the parasitiform mite *Varroa destructor*. Colors correspond to individual ALGs, with solid color indicating significant hits ($p < 0.05$) and transparent color indicating insignificant hits.

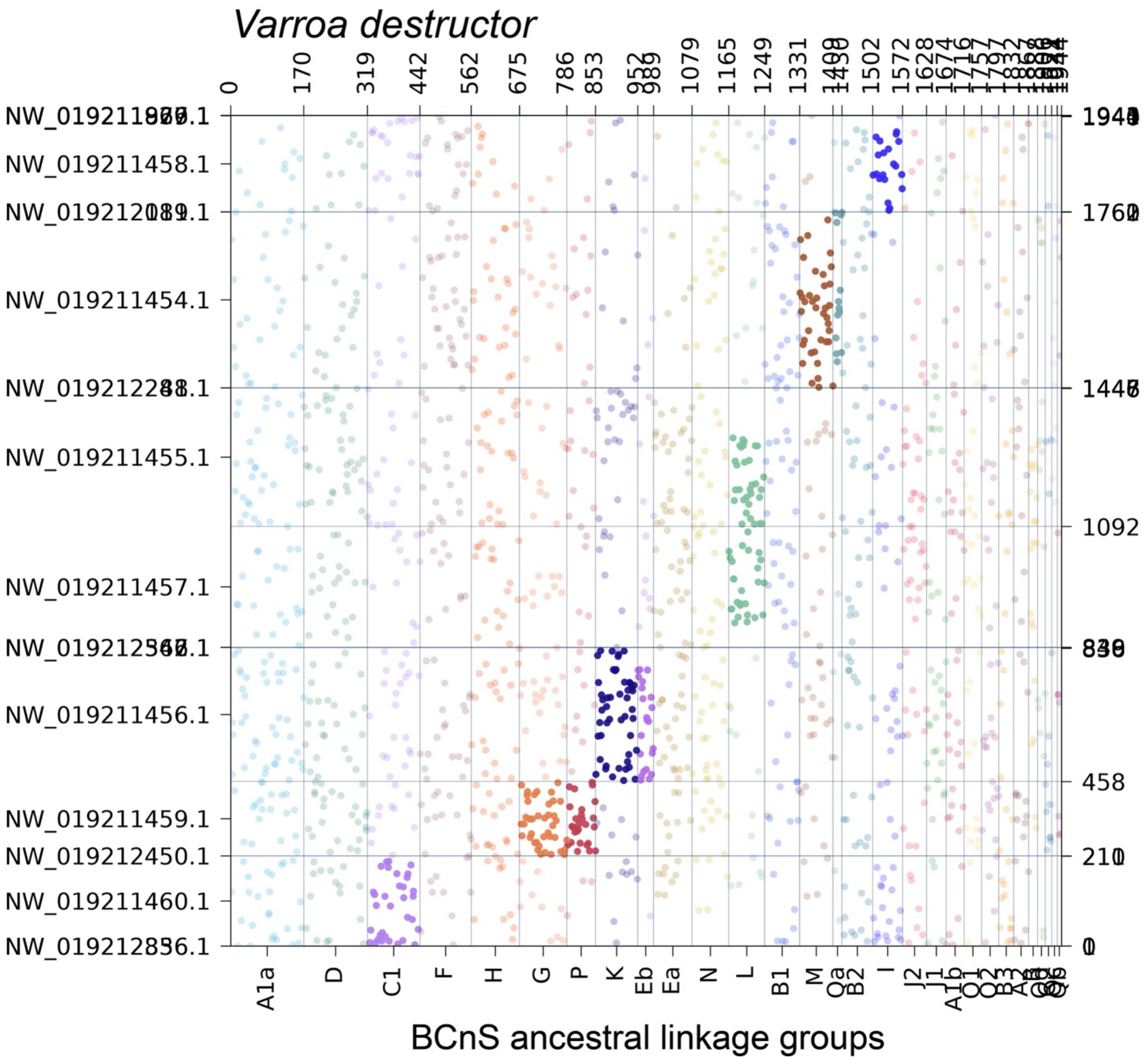

**Supplementary Figure S63.** Mapping of ortholog gains and losses on unconstrained tree topology *versus* topology constrained to recover alternative placements for Acariformes.

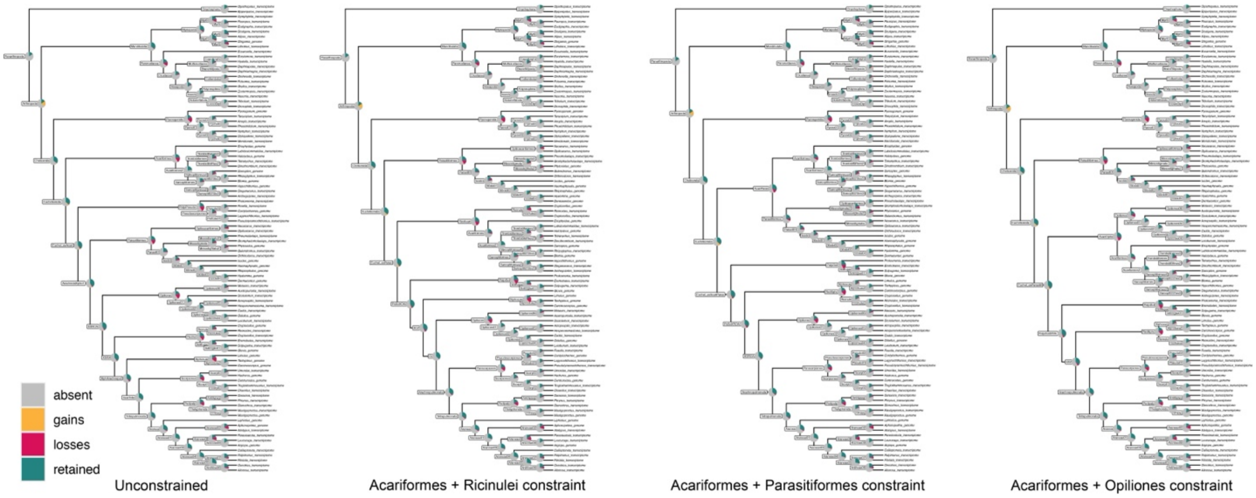

**Supplementary Figure S64.** Mapping of ortholog gains and losses on topology constrained to recover Arachnida *versus* topology constrained to recover Xiphosura + Arachnopulmonata.

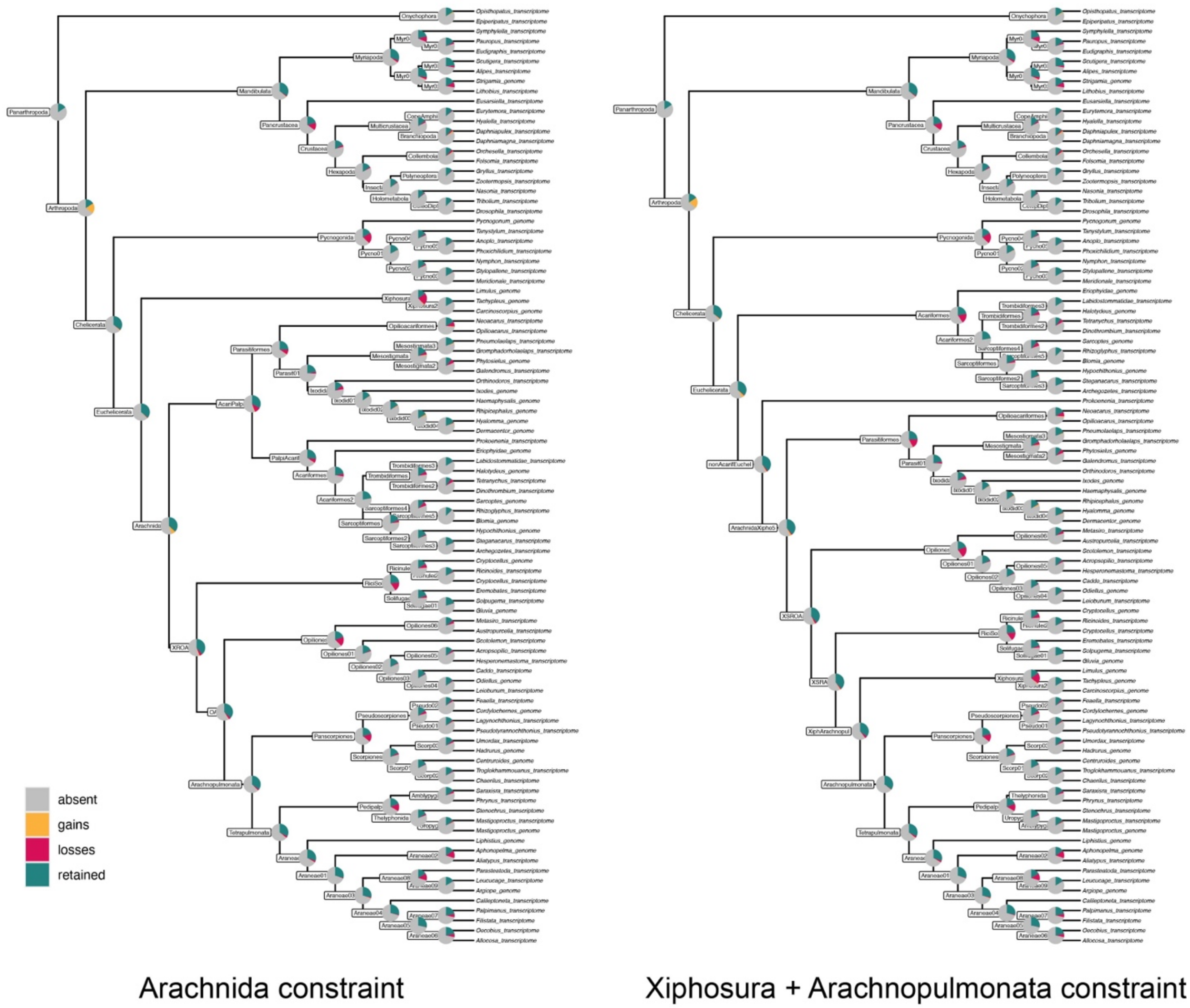

**Supplementary Figure S65.** Oxford dot plot showing the location of ancestral linkage group (ALG) homologs in the genomes of three mites for the apulmonate (top row) and the BCnS (bottom row) linkage groups. Left column: *Phytoseiulus persimilis* (Parasitiformes). Middle: *Tetranychus urticae* (Acariformes). Right: Eriophyidae gen. sp. (Acariformes). Colors correspond to individual ALGs, with solid color indicating significant hits ($p < 0.05$) and transparent color indicating insignificant hits.

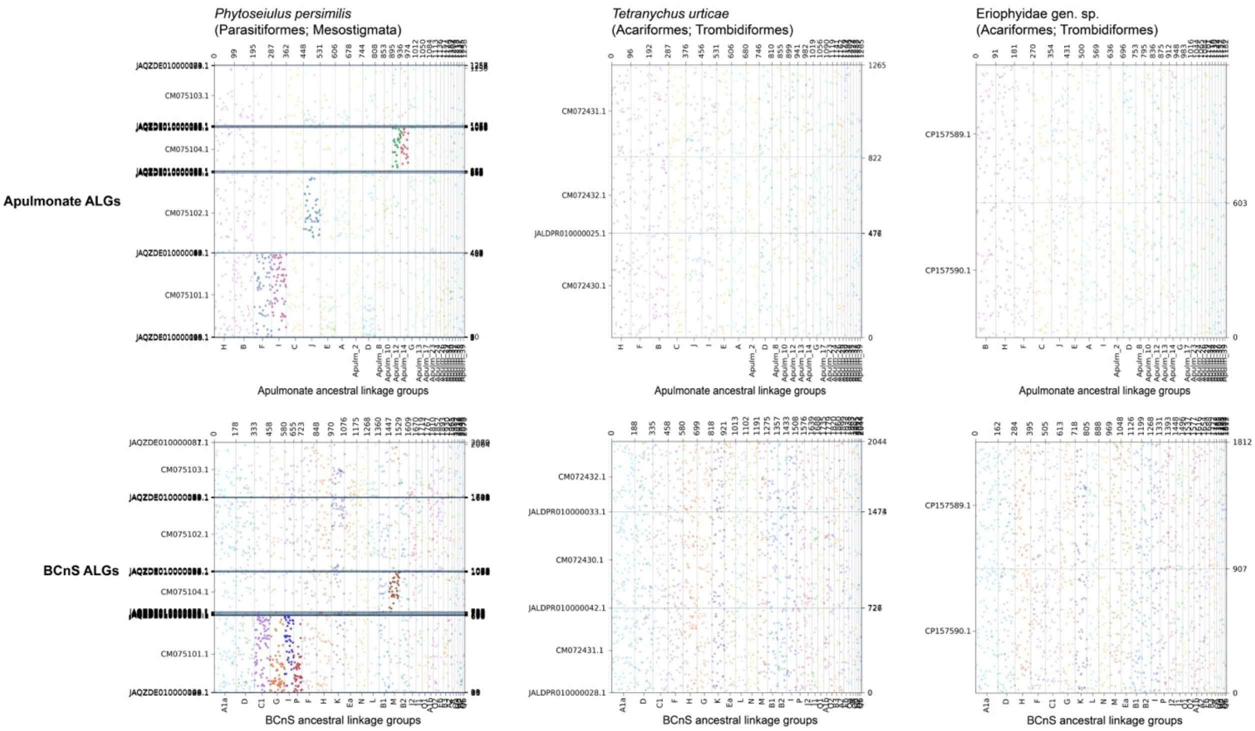
